# Menadione reduces the expression of virulence- and colonisation-associated genes in *Helicobacter pylori*

**DOI:** 10.1101/2024.11.07.622476

**Authors:** Stephen Thompson, Rebecca Omoyemi Ojo, Lesley Hoyles, Jody Winter

## Abstract

Novel treatment options are needed for the gastric pathogen *Helicobacter pylori* due to its increasing antibiotic resistance. The vitamin K analogue menadione has been extensively studied due to interest in its antibacterial and anti-cancer properties. Here, we investigated the effects of menadione on *H. pylori* growth, viability, antibiotic resistance, motility and gene expression using clinical isolates. The minimum inhibitory concentration (MIC) of menadione was 313 µM for 11/13 isolates and 156 µM for 2/13 isolates. The minimum bactericidal concentrations were 1.25-2.5 mM, indicating that concentrations in the micromolar range were bacteriostatic rather than bactericidal. We were not able to experimentally evolve resistance to menadione *in vitro*. Sub-MIC menadione (16 µM for 24 h) did not significantly inhibit bacterial growth but significantly (p < 0.05) changed the expression of 1291/1615 (79.9 %) genes encoded by strain 322A. Expression of the virulence factor genes *cagA* and *vacA* was downregulated in the presence of sub-MIC menadione, while genes involved in stress responses were upregulated. Sub-MIC menadione significantly (p < 0.0001) inhibited the motility of *H. pylori*, consistent with the predicted effects of the observed significant (p < 0.05) downregulation of *cheY*, upregulation of *rpoN*, and changes in expression of flagellar assembly pathway genes seen in the transcriptomic analysis. Through in-depth interrogation of transcriptomics data, we concluded that sub-MIC menadione elicits a general stress response in *H. pylori* with survival in the stationary phase likely mediated by upregulation of *surE* and *rpoN*. Sub-MIC menadione caused some modest increases in *H. pylori* susceptibility to antibiotics, but the effect was variable with strain and antibiotic type and did not reach statistical significance. Menadione (78 µM) was minimally cytotoxic to human gastric adenocarcinoma (AGS) cells after 4 hours but caused significant loss of cell viability after 24 hours.

Given its inhibitory effects on bacterial growth, motility and expression of virulence and colonisation-associated genes, menadione at low micromolar concentrations may have potential utility as a virulence-attenuating agent against *H. pylori*.

**Data summary:** RNAseq data reported on in this study are available to download from ArrayExpress under accession E-MTAB-14439. Supplementary materials associated with this publication are available from https://figshare.com/projects/Menadione_reduces_the_expression_of_virulence-_and_colonisation-associated_genes_in_Helicobacter_pylori/219631.

## Introduction

*Helicobacter pylori* is a Gram-negative, microaerophilic bacterium that persists in the human stomach and can cause ulcers and gastric cancer. *H. pylori* is exquisitely adapted to colonisation of, and survival in, the human stomach, producing a wide range of virulence- and colonisation-associated factors including adhesins, urease enzyme, and toxins. Strains expressing the oncogenic cytotoxin CagA, and more virulent forms of the vacuolating cytotoxin VacA, are more strongly associated with ulcers and gastric cancer(1–3). *H. pylori* is becoming increasingly difficult to treat due to antibiotic resistance(4), and new treatment options are needed.

Menadione (2-methyl-1,4-naphthoquinone, also known as vitamin K3) is a synthetic form of vitamin K that is used as a supplement in animal feed(5). It is an organic compound that displays oxidant activity by generating reactive oxygen species via redox cycling. Due to its cytotoxic effects against many different types of cancer cell lines, researchers have extensively studied the potential utility of menadione as an anti-cancer drug(6–10).

Menadione has been investigated for its anti-bacterial properties and shown to inhibit the growth of *Staphylococcus aureus*, *Bacillus anthracis* and *Streptococcus* spp.(11). It has also been reported to have antibiotic susceptibility-modifying activity, lowering the minimum inhibitory concentration (MIC) of antibiotics against multidrug-resistant strains of *Staphylococcus aureus*, *Escherichia coli* and *Pseudomonas aeruginosa*(12).

For *H. pylori*, menadione has been shown to inhibit bacterial growth using disk diffusion assays(13) and using agar plates supplemented with various concentrations of menadione(14). Lee et al (14) used RT-PCR and cellular assays to show that menadione reduced the expression of *vacA,* the *secA* gene involved in export of VacA, and the type IV secretion system components *vir*B2, *vir*B7 and *vir*B10 involved in secretion of CagA. They also used co-culture assays to show that pretreatment with menadione decreased the cytotoxic and pro-inflammatory effects of *H. pylori* on gastric adenocarcinoma (AGS) and monocytic leukaemia (THP-1) cells. Several studies have used RNA sequencing (RNASeq) to analyse the transcriptional responses of *H. pylori* to stressors including pH(15) and sodium chloride(16), but to our knowledge there has not yet been an RNASeq analysis of the effects of menadione on *H. pylori* gene expression.

In this study, we determined the minimum inhibitory concentration (MIC) of menadione for a collection of *H. pylori* clinical isolates and we examined the effects of sub-MIC menadione on *H. pylori.* We present the first in-depth transcriptomic study of the effects of a sub-inhibitory concentration of menadione on the expression of virulence- and colonisation-associated genes in *H. pylori.* We show that menadione can significantly downregulate the expression of numerous genes involved in colonisation, virulence, motility and epithelial cell interactions including *cagA, vacA, luxS, ureA/B/I* and *ruvC*. We also show that sub-MIC concentrations of menadione inhibit motility and may increase the susceptibility of some strains of *H. pylori* to antibiotics used in eradication therapy. Our findings indicate that menadione has potential utility as a virulence-attenuating agent against *H. pylori*.

## Methods

### Culture of *H. pylori*

*H. pylori* clinical isolates were kindly provided by John Atherton and Karen Robinson, University of Nottingham, having been isolated from gastric biopsies donated by patients attending Queens Medical Centre, Nottingham for upper gastrointestinal tract endoscopy. Written informed consent was obtained, and the study was approved by the Nottingham Research Ethics Committee 2 (08/H0408/195). Cultures were maintained on blood agar base #2 (Oxoid) supplemented with 5 % (v/v) defibrinated horse blood (Thermo Scientific) and incubated under microaerobic conditions (5 % O_2_, 10 % CO_2_, 85 % N_2_; Don Whitley DG250 cabinet) at 37 °C.

### Minimum inhibitory concentration (MIC) assay and growth curves

Forty-eight-hour *H. pylori* cultures on blood agar plates were suspended in Brucella broth (Sigma Aldrich) supplemented with 5 % heat-inactivated foetal calf serum (FCS; Sigma Aldrich) at an OD_600_ of 0.05, then menadione (Sigma Aldrich) was added to a final concentration of 0-10 mM, final volume 100 µl/well in triplicate in 96- well plates. Growth controls (bacteria in broth, no menadione) and sterility controls (broth only) were included in every 96-well plate. Plates were incubated under microaerobic conditions at 37 °C and bacterial growth was monitored by measuring the OD_600_ at 0, 24, 48, 72 and 96 h.

### Motility assay

Forty-eight-hour *H. pylori* cultures on blood agar plates were suspended in Brucella broth supplemented with 5 % FCS at an OD_600_ of 1.0, and 10 µl of the bacterial suspension was stabbed into the centre of soft agar plates (Brucella broth, 0.4 % agar, 5 % FCS) prepared with and without the addition of 8 and 16 µM menadione. After 7 days incubation under microaerobic conditions at 37 °C, bacterial motility was determined by measuring the diameter of the zone of growth through each plate. This experiment was conducted in triplicate technical replicates, independently repeated three times.

### Attempted experimental evolution of menadione resistance

*H. pylori* strain 322A was inoculated into MIC assays using the method described above, but with 16 technical replicates at each menadione concentration. After 24 and 48 h, plates were screened looking for evidence of bacterial growth in any wells containing menadione at or above 313 µM. This experiment was repeated independently seven times.

### RNA sequencing

*H. pylori* strain 322A was grown in 400 ml Brucella broth + 5 % FCS to an OD_600_ of 0.4 and this starter culture was checked for purity by Gram staining and streaking out on blood agar plates. The starter culture was divided into 10 x 15 ml cultures in T25 tissue culture flasks, then menadione was added to five of the flasks to a final concentration of 16 µM. All flasks were incubated at 37 °C under microaerobic conditions for 24 h. Further purity checks were then carried out and the CFU/ml in each flask was determined by Miles & Misra(17). Bacteria were harvested by centrifugation and the cell pellets were submerged in TRIzol (Invitrogen) and shipped on dry ice to Macrogen (South Korea) for RNA extraction, library preparation and sequencing. Samples were relabelled in a randomised order before sending to the sequencing provider, to avoid any potential for introduction of a technical batch effect based on which group (treated or untreated) the samples belonged to.

Total RNA extraction was conducted by Macrogen and quality checked to confirm that all 10 samples were of acceptable RNA Integrity Number (RIN >7). Libraries were constructed using the Illumina TruSeq Stranded Total RNA (Illumina Inc., San Diego) protocol with ribosomal RNA depletion (Ribo-Zero). For mRNA sequencing, the barcoded libraries were loaded into the flow cell of an Illumina NovaSeq 6000 to generate 40M 100-bp paired-end reads per sample. Post sequencing, the raw data were converted into fastq files using the Illumina package bcl2fastq by Macrogen, and the fastq files were returned to the research team for analysis.

### Identification of differentially expressed genes

The US web-based platform Galaxy (http://usegalaxy.org/)(18) was used in initial analyses of transcriptomic data. The genome sequence (fasta file) and annotations (gff file) for *H. pylori* 322A (reported in *Wilkinson et al*, 2022 (19) and available from doi:10.6084/m9.figshare.26931724) were uploaded to Galaxy along with the fastq output files for each sample from the RNASeq run. Quality of the sequence reads was assessed using FastQC v0.11.9 (20) within Galaxy; no trimming of reads was needed as all sequence data were of high quality (Phred >20) and free of adapter contamination. Sequence reads were then mapped to the 322A reference genome using HISAT2 v2.2.1 set to paired-end library (21) before counting the number of reads that mapped to genes using featureCounts (Galaxy v2.0.1+galaxy2)(22). To test for differential gene expression between the menadione-treated and untreated samples DESeq2 v1.34.0 (23) used the count data from the featureCounts outputs and generated a normalised counts file and summary plots (available from doi:10.6084/m9.figshare.27256641) for the entire dataset. Genes were considered significantly differentially expressed based on adjusted P value <0.05 (Benjamini-Hochberg procedure).

### Visualisation of significantly differentially expressed genes

The log_2_-transformed normalised gene count data for the significantly differentially expressed genes identified by the DESeq2 analysis were visualised in a heatmap generated using the R package heatmap.2 from gplots v3.1.3. Boxplots for *cagA* and *vacA* gene expression data and a volcano plot for all DESeq2 data were generated using the R package tidyverse v2.0.0.

A KEGG-based network analysis was undertaken using the significantly differentially expressed genes. To do this, KEGGREST v1.40.0 was used to download information on the KEGG entry (hpy, release 111.0+/08-24, August 2024) for *H. pylori* strain 26695. Nucleotide sequences of the 1632 genes associated with strain 26695’s genome by KEGG were downloaded and used to create a BLASTN database (BLAST v2.12.0+) against which the 1615 predicted genes of strain 322A were searched. Hits were filtered based on highest query coverage and identity, with this information used to map 322A genes (and log_2_ fold change gene expression data) to the genome of strain 26695. Data were analysed using SPIA v2.52.0 (24) and also used to create a KEGG pathway-based network graph (KEGGgraph v1.60.0; (25)). igraph v1.5.1 (26)was used to generate network statistics, with RCy3 v2.20.2 (27) used to export the network and visualise it with Cytoscape v3.10.2 (28).

### Gene over-representation analysis

KEGG mapping data were available for 101 pathways in the reference strain, covering 634/1632 of *H. pylori* 26695’s genes (hpy, release 111.0+/08-24, August 2024). Almost two-fifths (632/1615; 39.1%) of strain 322A’s genes mapped to one or more KEGG pathways annotated in *H. pylori* 26695. The number of significantly (adjusted P value < 0.05, Benjamini-Hochberg) differentially expressed genes in 322A that mapped to genes in each of the 101 *H. pylori* 26695 pathways was determined, with a one-sided Fisher’s exact test used to determine the significance of gene over-representation. Generic pathways (i.e. hpy01100 – Metabolic pathways, hpy01110 – Biosynthesis of secondary metabolites and hpy01120 – Microbial metabolism in diverse environments) were excluded from analyses.

### Antibiotic susceptibility modifying effects

The effects of sub-MIC concentrations of menadione on antibiotic susceptibility of *H. pylori* were investigated using modified agar dilution and disc diffusion assays. For the agar dilution assays, a 10-fold dilution series of a bacterial suspension starting at OD_600_ of 3.0 +/- 0.2 in 0.85% saline was spotted onto blood agar plates containing 0.1, 0.5 or 2.5 µg/ml clarithromycin with and without 15.6 µM menadione. Spots (10 µl) of each bacterial dilution were applied to each plate in triplicate, and three different *H. pylori* clinical isolates were tested.

After incubation for 48 h, the density of bacterial growth in each spot was assessed visually and assigned a semi-quantitative score (0 = no growth, 1 = low density (individual colonies visible), 2 = moderate growth, 3 = dense growth/lawn). For the disc diffusion assays, *H. pylori* strains at OD_600_ of 0.1 were spread on Mueller Hinton agar (Thermo Scientific) supplemented with 7.5 % defibrinated horse blood and 0, 1 or 10 µM menadione. Discs containing 5 µg metronidazole, 15 µg clarithromycin or 5 µg levofloxacin were applied to the plates, in triplicate for each strain, antibiotic and experimental condition. Zone of inhibition diameters were measured after 3 days of incubation. The *H. pylori* strains selected for each disc diffusion assay had previously(4) been determined to be clinically resistant to the relevant antibiotics (n = 3 strains for metronidazole and clarithromycin; n = 2 strains for levofloxacin).

### Epithelial cell culture and cytotoxicity assays

Gastric adenocarcinoma (AGS) cells (ATCC CRL-1739) were maintained in Hams F12 medium (Sigma Aldrich) supplemented with 10 % (v/v) FCS at 37 °C in 10 % CO_2_. Cells were split, washed, seeded into 96-well opaque-walled, clear-bottom tissue culture treated plates (Greiner Bio-One) at 1 x 10^4^ cells per well, and treated with 0-5 mM menadione for 0-24 h. At each timepoint, cell viability was determined using a CellTiter Glo assay (Promega) according to the manufacturer’s instructions. Cytotoxicity assays were conducted in triplicate wells for each time point and menadione concentration.

## Results

### Menadione inhibits the growth of *H. pylori*

MICs were determined for 13 clinical isolates of *H. pylori* from seven patients (paired antrum and corpus isolates from six patients and an antrum isolate from a seventh patient). The MIC was 313 µM for 11/13 isolates (including strain 322A) and 156 µM for 2/13 isolates. Replating of 10 µl samples from each well in which *H. pylori* growth had been suppressed onto fresh blood agar plates yielded viable bacteria up to 1.25 mM (n = 2 strains) or 2.5 mM (n = 7 strains) menadione, indicating that menadione concentrations in the micromolar range were bacteriostatic rather than bactericidal. A complete MIC dataset for strain 322A is shown in Figure 1 and data for all tested strains is summarised in Supplementary Table 1.

**Figure 1.**
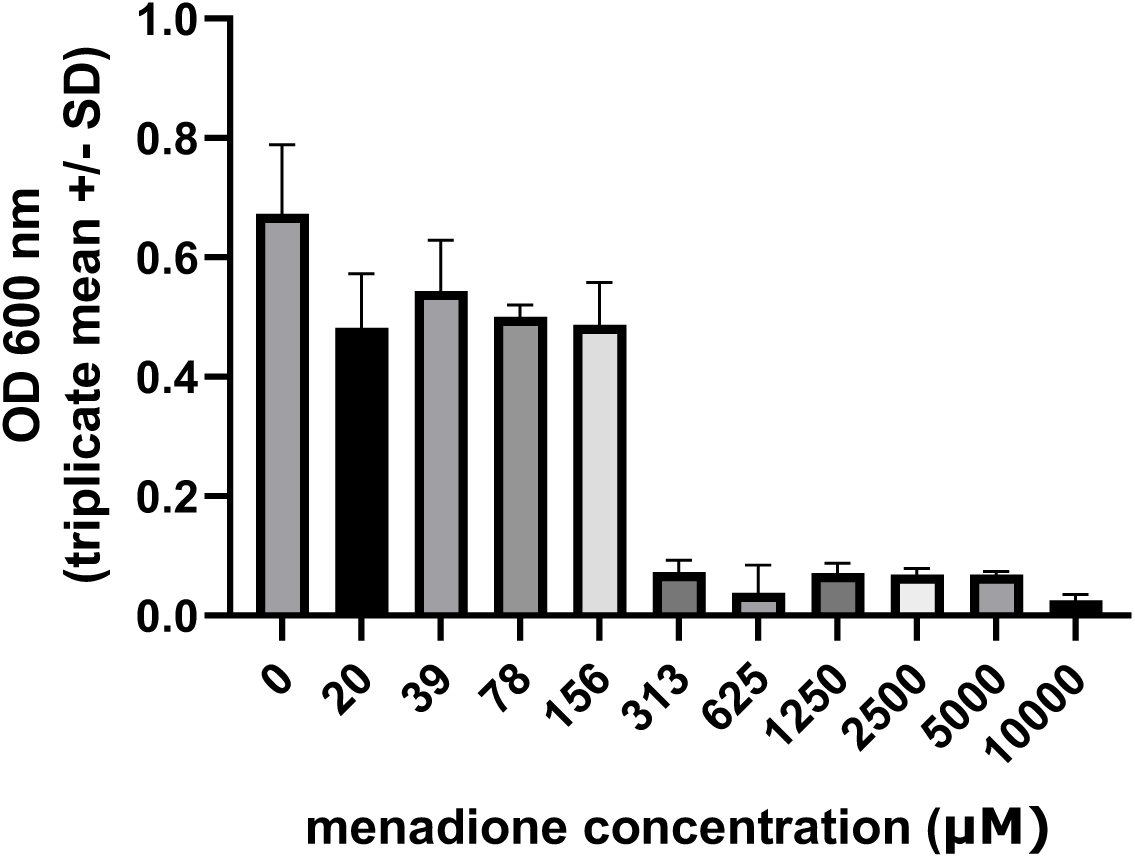
Inhibitory effects of menadione on the growth of *H. pylori* strain 322A. Bacteria were exposed to 0-10,000 µM menadione in liquid culture and growth was monitored by measuring OD_600_. Blank-corrected triplicate means +/- SD from the 72 h time point are shown. The MIC for this strain was 313 µM menadione. Summary MIC data from all strains tested are in Supplementary Table 1.

### Efforts to experimentally evolve resistance to menadione were not successful

Across many rounds of repeats and bacterial generations, broth microdilution-based experimental evolution assays using *H. pylori* strain 322A failed to yield any viable menadione-resistant mutants. There were no observable mutants that crossed the MIC threshold for menadione resistance after screening >100 replicates for growth at a range of menadione concentrations in the broth microdilution format (data not shown).

### Sub-MIC menadione has a profound effect on the transcriptome of strain 322A

Replicate flasks of *H. pylori* strain 322A were randomised into treated (n = 5) and untreated (n = 5) groups. Menadione was added to the treated group flasks at 16 µM and after 24 h the bacteria from all flasks were recovered for RNAseq. The mean CFU/ml was 1.5 x 10^8^ in the treated flasks and 3.0 x 10^8^ in the untreated flasks after 24 h, and monitoring of strain 322A growth over 24-96 h +/- 19.5 µM or 39 µM indicated that there was no significant inhibition of bacterial growth using these concentrations of menadione (Supplementary Figure 1) (p>0.05, 2-way ANOVA with Dunnett’s post-hoc test).

Treatment with 16 µM menadione significantly (adjusted P value < 0.05; Benjamini-Hochberg) changed the expression of 1291/1615 (79.9 %) genes encoded by strain 322A (Supplementary Table 2). Principal component analysis (doi:10.6084/m9.figshare.27257532) showed that the treated samples were very different from the untreated samples (88 % of variance explained by principal component 1), and the significantly differentially expressed genes clustered separately from one another based on treatment group (Figure 2a). The profound effect of menadione on strain 322A’s transcriptome was even more evident when data were visualised in a volcano plot, with the expression of 657 and 634 genes, respectively, significantly downregulated and upregulated by the compound (Figure 2b). Among the significantly differentially expressed genes were the virulence factors *cagA* and *vacA*, both of which were downregulated in the presence of menadione (Figure 2b, c).

**Figure 2.**
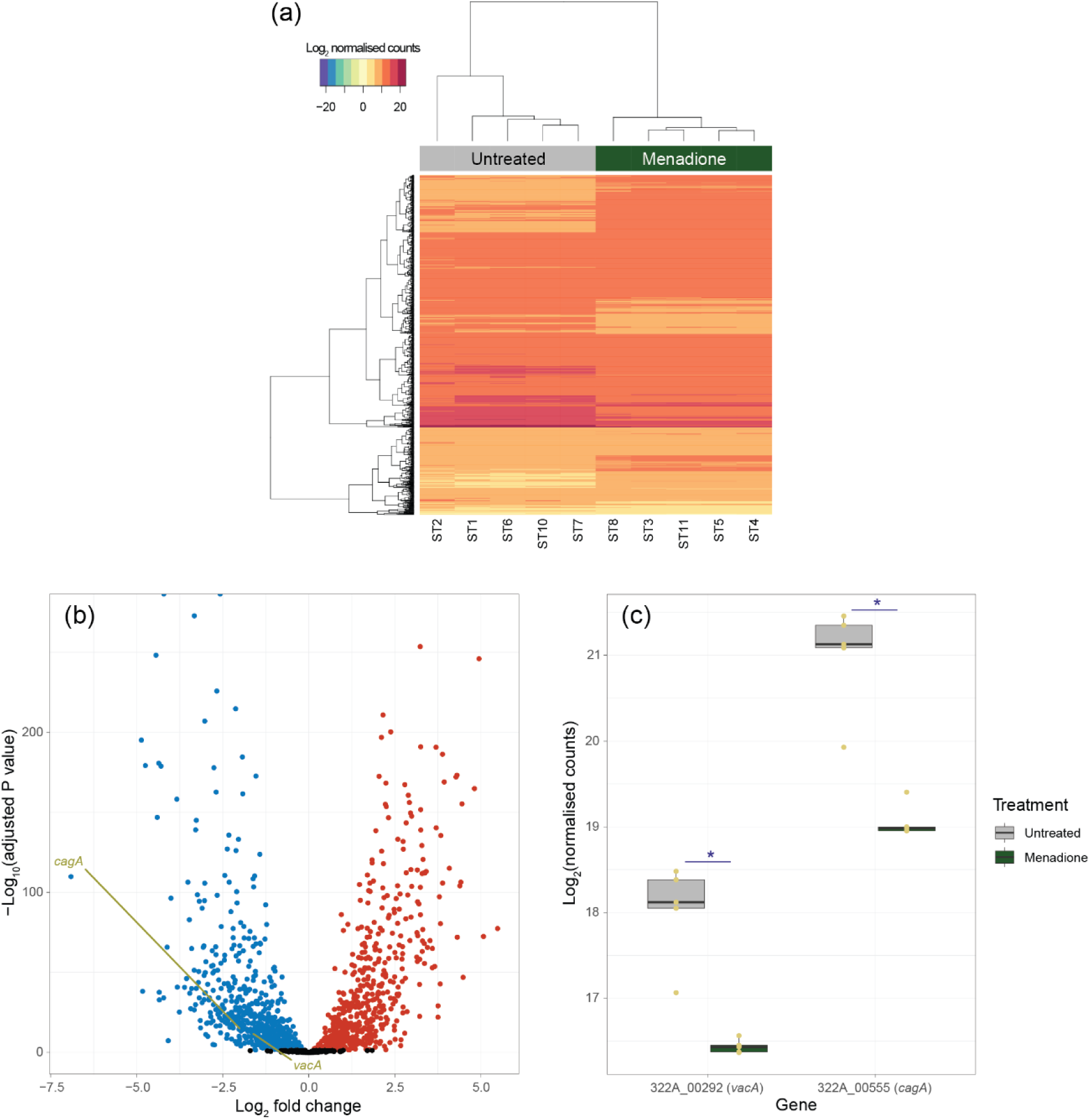
Visualisation of transcriptomic data. (a) Bidirectional heatmap, generated using log_2_-transformed normalised gene count data, showing clustering of the 1291 significantly (adjusted P value < 0.05; Benjamini-Hochberg) differentially expressed genes into distinct menadione-treated and untreated groups. (b) Volcano plot showing menadione treatment has a profound effect on gene expression in strain 322A. Significantly differentially expressed genes (adjusted P < 0.05; Benjamini- Hochberg) are shown by blue (downregulated by menadione) and red (upregulated by menadione) dots; non-significant data are shown by black dots. (c) Among the significantly downregulated genes were *cagA* and *vacA*. Boxplots show data points (yellow dots, n = 5 per group) and median values, with the hinges corresponding to the first and third quartiles; the whiskers extend from the hinges no further than +/- 1.5 * interquartile range. *Significantly differentially expressed in menadione-treated group compared with untreated control: adjusted P < 0.05 (Benjamini-Hochberg). (a–c) All data presented are from the same menadione treated (n = 5; 16 µM, 24 h) and untreated (n = 5) samples.

### Functional analyses of transcriptomic data

The genome of strain 322A is not included in the Kyoto Encylopedia of Genes and Genomes (KEGG), so to interrogate the effects of menadione on the biological functions of 322A we used *H. pylori* strain 26695 as a reference for KEGG-based analyses and used BLASTN to match genes from strain 322A (n = 1615) with those encoded by strain 26695 (n = 1632). The majority (1473/1615, 91.2 %) of strain 322A’s genes matched those in strain 26695. Among strain 26695’s genes, 980 (60 %) have KEGG orthology annotations and these all matched genes encoded in strain 322A’s genome (980/1615, 60.7 %).

Signalling Pathway Impact Analysis (SPIA), previously used to identify treatment effects in the transcriptomes of mammalian cell lines (29), was used to determine whether the approach had utility in analysis of microbial transcriptomic data. While non-significant, SPIA results implied menadione caused inactivation/downregulation of chemotaxis in strain 322A (Supplementary Table 3). Two-component systems, including the flagellar regulon, were also affected by menadione based on SPIA. Of note were upregulation of the gene (*dnaA*; 322A_00187) encoding the chromosomal replication initiator protein and that encoding RNA polymerase sigma-54 factor (*rpoN*; 322A_00890), known to have roles in stress responses (30). *rpoN* is linked with regulation of stationary phase, amino sugar metabolism, attaching and effacing lesions, flagellar assembly, acidic amino acid uptake and metabolism, biofilm formation, motility and virulence, and regulation of swarming activity.

SPIA also highlighted the expression of genes linked to redox signalling, and thereby energy production, was downregulated: electron transfer system – ubiquinol cytochrome *c* oxidoreductase, Rieske 2Fe-2S subunit (*fbcF*; 322A_00156) and ubiquinol cytochrome *c* oxidoreductase, cytochrome *b* subunit (*fbcH*; 322A_00157); aerobic respiration – cytochrome *c* oxidase, heme *b* and copper-binding subunit, membrane-bound (*fixN*; 322A_01308), cytochrome *c* oxidase, monoheme subunit, membrane-bound (*fixO*; 322A_01307), cbb3-type cytochrome *c* oxidase subunit Q (*ccoQ*; 322A_01306) and cytochrome *c* oxidase, diheme subunit, membrane-bound (*fixP*; 322A_01305). Genes associated with nitrogen assimilation (glutamine synthetase, *glnA*; 322A_00568) and regulation of carbon storage (*csrA*; 322A_00279) were also downregulated by menadione. *glnA* contributes to glutamate metabolism, which – in concert with *rpoN* – controls acidic amino acid uptake and metabolism in bacteria (30).

A KEGG-based network analysis was undertaken with the significantly differentially expressed genes (Supplementary Figure 2, Supplementary Table 4). The degree of a gene within a network is the number of connections it has to other genes in the network. Most genes have a low degree, but a small number have a high degree: these “hub” genes are considered important in controlling biological processes. The mean degree across the entire gene network was 2.18 +/- 3.44. Genes associated with virulence (*cagA*, 322A_00555, degree 28) and stationary phase (*surE* – survival protein, 322A_01007, degree 28) were the most connected genes in the network (Supplementary Figure 2). Gene *cheY*, encoding a chemotaxis protein associated with two-component systems, was among the genes with highest degree (11), making the above-mentioned effect of menadione on flagellar assembly clearer (Figure 3a). That is, downregulation of *cheY* prevents full activation of genes associated with flagellar assembly and impedes motility. *In vitro* testing confirmed the inhibitory effect of menadione on motility of *H. pylori* (Figure 3b). Other genes with a high degree were associated with nitrogen assimilation (e.g. *glnA*, degree 9), energy generation or chemotaxis (Supplementary Table 4).

**Figure 3.**
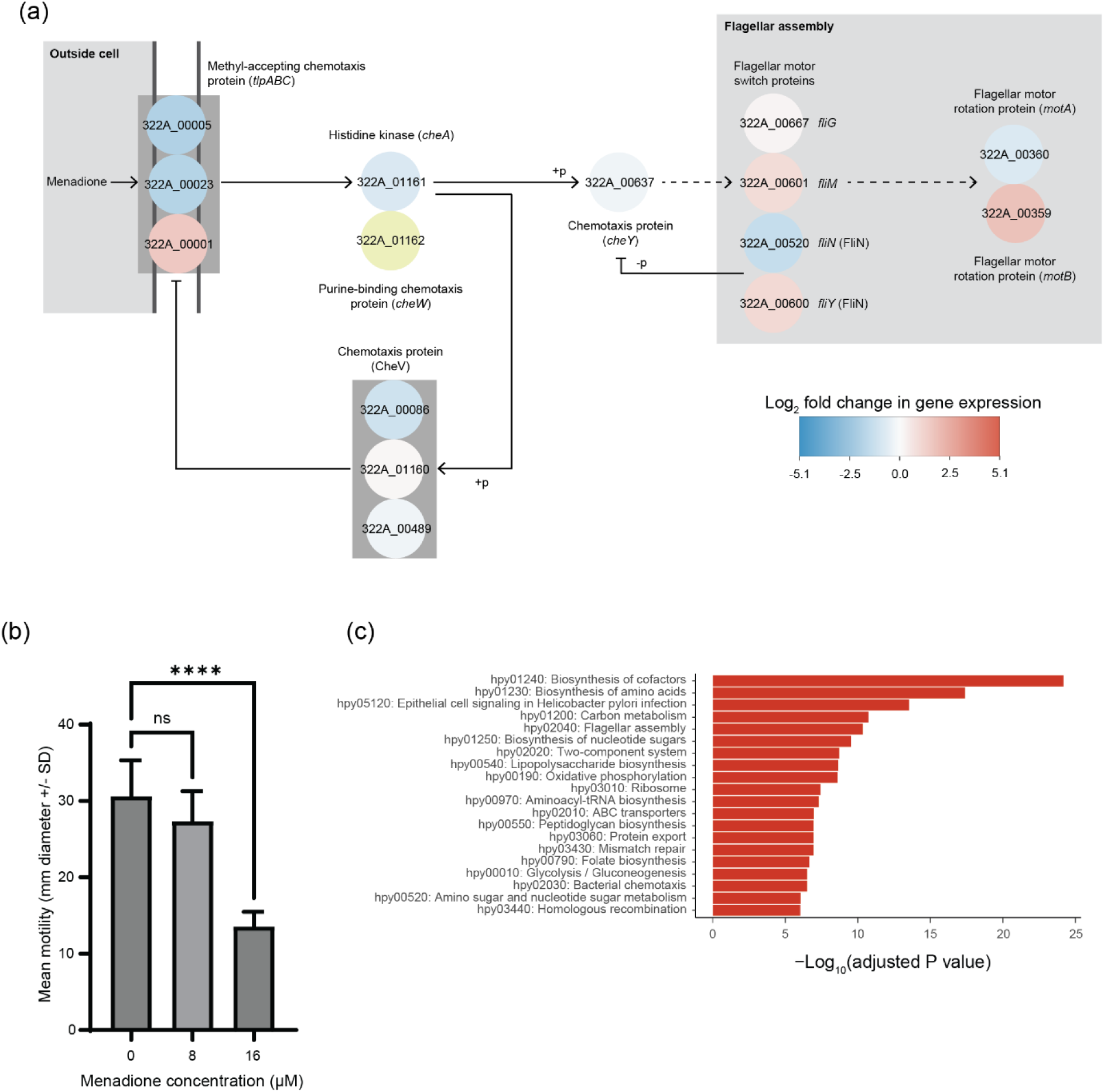
Effect of a sub-inhibitory concentration (16 µM) of menadione on the motility of *H. pylori* strain 322A. (a) The KEGG “Bacterial chemotaxis” pathway with gene expression data overlaid onto it. Significantly differentially expressed genes (adjusted P value < 0.05; Benjamini-Hochberg) are shown by blue (downregulated by menadione) and red (upregulated by menadione) circles, with log_2_-fold changes represented by the colours shown in the legend; non-significant data, yellow circle. (b) The effect of menadione on motility was determined by measuring the diameter of *H. pylori* strain 322A growth 7 days after 10 µl of a dense bacterial suspension was stabbed into the centre of soft agar plates (refer to Methods). Menadione at 16 µM significantly inhibited the motility of *H. pylori* strain 322A (**** = p < 0.0001, one-way ANOVA with Dunnett’s post-hoc test). Data shown are n = 9 (compiled from triplicate technical replicates, independently repeated three times). (c) Top 20 KEGG pathways in strain 322A significantly affected (adjusted P value < 0.05; Benjamini-Hochberg) by 16 µM menadione based on a gene over-representation analysis. Details of all significant pathways can be found in Supplementary Table 5.

Gene overrepresentation analysis using the significantly differentially expressed genes mapped to KEGG identified 77/98 metabolic pathways (P < 0.05) affected by the sub-inhibitory concentration of menadione (Figure 4c, Supplementary Table 5). After biosynthesis of co-factors and amino acids, “Epithelial cell signalling in *Helicobacter pylori* infection” was the most significant (adjusted P value < 0.05) over-represented metabolic pathway, incorporating 35/39 of the significantly differentially expressed genes and including 24/26 cag pathogenicity island (*cag*PAI) genes, *vacA*, *hpaA* and *ureA/B/I*. Also among the top 20 most significant pathways were “Flagellar assembly”, “Two-component system”, “Oxidative phosphorylation”, “Bacterial chemotaxis”, “Amino sugar and nucleotide sugar metabolism” and “Homologous recombination” (Figure 3c, Supplementary Table 5), supporting our findings from SPIA and the KEGG network-based analysis.

**Figure 4.**
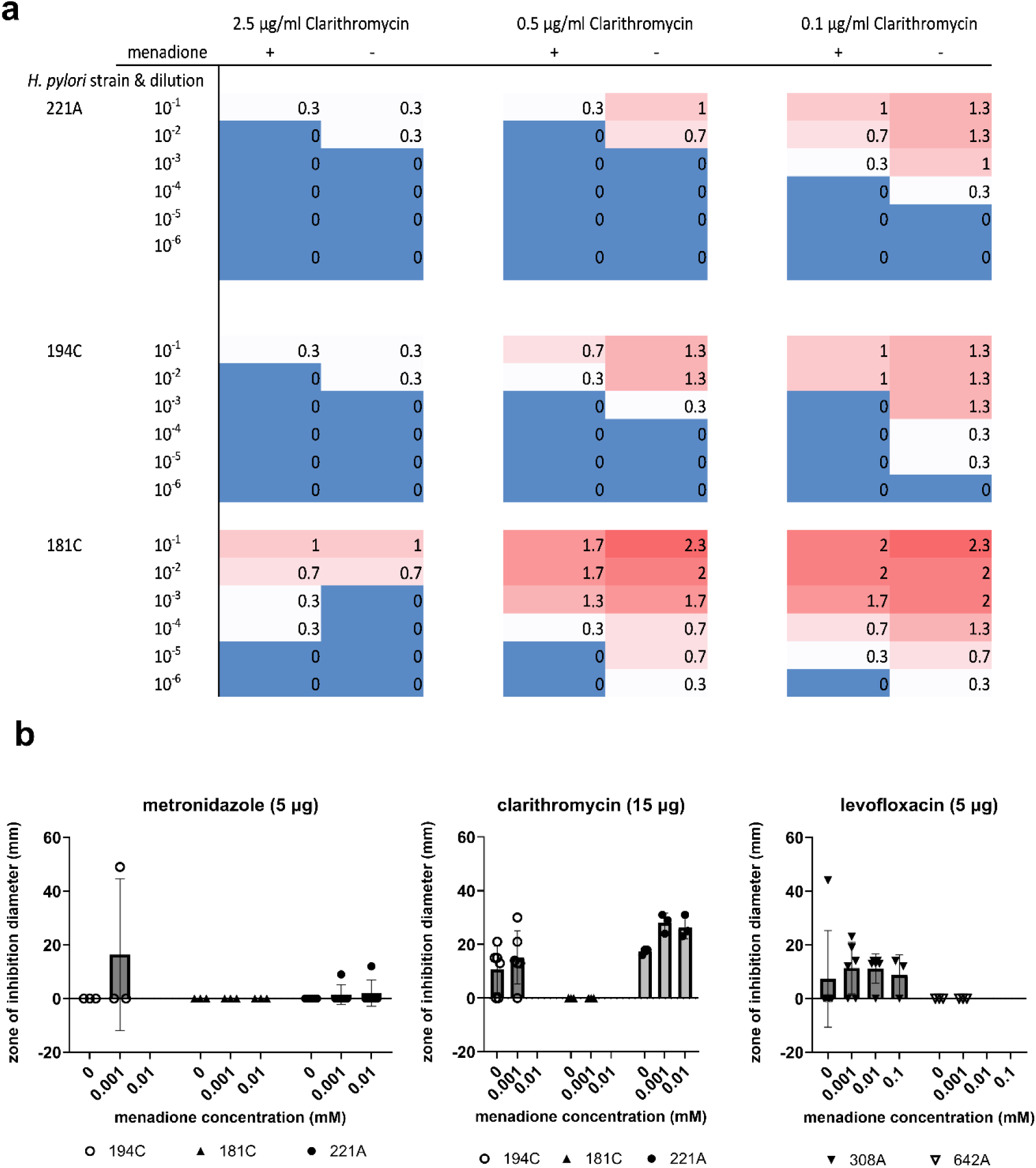
Effects of sub-MIC menadione on antibiotic susceptibility. (a) Modified agar dilution method. Numbers and colours indicate means of semi-quantitative scores for bacterial growth density from triplicate 10 µl spots where 0 = no growth, 3 = dense growth. Addition of menadione to the growth media caused increases in susceptibility to the antibiotic clarithromycin in most cases. (b) Modified disc diffusion method. Graphs show antibiotic susceptibilities in the presence and absence of menadione, as individual data points. Means +/- SD are also shown as grey bars and error bars. Data points were excluded from plates where the menadione concentration entirely inhibited bacterial growth. Refer to Methods for details of the modified agar dilution and disc diffusion assays.

### Sub-MIC menadione may reduce the antibiotic susceptibility of some *H. pylori* strains

A selection of *H. pylori* strains previously shown (4) to be clinically resistant to clarithromycin (excluding strain 322A, which is susceptible) were exposed to various concentrations of clarithromycin in the presence or absence of 16 µM and the bacterial viability was assessed semi-quantitatively after 48 h (Figure 4a). The presence of 16 µM menadione caused modest reductions in bacterial viability in most cases. In antibiotic disc diffusion assays, the presence of 1 to 10 µM menadione in the agar caused increases in antibiotic zone of inhibition diameter in some cases, but the effect varied with strain and with antibiotic type and did not reach statistical significance (Figure 4b). According to a proteome-based analysis (via the Resistance Gene Identifier of The Comprehensive Antibiotic Resistance Database), strain 322A does not encode any antimicrobial resistance genes, hence its transcriptomic data were not interrogated further in relation to effect of menadione in this context.

### Menadione is cytotoxic to gastric adenocarcinoma cells

Menadione cytotoxicity to human gastric adenocarcinoma (AGS) cells was assessed using a CellTiter Glo assay (Figure 5). The minimum inhibitory concentration we determined for *H. pylori* in this study, 313 µM menadione, and higher concentrations, caused complete loss of AGS cell viability by 4 h. A sub-MIC menadione concentration, 78 µM, was minimally cytotoxic after 4 h but caused significant loss of cell viability after 24 h.

**Figure 5.**
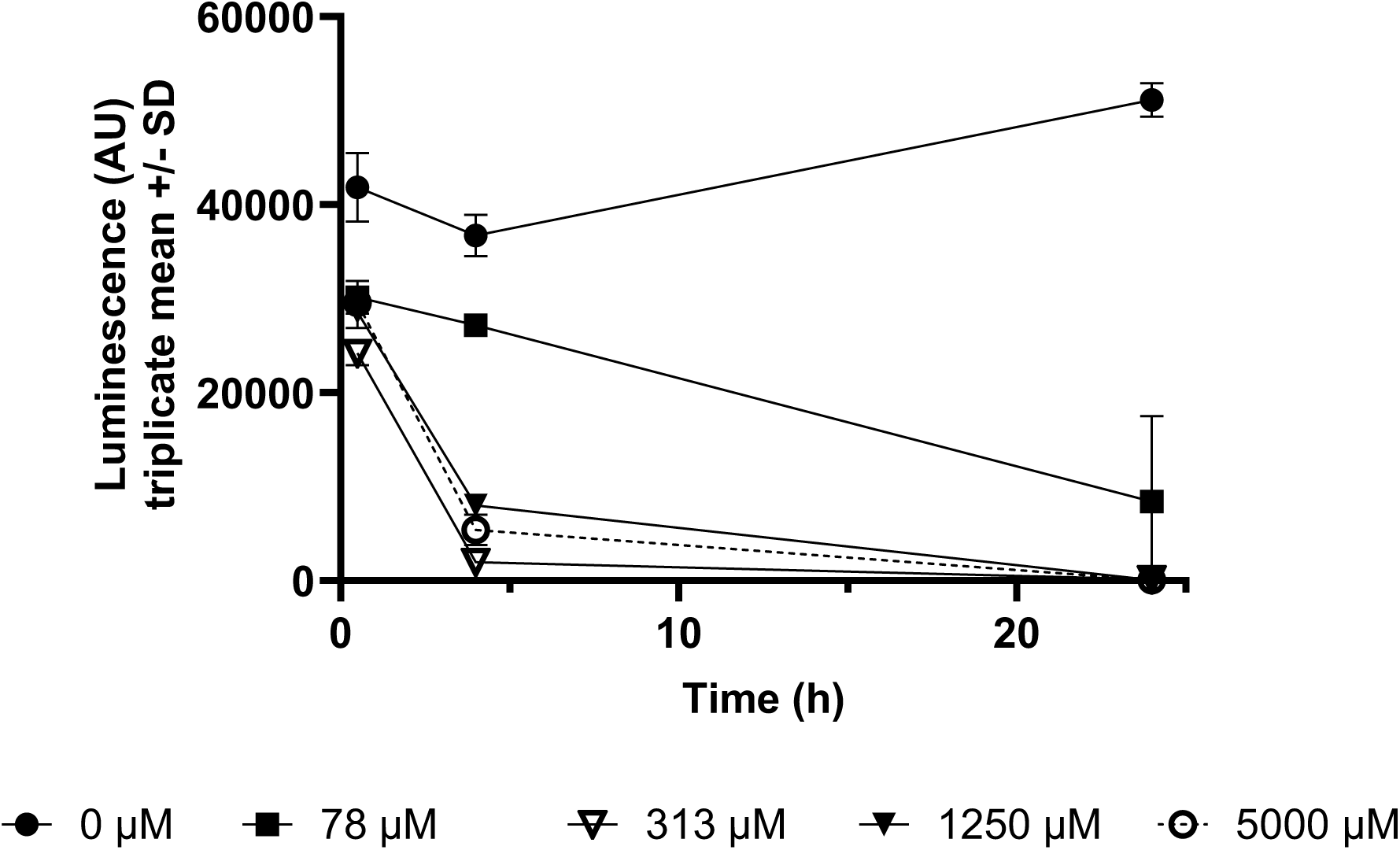
Cytotoxicity of menadione. AGS cells were treated with 0-5000 µM menadione for up to 24 h. To determine cell viability at each time point, CellTiter Glo reagent was added to the cell suspensions and luminescence (Arbitrary Units, AU) was measured after 10 min. Luminescence AU is directly proportional to the number of viable cells.

## Discussion

In this study, we confirmed the antimicrobial activity of menadione against *H. pylori* and showed that a sub-MIC (16 µM) concentration of menadione can significantly downregulate the expression of numerous virulence- and colonisation-associated genes, via a general stress response. We also found that *H. pylori* does not easily evolve resistance to menadione and that menadione may modulate *H. pylori* susceptibility to antibiotics. In addition, we confirmed that menadione has cytotoxic effects on gastric adenocarcinoma cells.

Several previous studies have shown that naphthoquinones, including menadione, inhibit the growth of *H. pylori.* Inatsu et al. (31) determined the MIC_90_ (concentration at which bacterial viability was reduced by 90 %) to be 3.2 µg/ml (∼20 µM) and Park et al. (13) reported strong inhibitory activity of menadione against *H. pylori* in disc diffusion assays with doses down to 5 µg/disc. Lee et al. (14) used agar dilution to determine the MIC of menadione against four laboratory strains and 38 clinical isolates of *H. pylori* and found it to be 8 µM in the reference strains and 2-8 µM in the majority of clinical isolates. In our laboratory, we found the MIC of menadione to be 313 µM for most clinical isolates we tested, using a broth microdilution assay in Brucella broth + 5 % FCS. These differences in MICs may be due to the different methods used, since both our own study and Inatsu et al. (31), which used broth microdilution methods, reported higher MICs than the studies that used agar-based methods. Future studies should consider applying more than one method to MIC determination and should also determine minimum bactericidal concentrations. Using nine strains from five patients, we found that MBCs were much higher than MICs indicating that menadione is bacteriostatic rather than bactericidal in the micromolar range. Menadione’s mode of action has recently been confirmed to be bacteriostatic in nature in *Escherichia coli* (32).

Culturing bacteria in the presence of sub-MIC concentrations of bacteriostatic or bactericidal agents is a commonly used method for generating resistant mutants in the laboratory (33,34). Our intention was to take a directed evolution approach, culturing *H. pylori* in the presence of increasing concentrations of menadione, to generate persistently resistant mutants and study menadione resistance mechanisms. However, none of our attempts to generate menadione-resistant mutants using a wide range of sub-MIC concentrations (>100 replicates at each concentration, all incubated for 48 h which is equivalent to approximately 16 generations given the estimated *H. pylori* doubling time of 3 h(35)) yielded detectable bacterial growth at concentrations above 313 µM. This suggests that *H. pylori* is not able to easily evolve resistance to menadione. Findings from our transcriptomic analyses (discussed below) may provide insights as to why.

The sub-MIC concentration of menadione used in this study had a profound effect on *H. pylori* strain 322A’s transcriptome, with the expression of almost 80 % of genes encoded by the bacterium significantly altered upon prolonged exposure (24 h) to the bacteriostatic agent (Figure 2). By using a combination of bioinformatics approaches to interpret our transcriptomic data, we were able to develop an in-depth understanding of how menadione affects the biological processes of *H. pylori*. Menadione elicits a general stress response –contributing to survival in the stationary phase – that is mediated by upregulation of *surE* (36) and *rpoN* (37,38). In our network-based analysis, the gene encoding for survival protein SurE – *surE* – was, along with *cagA*, the most important “hub” gene (Supplementary Figure 2). SurE is considered a virulence factor in *Brucella abortus*, due to its ability to maintain cell viability in stationary phase (39). Therefore, SurE may also be considered a virulence factor of *H. pylori* if it is found to be essential to the survival of the bacterium in its stationary phase. Because of its absence in mammals, SurE has been proposed as a novel drug target (39).

Upregulation of *dnaA* expression was intriguing, and may be part of the general stress response elicited by menadione. Although not annotated in KEGG, the expression of *hobA* (HP_1230; 322A_01058) was also significantly upregulated by menadione; HobA forms a complex with DnaA in *H. pylori*, and is thought to play a crucial scaffolding role during the initiation of replication in the bacterium (40). Antibiotic-induced replication stress elicits competence in some bacteria by increasing transcription of competence-associated genes around the origin of replication, with this considered a general molecular mechanism through which bacteria can challenge this type of insult (41). *H. pylori* encodes two operons of competence genes (*comB2*–*comB3*–*comB4*, annotated in 322A’s genome based on (42); *comB6*–*comB7*–*comB8*–*comB9*–*comB10*, annotated in 322A’s genome based on (43)) upstream of *dnaA*. The expression of 4/5 of the genes in the larger operon was significantly upregulated in the presence of menadione, along with *dprA*, necessary for transformation (43) (Figure 6). Competence is known to be activated in *Streptococcus pneumoniae* in response to antibiotics and/or DNA-damaging stressors (41,44). We are not aware of this process being associated with bacteriostatic agents, although menadione is known to damage DNA in mammalian cell lines (45). *H. pylori* is unusual among Gram-negative bacteria in having a constitutive competence state across all growth phases (46), with its competence increasing when the bacterium is exposed to DNA-damaging agents such as ciprofloxacin (47). As such, any stressor that could increase the competence of *H. pylori* requires further study.

**Figure 6.**
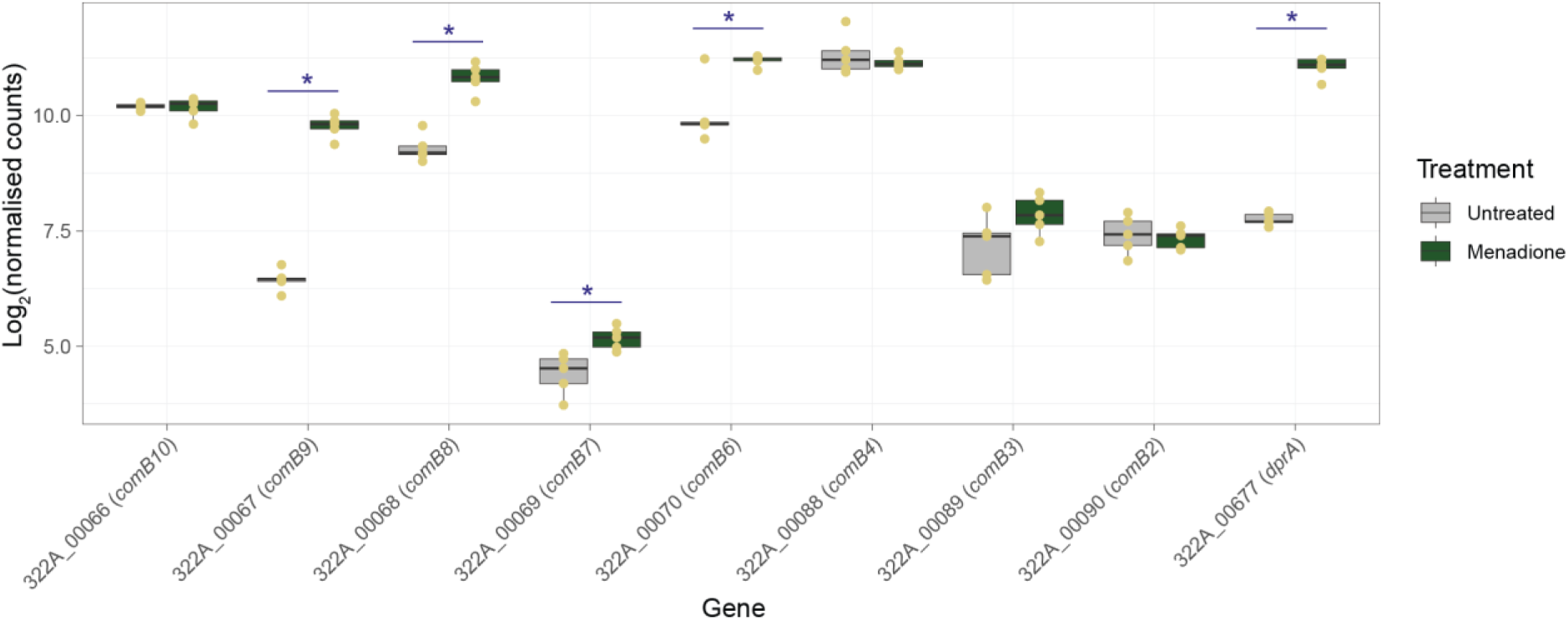
Effect of sub-inhibitory concentration of menadione (16 µM) on expression of *H. pylori* 322A competence-associated genes. Boxplots show data points (yellow dots, n = 5 per group) and median values, with the hinges corresponding to the first and third quartiles; the whiskers extend from the hinges no further than +/- 1.5 * interquartile range. *Significantly differentially expressed in menadione-treated group compared with untreated control: adjusted P < 0.05 (Benjamini-Hchberg). Both the *comB2*–*comB3*–*comB4* operon (positions 92504–95413, contig 1) and the *comB6*–*comB7*–*comB8*–*comB9*–*comB10* operon (position 68539–72581, contig 1) are upstream of *dnaA* (positions 189299–190666, contig 1), with both operons in reverse orientation. *dprA* (positions 100320–101120, contig 4) is not found close to the origin of replication in 322A.

Various analyses of KEGG data (SPIA, network and over-representation) supported by phenotypic (motility) data demonstrated menadione significantly affected the chemotaxis and flagellar assembly of *H. pylori* strain 322A (Figure 3). Stable accumulation of sigma-54 factor (*rpoN*) is required for the expression of several flagellar genes in *H. pylori*, and is dependent on the expression of the little-studied protein HP_0958 (48). In our study, expression of HP_0958’s encoding gene (322A_00976) was significantly downregulated (log_2_-fold change -0.48, adjusted P value = 0.002; Supplementary Table 2) in the presence of menadione. The role of HP_0958 in sigma-54 factor accumulation was first demonstrated in the stress response of *H. pylori* ATCC 43504 to nutrient limitation (48).

Expression of *rpoN* in stationary phase also downregulates genes associated with energy metabolism and biosynthetic processes in *H. pylori* strain 26695, with concomitant upregulation of genes linked to protein processing and redox reactions (37). Our transcriptomic analyses confirm these previous findings. Hence, here we demonstrate that the role of the sigma-54 factor stress response extends to bacteriostatic agents such as menadione.

The virulence- and colonisation-associated genes *luxS* and *ruvC* were all significantly downregulated by sub-MIC menadione (16 µM in broth culture). LuxS, associated with biosynthesis of amino acids and quorum sensing, may play a role in cysteine synthesis in *H. pylori*(49), and deletion of the *luxS* gene reduced bacterial motility and colonisation ability in rodent infection models(50,51). RuvC (Holliday junction endodeoxyribonuclease) plays many roles in *H. pylori*. Inactivation of *ruvC* reduced the rate of homologous recombination, increased bacterial sensitivity to levofloxacin, metronidazole, and oxidative stress, and impaired colonisation in a mouse infection model(52).

*H. pylori* uses urease to catalyse the conversion of urea to ammonium, buffering the pH in its immediate vicinity(53,54). *ureA* and *ureB* are structural urease genes(55) while the accessory gene *ureI* encodes a urea channel required for import of urea to the cytoplasmic location of the majority of urease enzyme(56). Mutants of *ureA, ureB* or *ureI* are avirulent or have impaired colonisation in animal models of infection(57–59). In addition to its role in acid survival in the human stomach, urease activity may contribute to virulence through the disruption of epithelial tight junctions(60,61) and stimulation of pro-inflammatory cytokine production(62,63). The *ureA/B/I* genes were all significantly downregulated by menadione in our study.

Extensive previous work in the *H. pylori* field has proven the importance of CagA and VacA in bacterial virulence and the development of peptic ulcer disease and gastric cancer in *H. pylori-*infected people. Infection with type I strains (*cagA+* and expressing s1i1m1 forms of *vacA*) is more strongly associated with ulcer disease and cancer than infection with strains lacking these virulence factors(1–3). CagA is an oncoprotein that is injected into gastric epithelial cells by a type IV secretion system, the components of which are encoded by the *cag*PAI(64–66). VacA is a secreted multifunctional cytotoxin that induces vacuolation, autophagy and apoptosis in gastric epithelial cells, increases epithelial barrier permeability, and inhibits T cell activity (reviewed by (67,68)). Expression of *cagA* and *vacA* were significantly downregulated by menadione in our study.

Our findings from whole genome transcriptomic analysis are consistent with those of a previous study that used RT-PCR on a subset of genes to show that menadione can downregulate the expression of virulence-associated genes in *H. pylori.* Lee et al. (14) reported that *H. pylori* treated with menadione had significantly decreased expression of *vacA* and also the *secA* gene involved in VacA toxin secretion. Expression of *cagA* was not significantly decreased (unlike our study) but expression of the *virB2, virB7, virB8, virB10* genes involved in CagA injection into host cells via the *cag* type IV secretion system was reduced. They also used co-culture assays and Western blotting to show that CagA and VacA protein translocation to AGS cells and the consequent cellular effects of these virulence factors (hummingbird phenotype and vacuolation) were reduced after pretreatment of *H. pylori* with menadione.

Gene over-representation analysis was used to map the genes that were significantly differentially expressed in response to menadione treatment to KEGG *H. pylori* biological pathways. Among the most significant pathways was “Epithelial cell signalling in *Helicobacter pylori* infection”. This pathway encodes numerous known virulence- and colonisation-associated genes of *H. pylori* including *ureA/B/I, vacA, cagA* and *hpaA*, and other genes of the *cag*PAI. The effects of menadione on the genes within this pathway, and how these might influence *H. pylori* survival and virulence within the human host, are summarised in Supplementary Figure 3. We propose that the down-regulation of *cagA* caused by menadione could prevent damage to host epithelial cells, while down-regulation of *vacA* could prevent or reduce ulcerogenesis, vacuolation, and apoptotic effects associated with *H. pylori* infection. Down-regulation of urease-associated genes could limit *H. pylori’s* ability to protect itself from host stomach acid, reducing its infective potential.

Menadione is known to have cytotoxic effects on a range of different cancer cell types and has been extensively studied for its potential utility as an anti-cancer agent(7,9,10,69). Consistent with our finding that menadione at 78 µM significantly reduced the viability of gastric adenocarcinoma (AGS) cells, previous studies have shown that 15 µM menadione reduced the growth of AGS cells by inducing G2/M cell cycle arrest(70) and concentrations at or above 20 µM induced apoptosis(71), while similar doses had no inhibitory effect on a non-cancerous gastrointestinal cell line. Suresh et al. (8) also showed that menadione was more cytotoxic to oral squamous carcinoma cells (IC_50_ = 8.45 µM) than to non-tumorigenic HEK293 (IC_50_ = 98.5 µM) and HaCaT (IC_50_ = 74.5 µM) cells.

In modified disc diffusion and agar dilution assays, we found that sub-MIC menadione caused some increases in *H. pylori* susceptibility to antibiotics, particularly clarithromycin, but the effects were modest and highly variable between strains. Menadione has previously been shown to reduce the MIC of antibiotics against multidrug-resistant strains of *Staphylococcus aureus*, *Escherichia coli* and *Pseudomonas aeruginosa*(12), and had good synergistic activity with several antibiotics against *Acinetobacter baumannii*(72). Menadione has also been shown to increase the susceptibility of *Stenotrophomonas maltophilia* to fluoroquinolones(73) and to induce expression of a multidrug efflux pump in *S. maltophilia*(74). Since we have shown that sub-MIC menadione causes extensive changes in gene expression, more definitive studies of the effects of menadione on antibiotic susceptibility in *H. pylori* are recommended.

Our study was limited by sample size, in particular for the RNA sequencing where only one strain and one menadione concentration were tested, and for the antibiotic susceptibility assays in which a limited number of clinically resistant isolates were available for each antibiotic. *H. pylori* strains are highly polymorphic, and it is possible that different strains may respond differently to menadione, so future studies should test multiple strains to confirm whether the results are consistent. Our use of 2-fold dilution series limited the precision with which we could determine MIC, MBC, and cytotoxic concentrations of menadione in our assays and further work is needed to define these more precisely using a range of methodological approaches, strains, and cell types.

Our findings raise the possibility that low doses of menadione could suppress *H. pylori* virulence and persistence by downregulating bacterial expression of virulence- and colonisation-associated genes. In addition, the bacteriostatic, antibiotic susceptibility modulating and adenocarcinoma cell cytotoxic effects of menadione make it an interesting prospect for further study.

## Data availability

RNAseq data reported on in this study are available to download from ArrayExpress under accession E-MTAB-14439. Supplementary materials associated with this publication are available from https://figshare.com/projects/Menadione_reduces_the_expression_of_virulence-_and_colonisation-associated_genes_in_Helicobacter_pylori/219631.

## Author statements

### Author contributions

Thompson – Data curation, Formal analysis, Investigation, Methodology, Visualization, Writing – original draft, Writing – review & editing. Omoyemi – Investigation, Formal analysis, Writing – review & editing. Hoyles – Data curation, Formal analysis, Investigation, Methodology, Visualization, Writing – original draft, Writing – review & editing. Winter – Conceptualization, Data curation, Formal analysis, Funding acquisition, Investigation, Methodology, Project administration, Supervision, Visualization, Writing – original draft, Writing – review & editing.

### Ethical approval

*H. pylori* strains were isolated from gastric biopsies donated by patients attending Queens Medical Centre Hospital, Nottingham, UK with written informed consent and ethical approval from the Nottingham Research Ethics Committee 2(08/H0408/195)

## Acknowledgements

The *H. pylori* strains used in this study were kindly provided by Profs Karen Robinson and John

Atherton, University of Nottingham, UK. We also thank Ms Melanie Lingaya, Mrs Joanne Rhead and Mr Macsen Fryer for technical support.

---

**Supplementary Figure 1.**
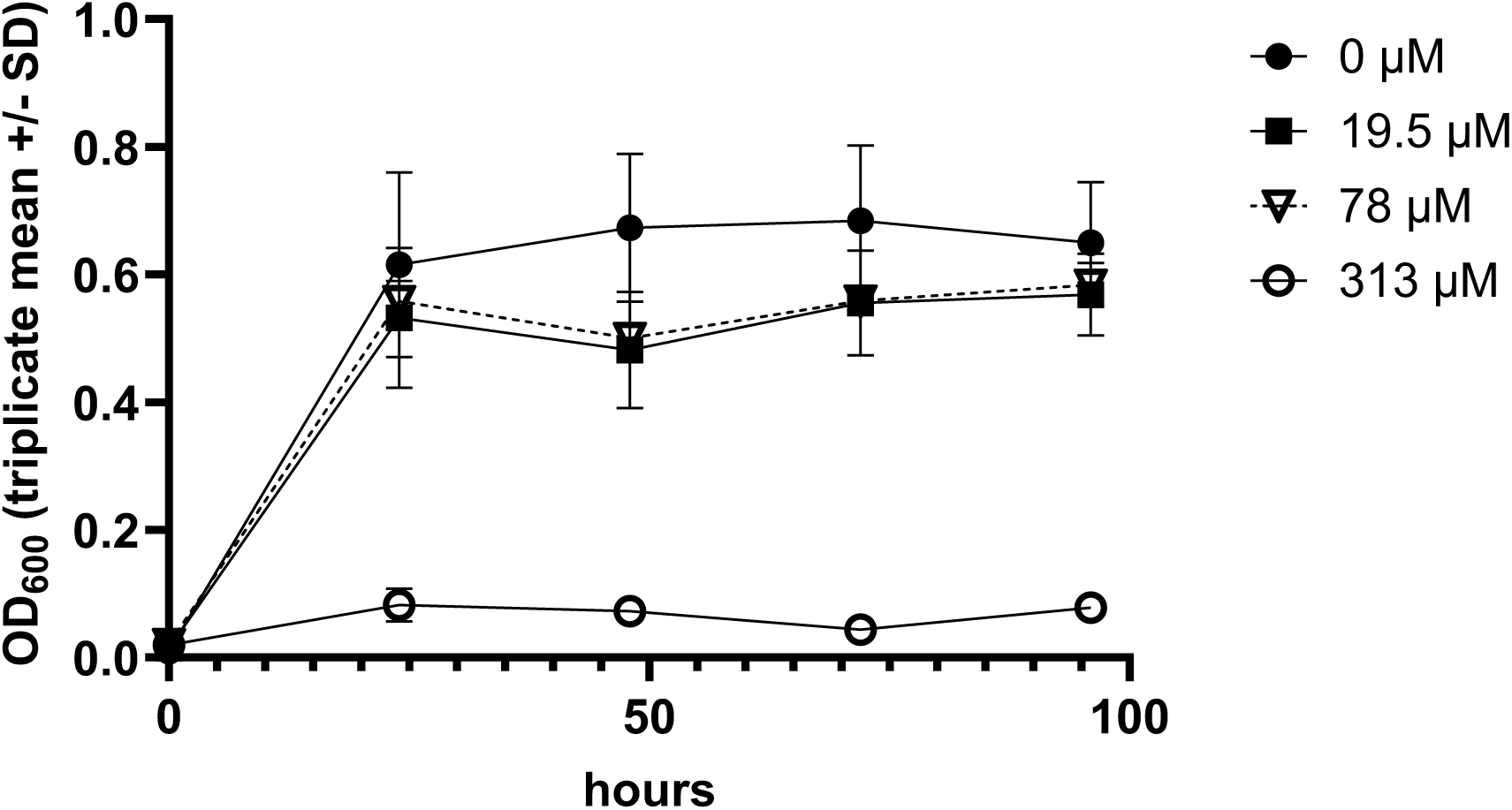
Effects of menadione on the growth of *H. pylori* strain 322A. *H. pylori* strain 322A was incubated in Brucella broth + 5 % heat-inactivated foetal calf serum, with the indicated concentrations of menadione, final volume 100 µl/well in 96-well plates at 37 °C under microaerobic conditions. Bacterial growth was monitored by measuring the OD_600_ at 0, 24, 48, 72 and 96 h. Data shown are blank corrected means +/- SD for triplicate wells. Menadione at 313 µM significantly inhibited *H. pylori* growth, but the modest reductions in OD_600_ with 78 µM and 19.5 µM menadione did not reach statistical significance (p>0.05, 2-way ANOVA with Dunnett’s post-hoc test).

**Supplementary Figure 2.**
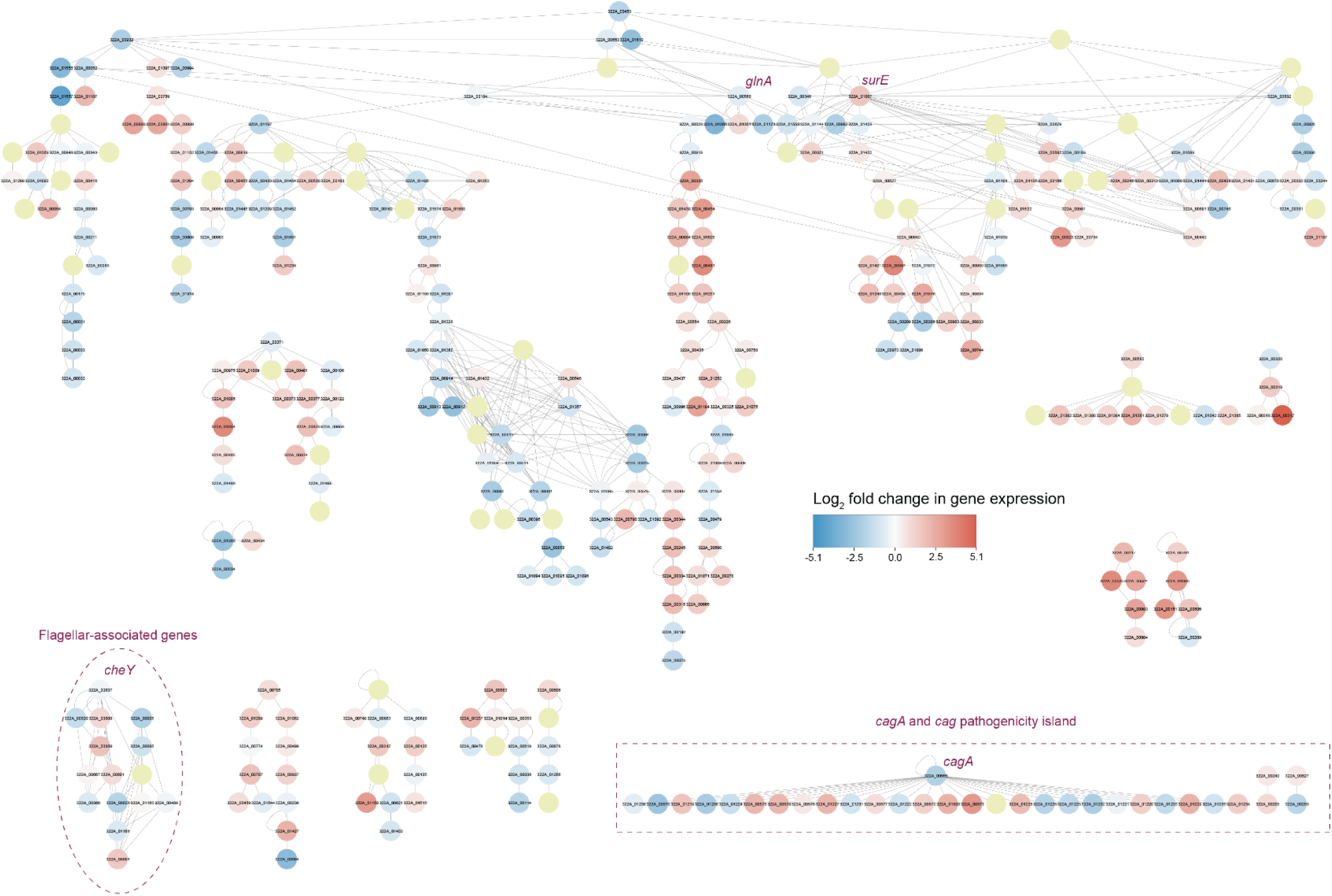
Network-based analysis of strain 322A genes that could be mapped to KEGG metabolic pathways. Significantly differentially expressed genes (adjusted P < 0.05; Benjamini-Hochberg procedure) are shown by blue (downregulated by menadione) and red (upregulated by menadione) dots; non-significant data are shown by yellow circles. Singletons have been removed from the network to reduce complexity.

**Supplementary Figure 3.**
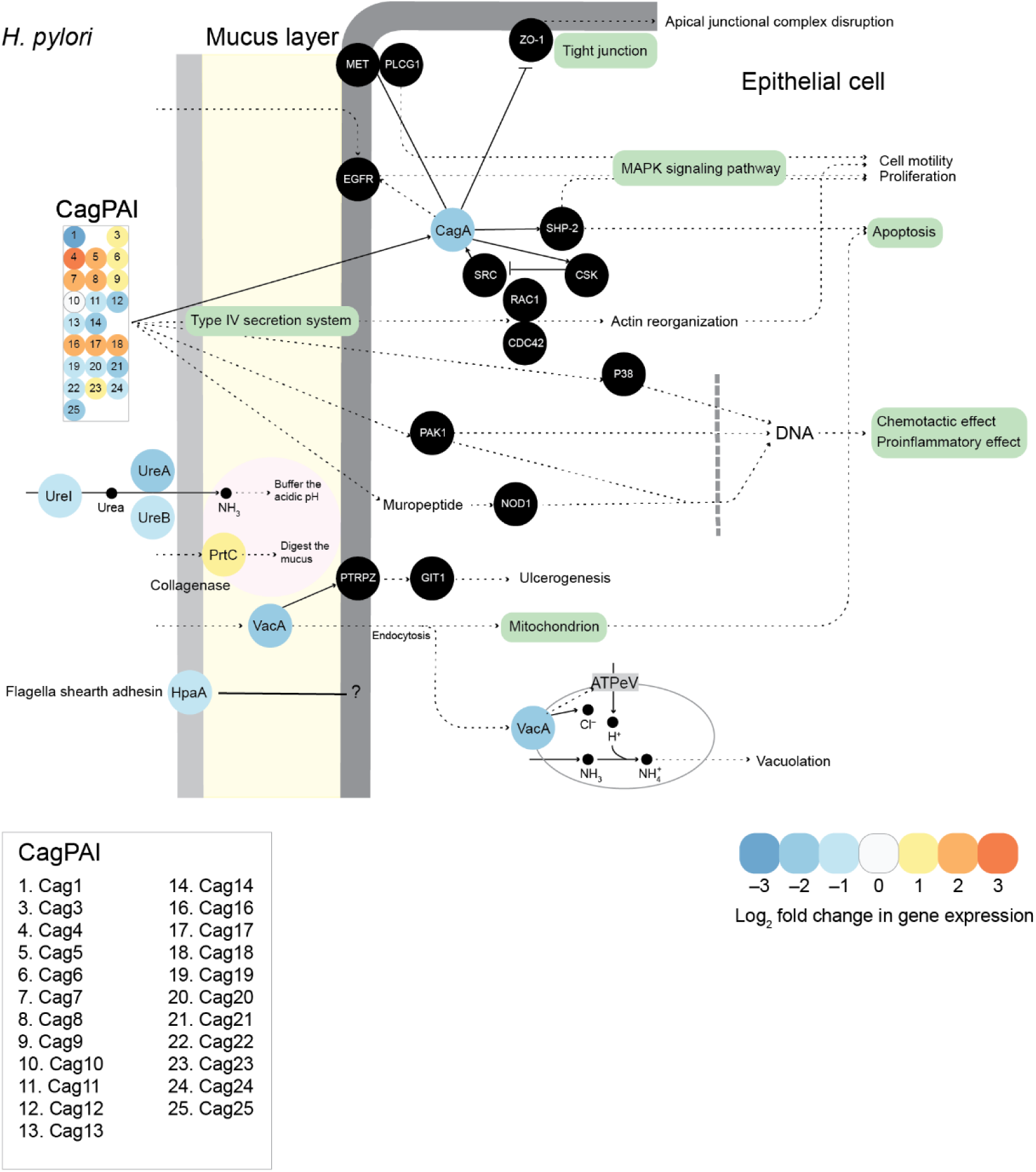
Differential gene expression in the KEGG pathway “Epithelial cell signalling in *Helicobacter pylori* infection” in response to menadione (16 µM) treatment. This pathway, including the major virulence factor-encoding genes *vacA, cagA* and the genes of the *cag*PAI, was one of the most over-represented metabolic pathways associated with the genes that were significantly affected in menadione-treated *H. pylori* cells. Proposed mechanism of action is as follows. Down-regulation of *cagA* prevents inhibition of ZO-1, maintaining tight junction integrity in host epithelial cells, along with prevention activation of MAPK signalling. Down-regulation of *vacA* prevents or reduces ulcerogenesis, vacuolation, and apoptotic effects associated with *H. pylori* infection. Down-regulation of urease-associated genes limits *H. pylori’s* ability to protect itself from host stomach acid. *H. pylori* gene products in the epithelial cell signalling pathway are shown as circles coloured by the log_2_-fold change in expression levels of their genes in this study (refer to colour legend in figure). Black circles with white text represent host-associated gene products not assayed in this study but with which *H. pylori* gene products interact (note: not all host-associated gene products included in the pathway are shown to aid clarity). The pathway was created based on *H. pylori-*specific information available from KEGG on 3 September 2024.

**Supplementary Table 1.** Minimum inhibitory concentrations (MIC) and minimum bactericidal concentrations (MBC) of menadione against *H. pylori* strains.

| *H. pylori* strain | MIC (mM) | MBC (mM) |
| --- | --- | --- |
| 194A | 0.16 | 1.3 |
| 194C | 0.31 | 2.5 |
| 265A | 0.31 | 2.5 |
| 265C | 0.31 | 2.5 |
| 295A | 0.31 | 2.5 |
| 295C | 0.31 | 2.5 |
| 322A | 0.31 | not tested |
| 322C | 0.31 | not tested |
| 326A | 0.16 | 1.3 |
| 326C | 0.31 | 2.5 |
| 439A | 0.31 | 2.5 |
| 791A | 0.31 | not tested |
| 791C | 0.31 | not tested |

The number in the strain name refers to the patient while A indicates an isolate from the antrum and C indicates an isolate from the corpus of the patient's stomach. The menadione concentrations tested were 10, 5, 2.5, 1.25, 0.63, 0.31, 0.16, 0.08, 0.04, 0.02 mM alongside appropriate bacterial growth, sterility, and menadione negative controls. All strains and concentrations were tested in triplicate.

**Supplementary Table 2.**
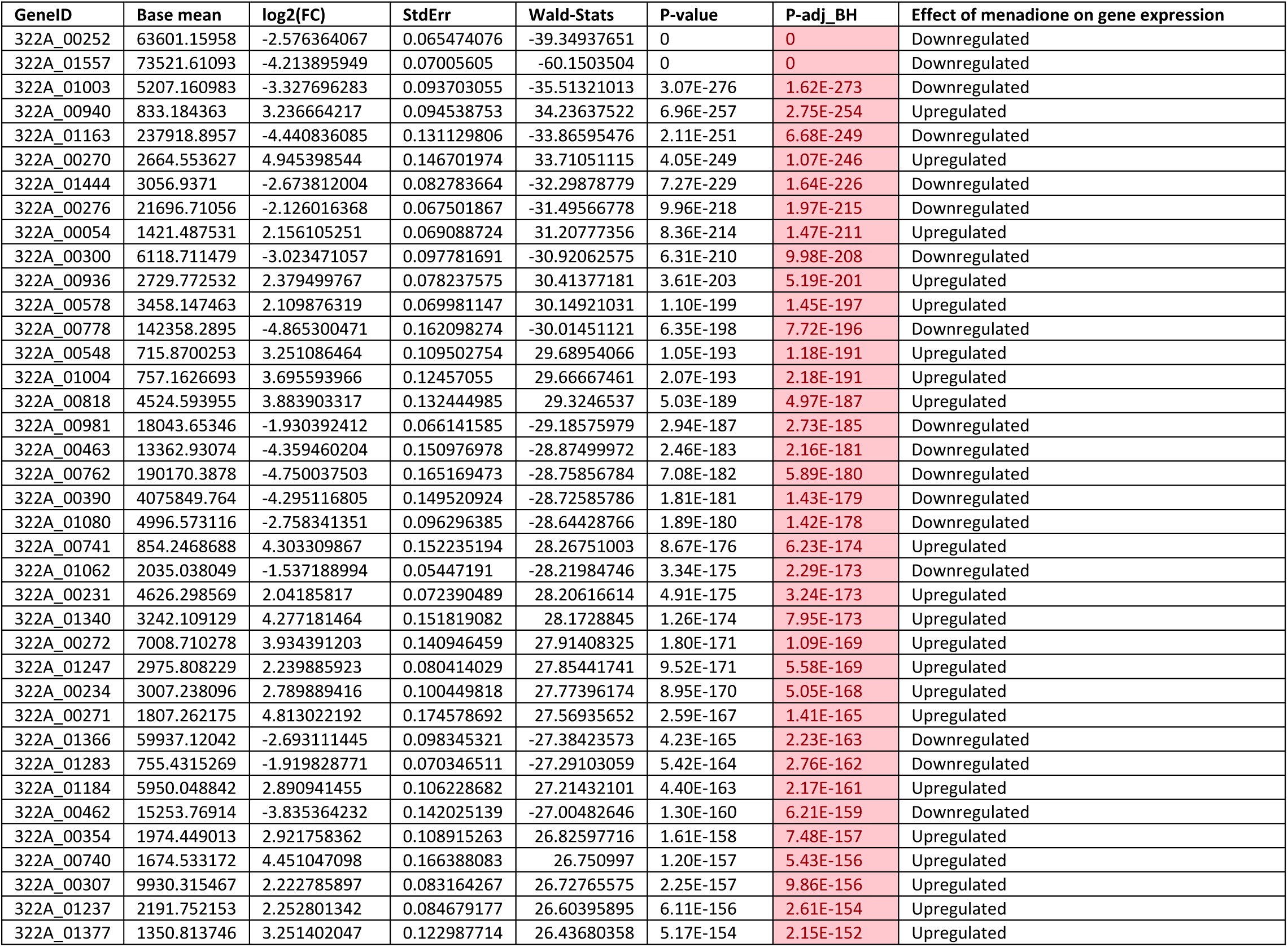
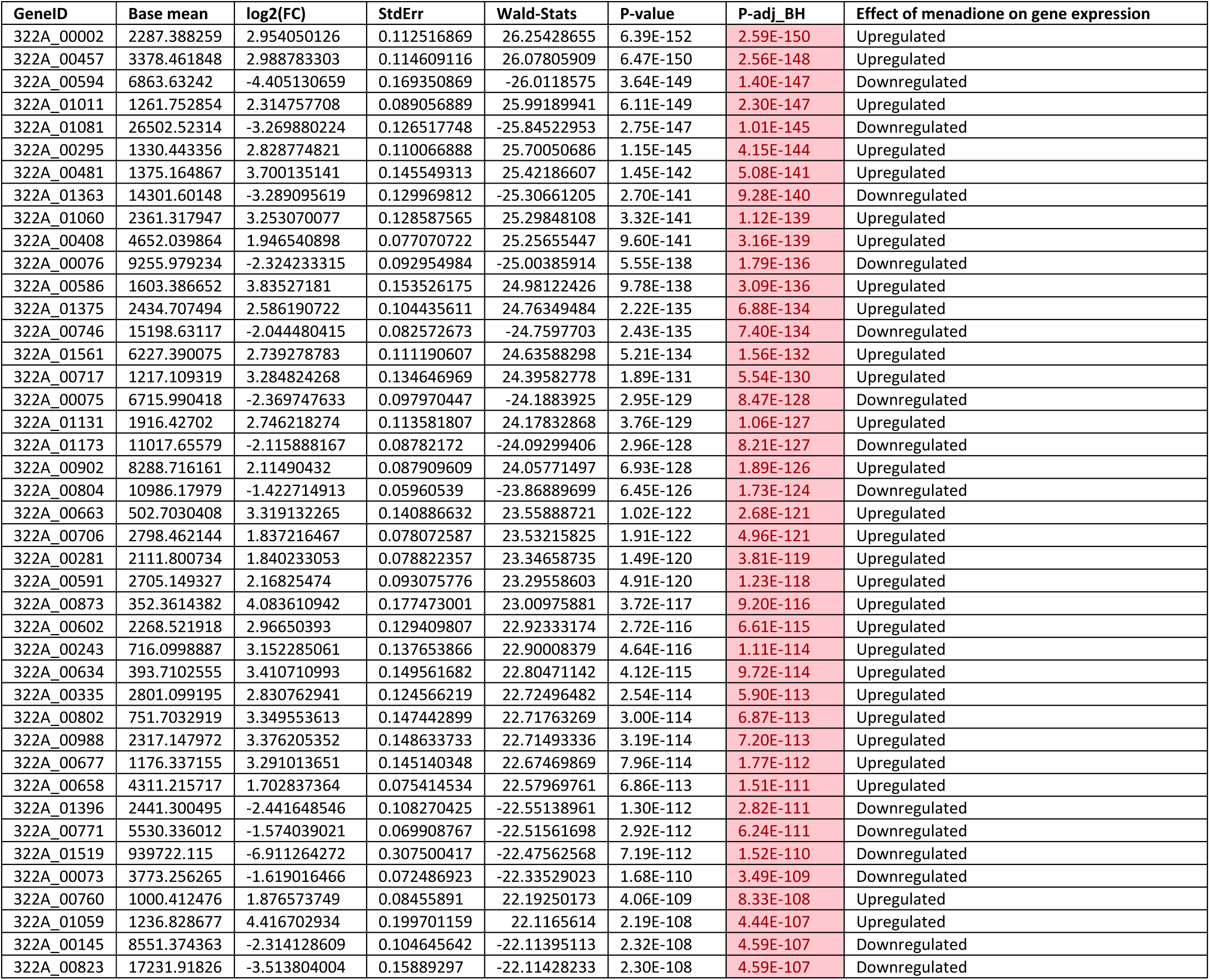
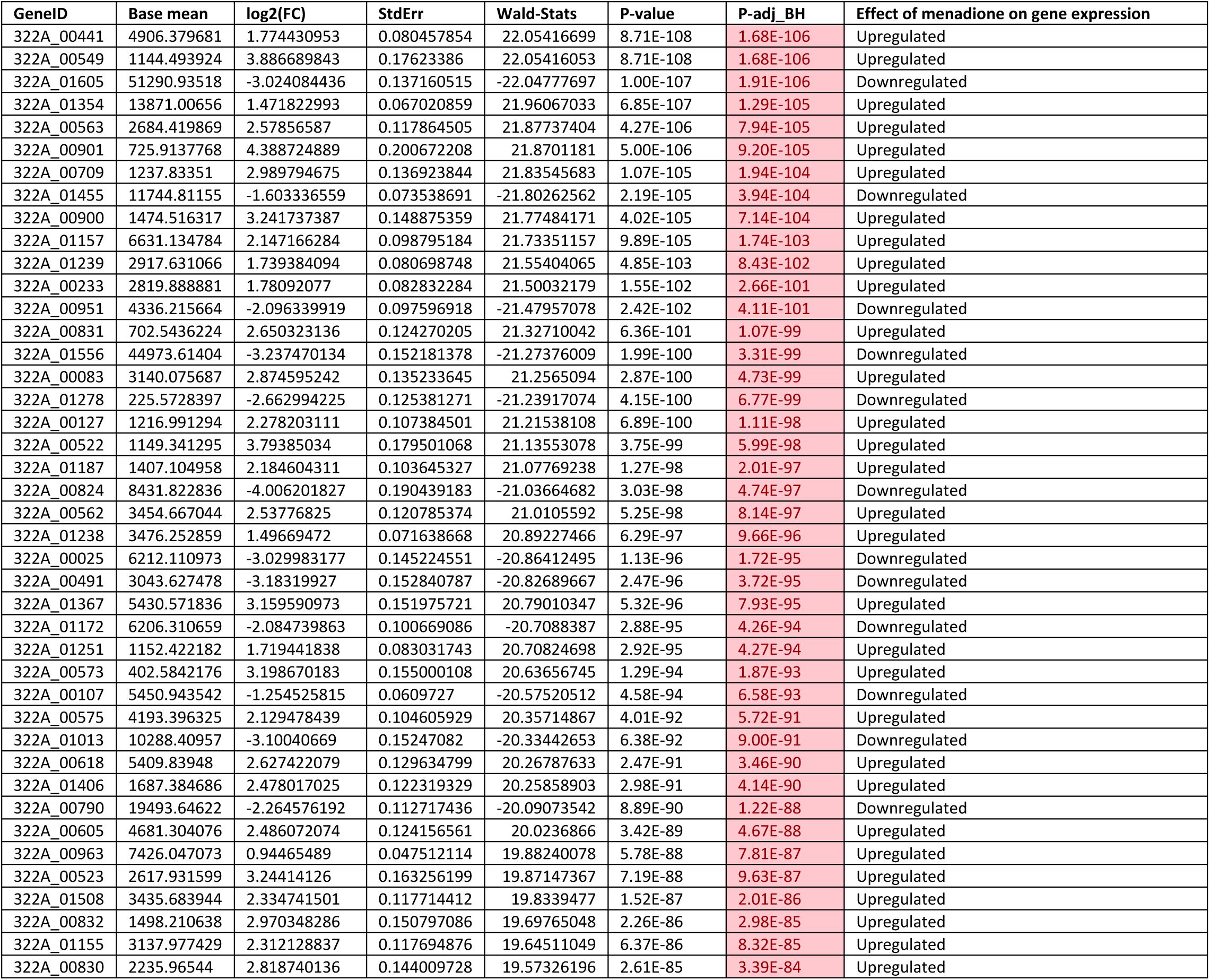
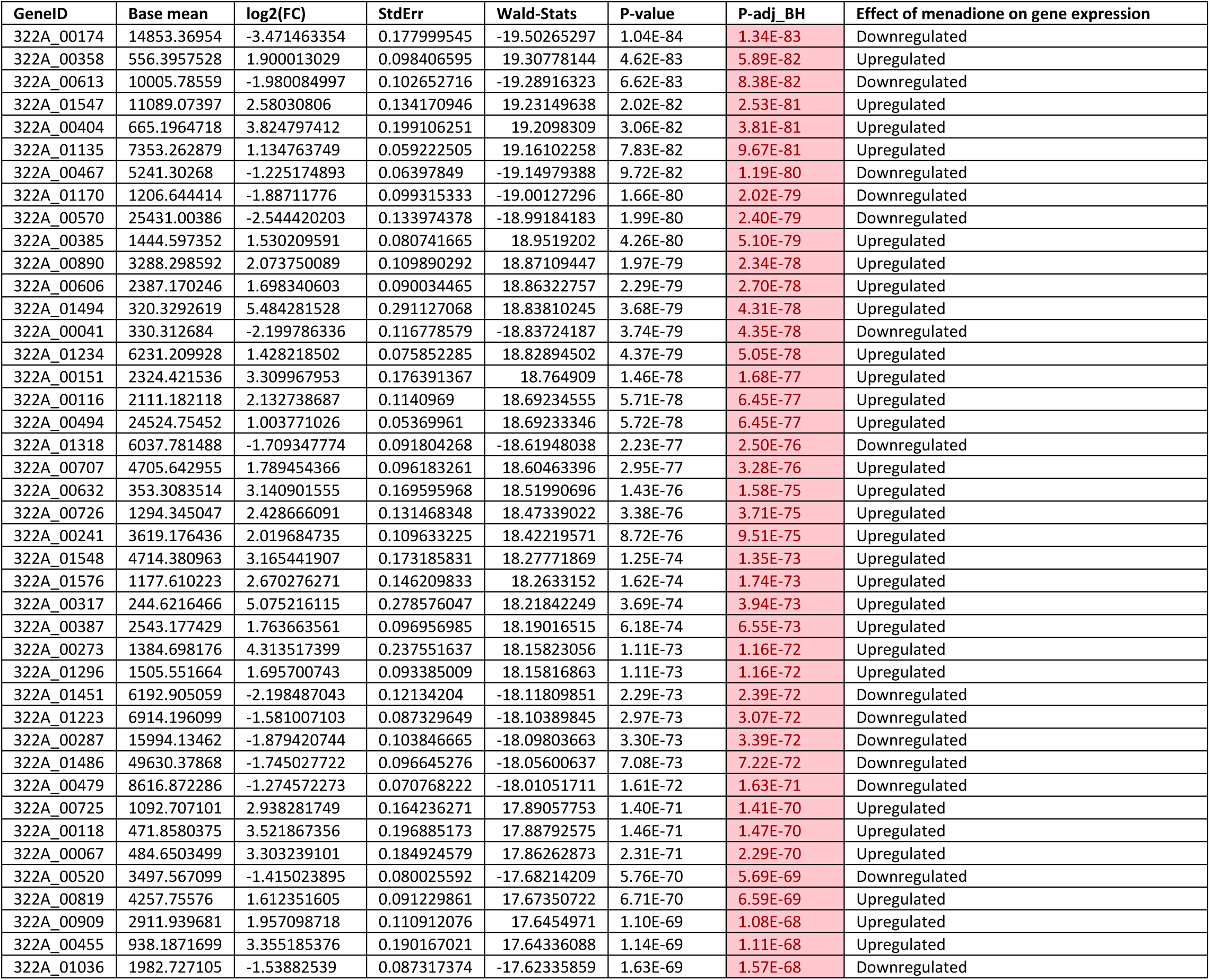
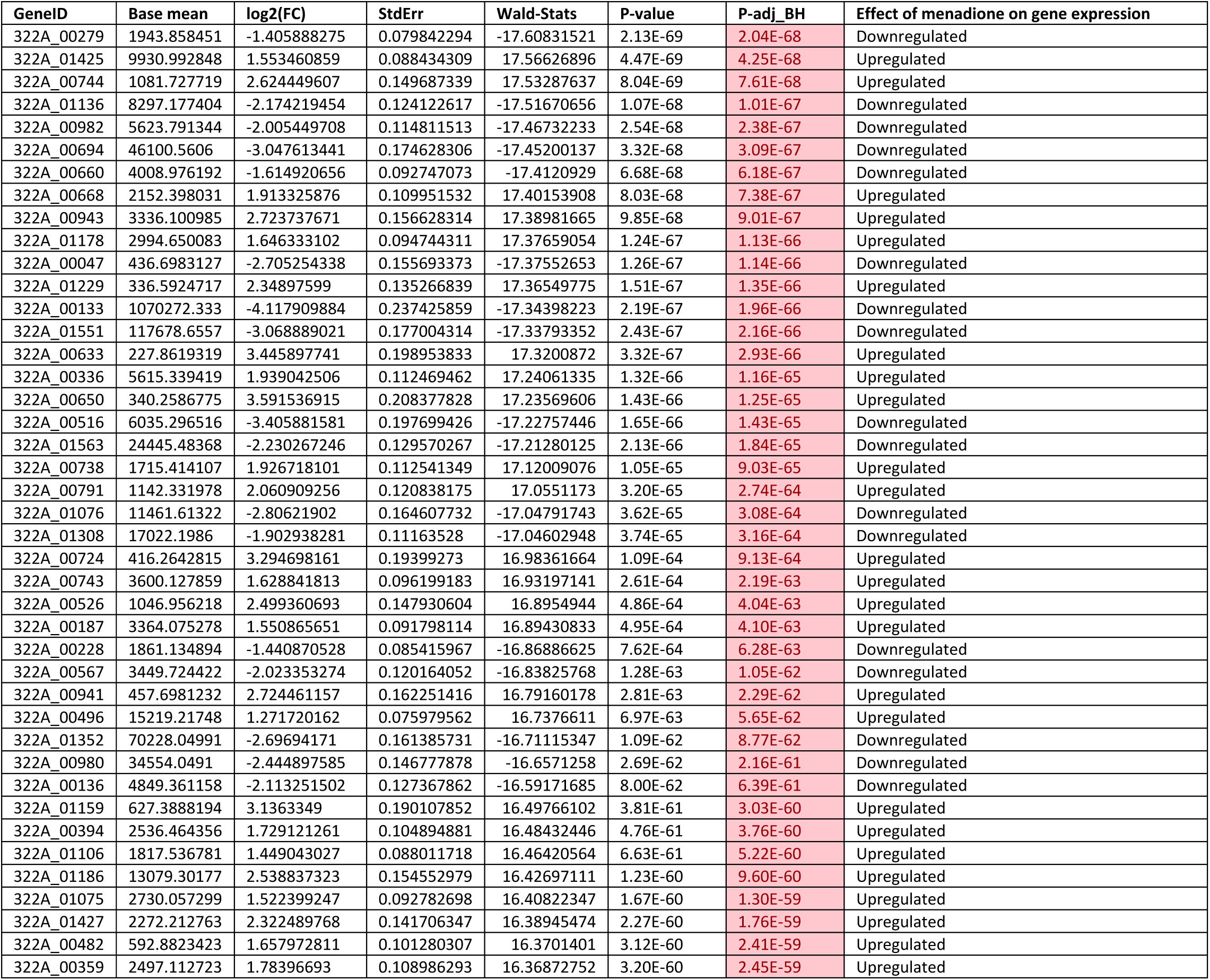
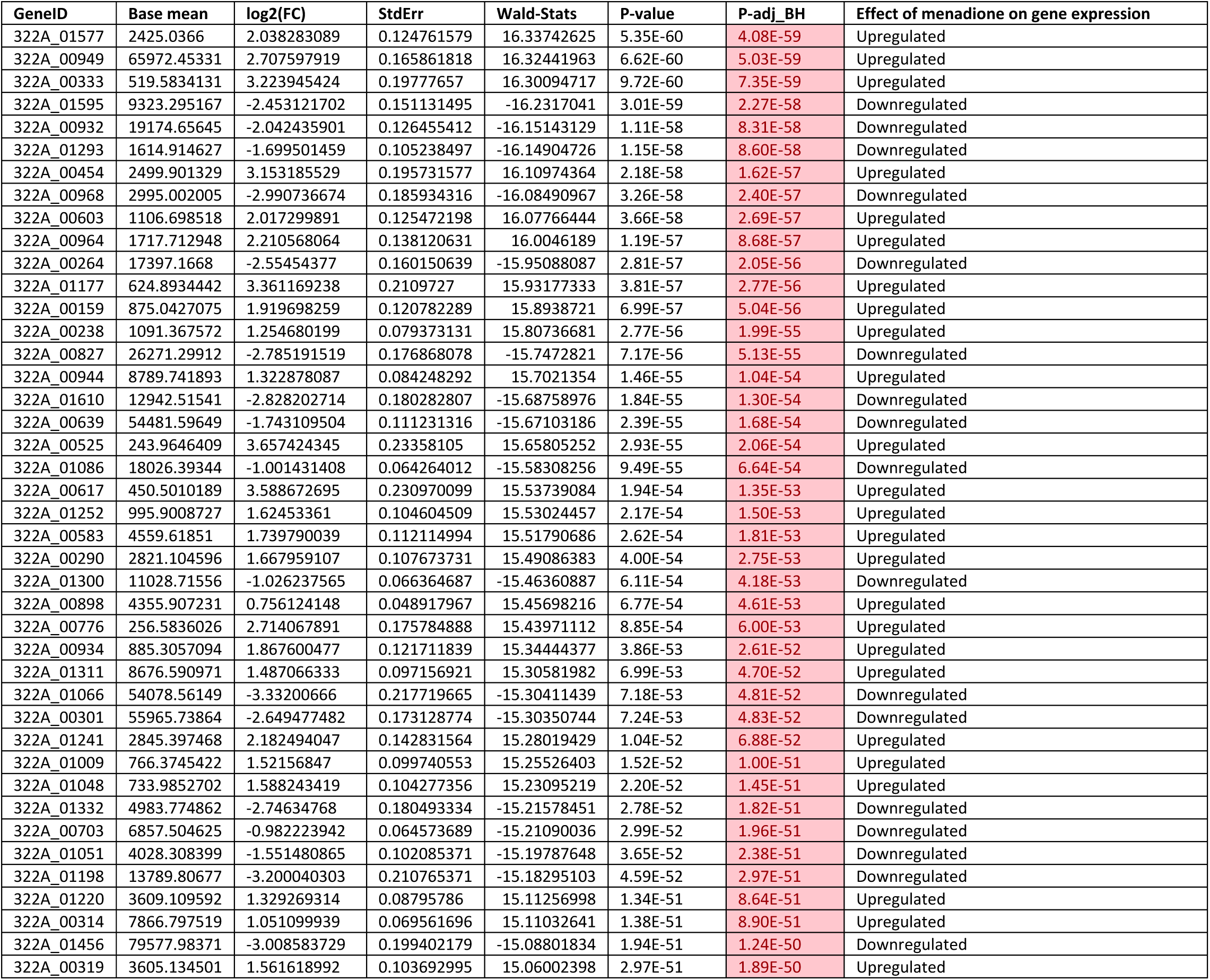
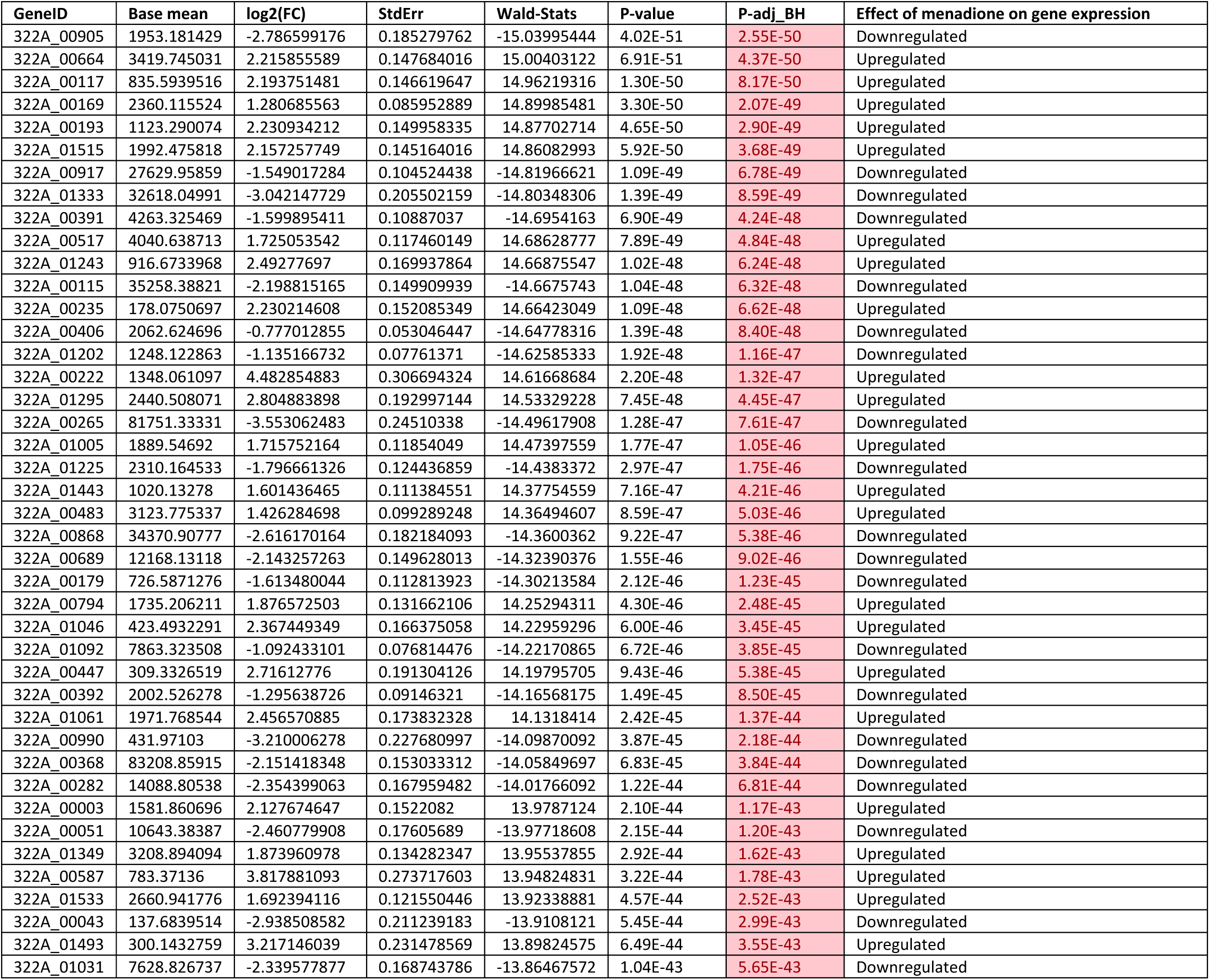
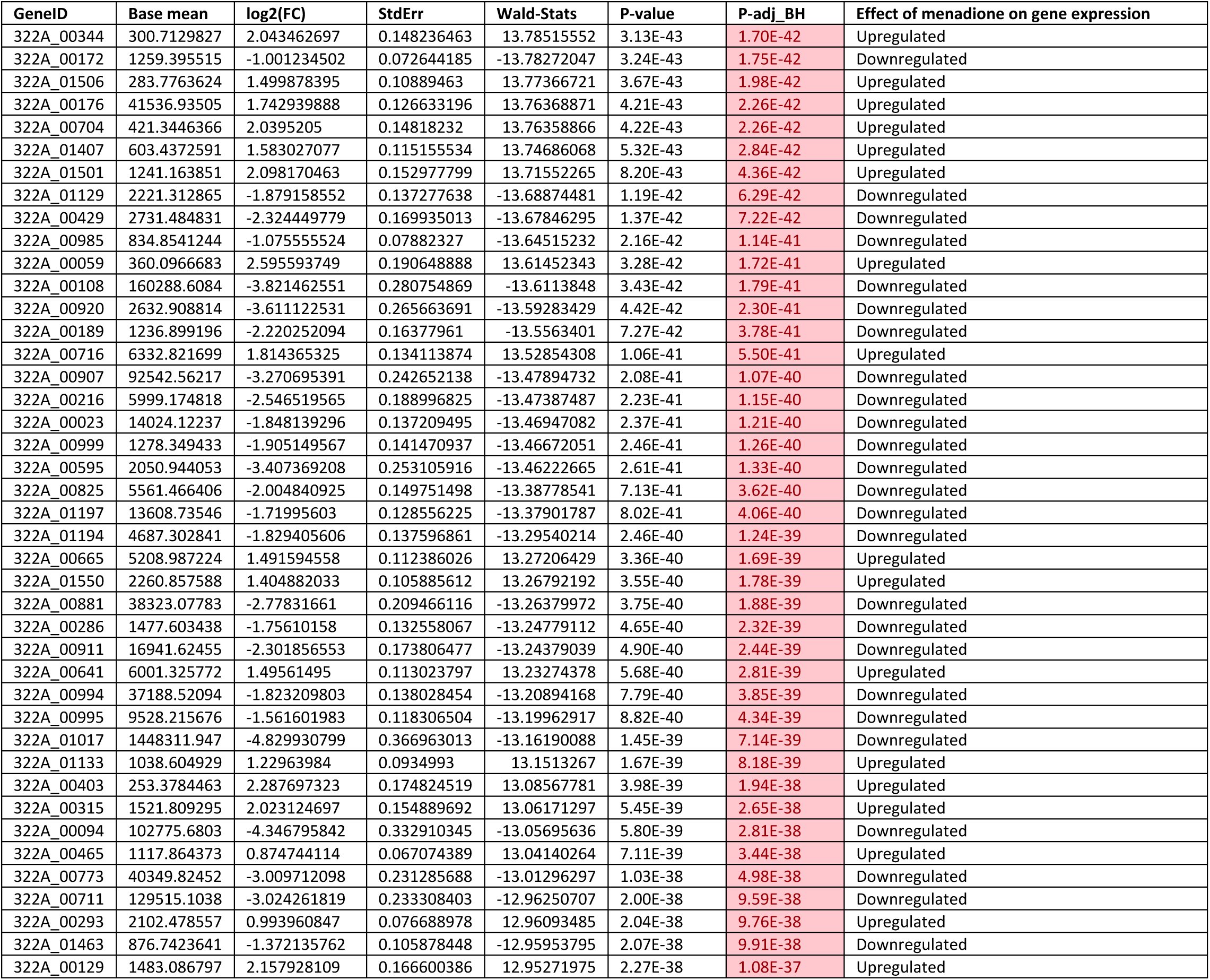
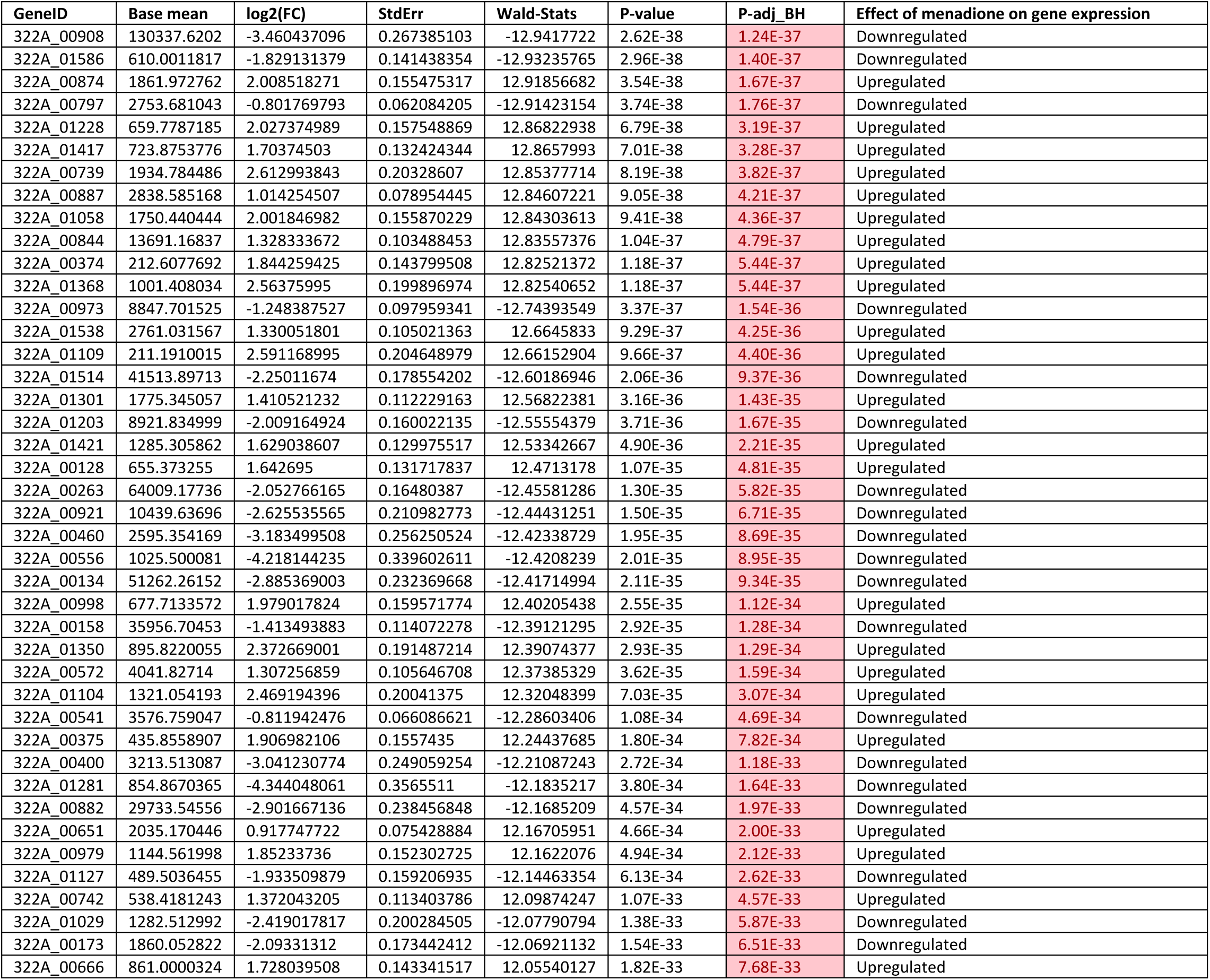
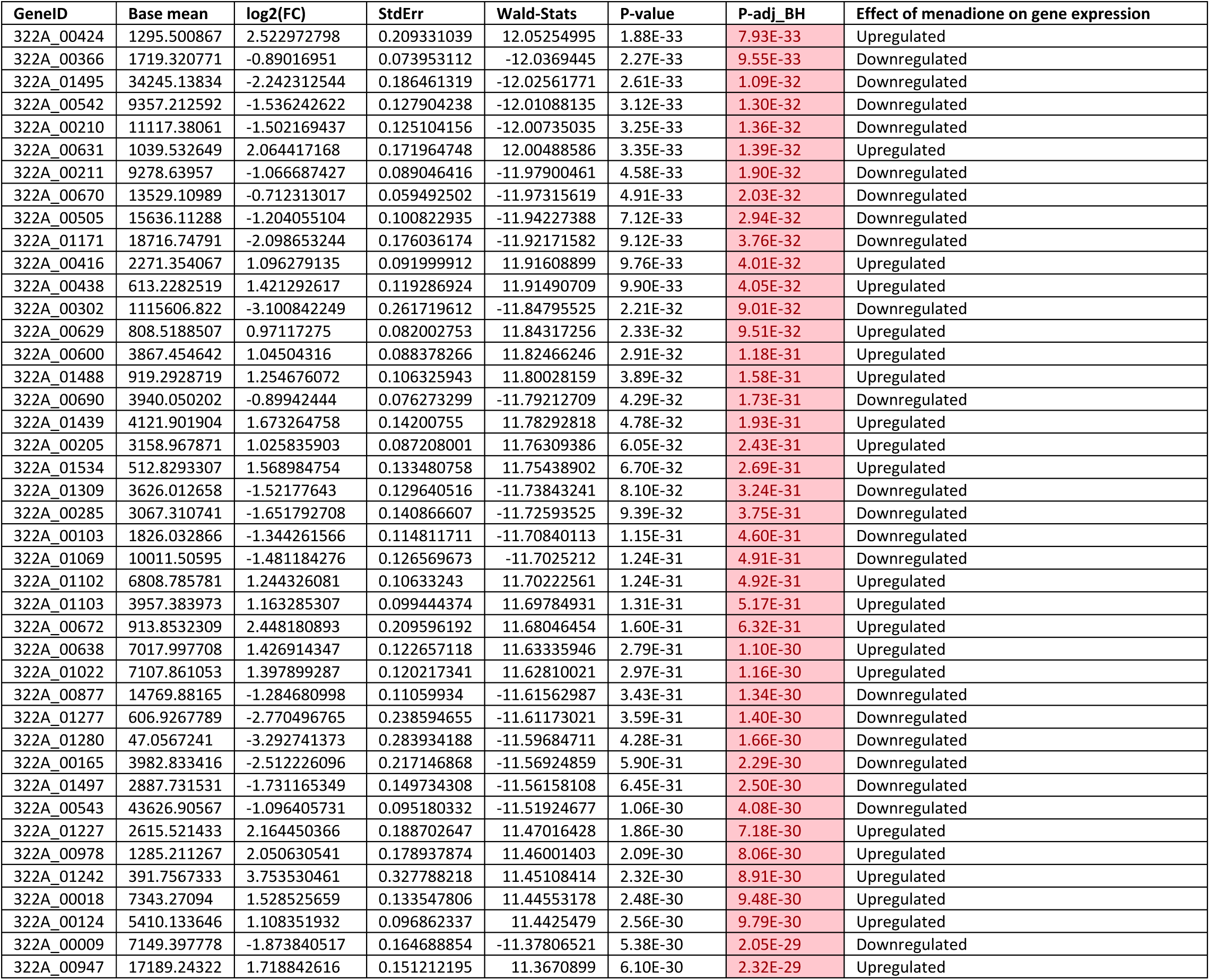
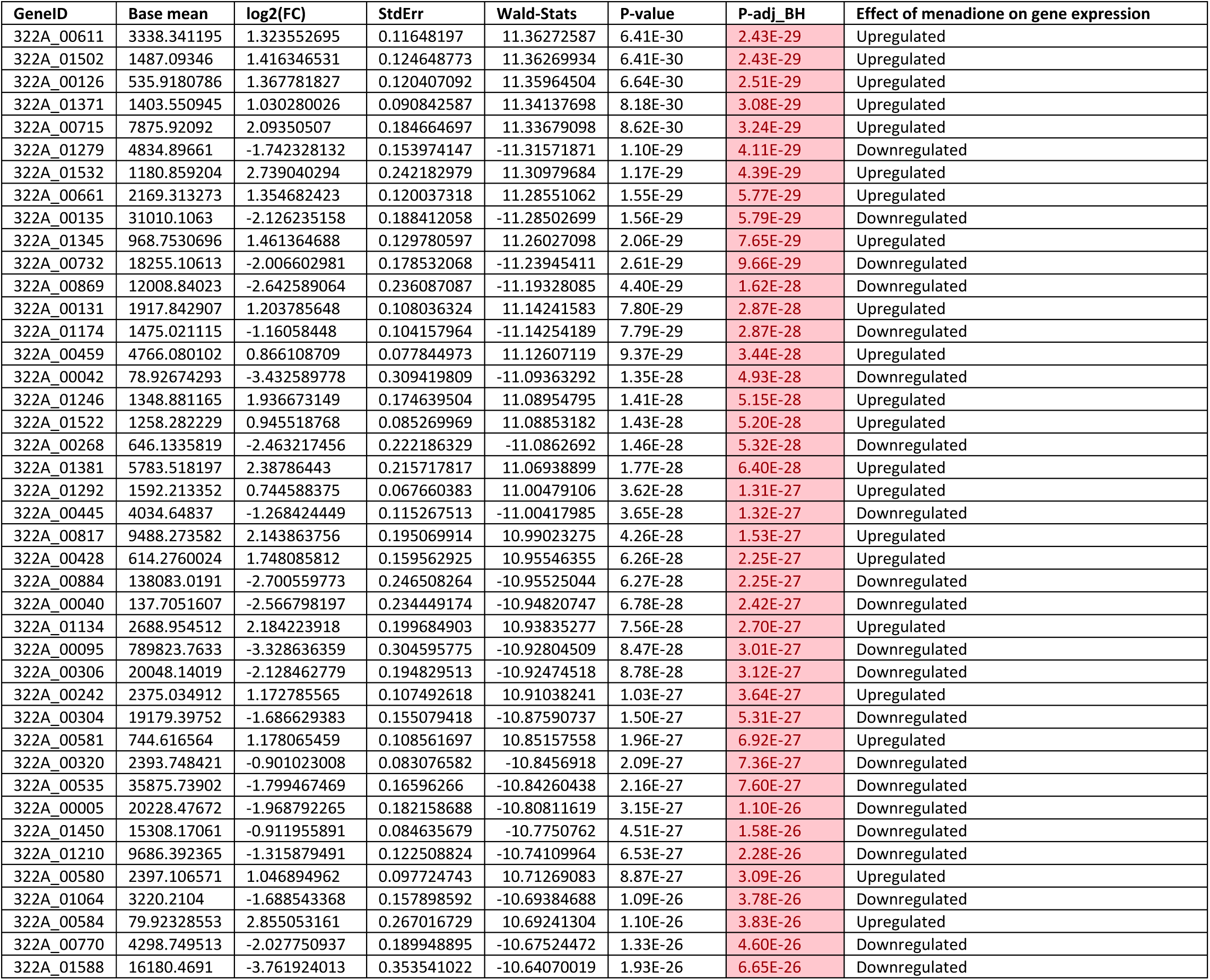
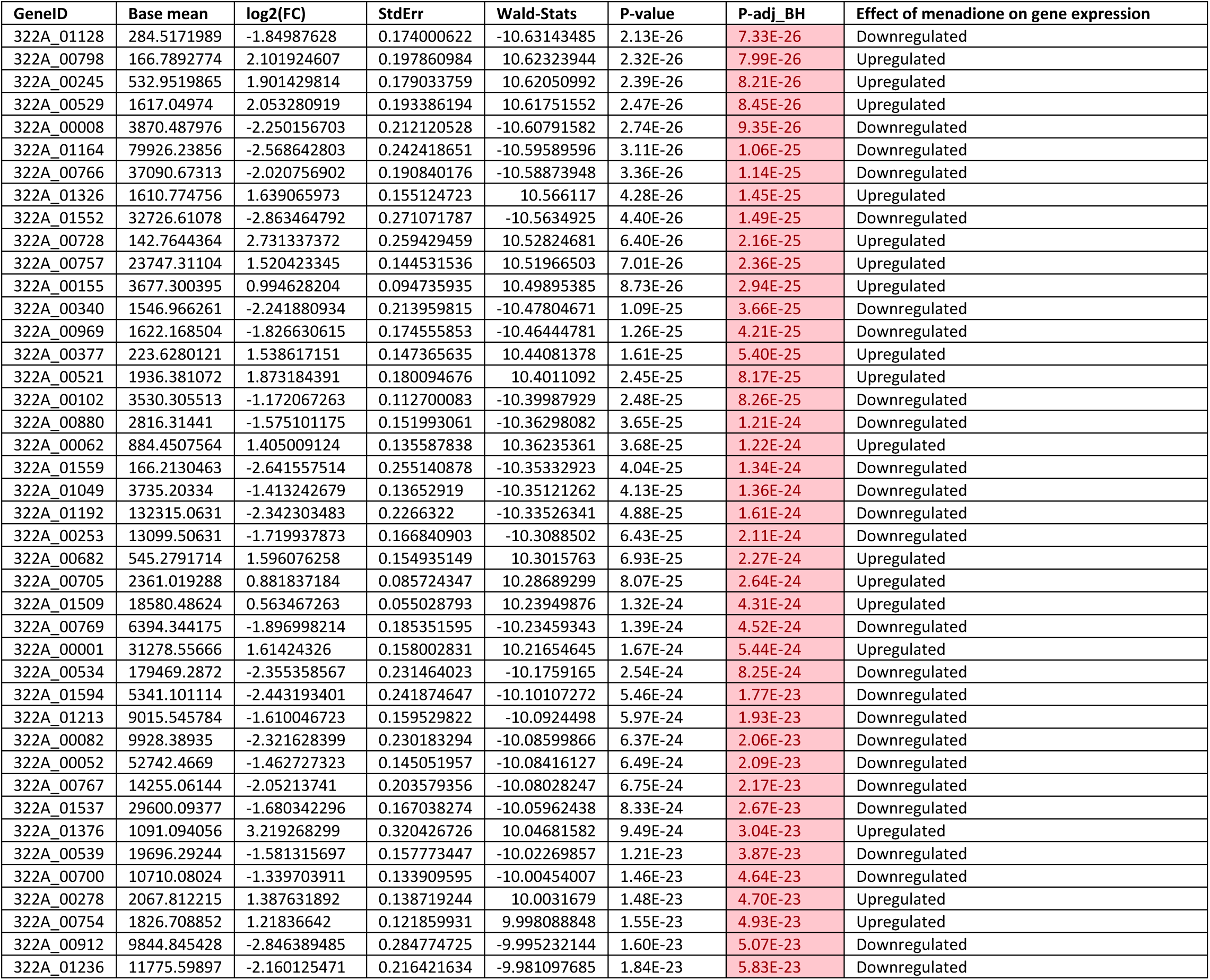
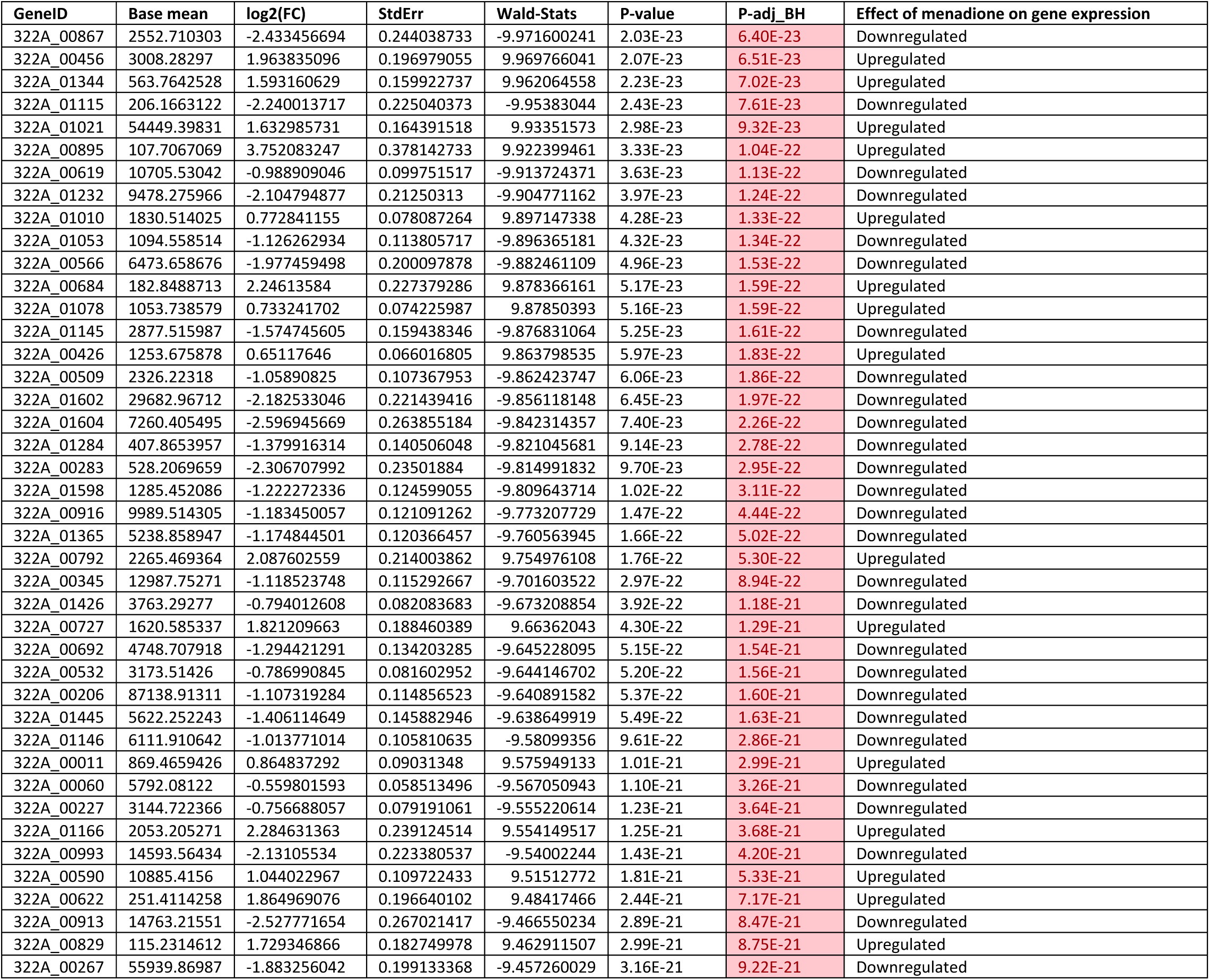
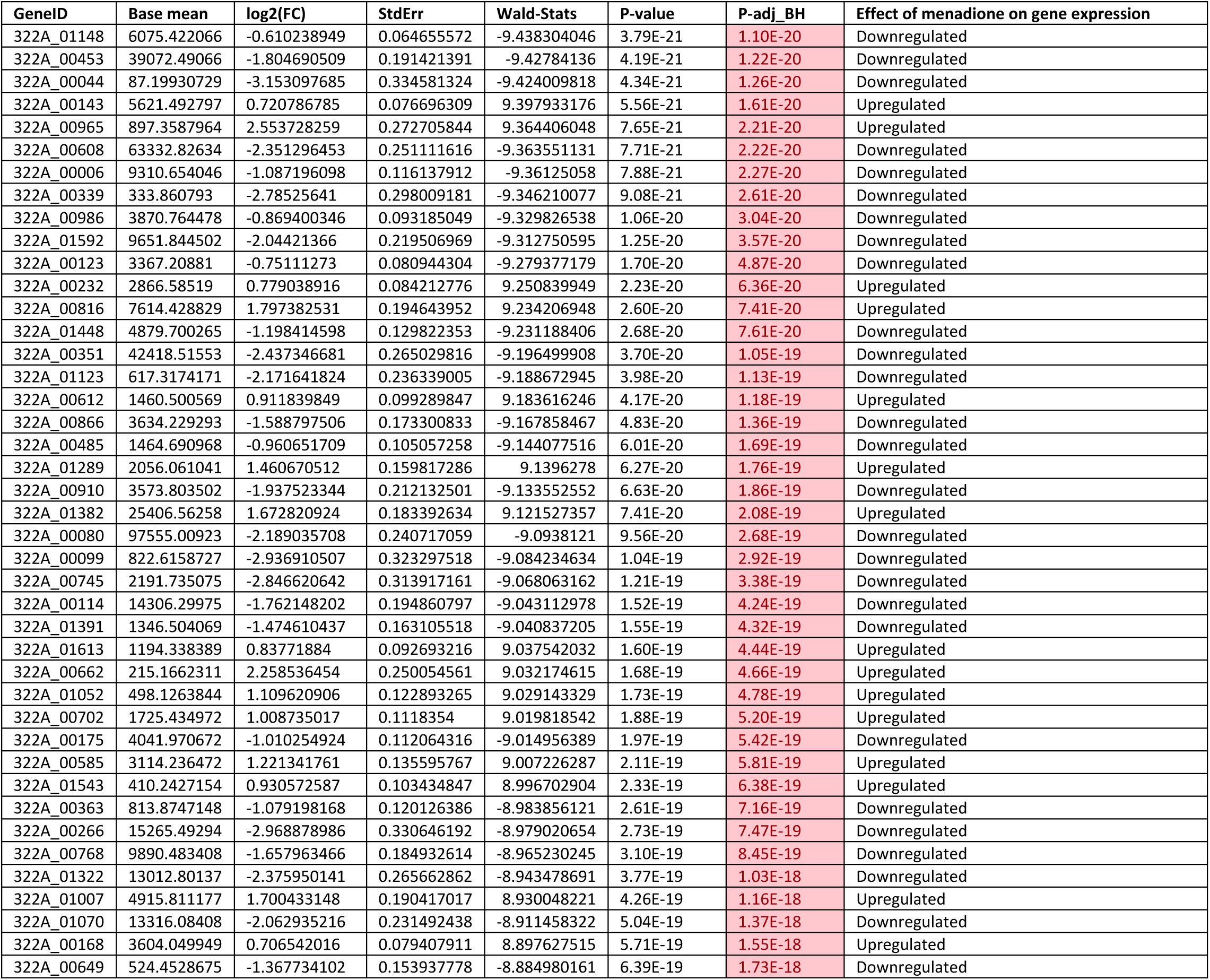
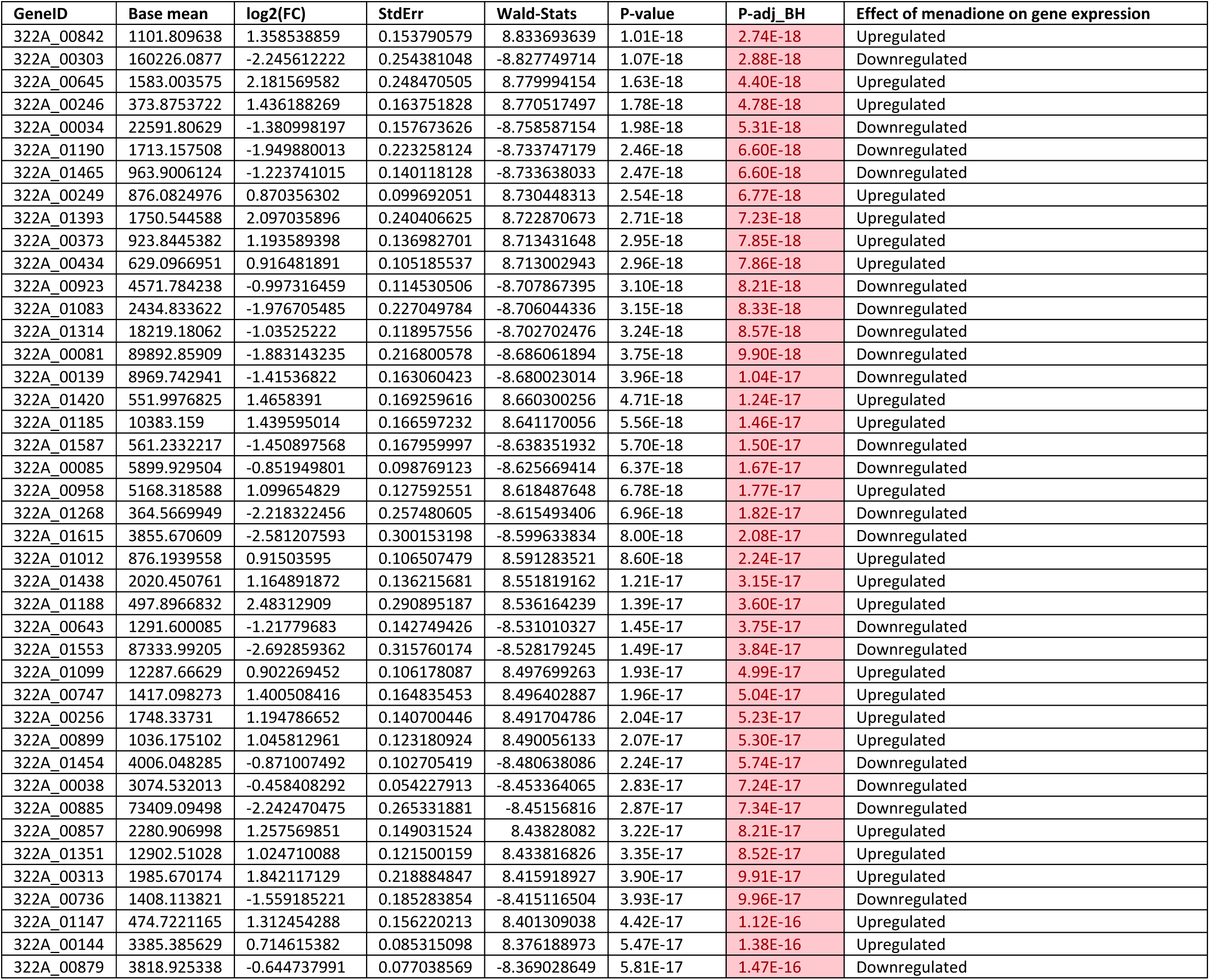
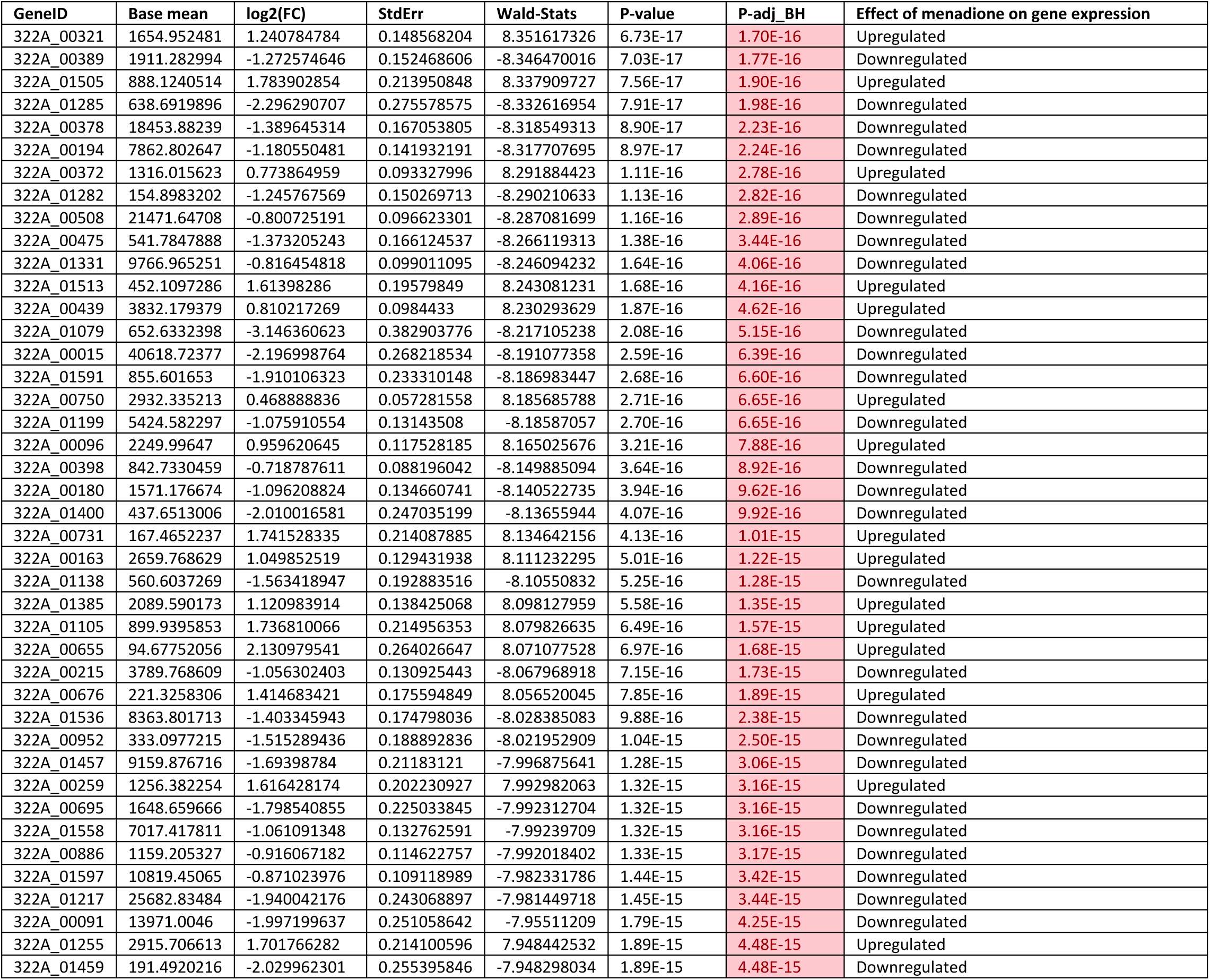
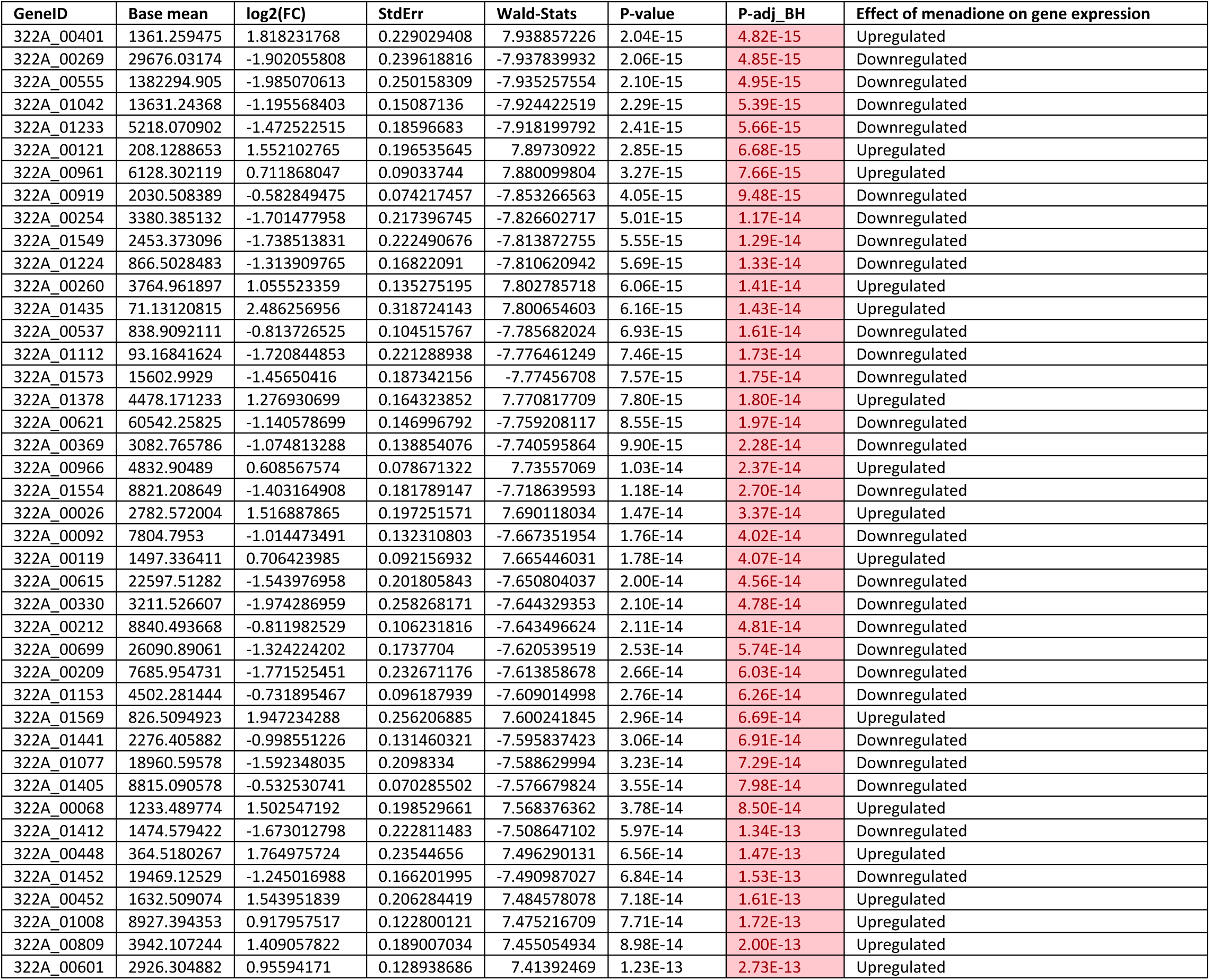
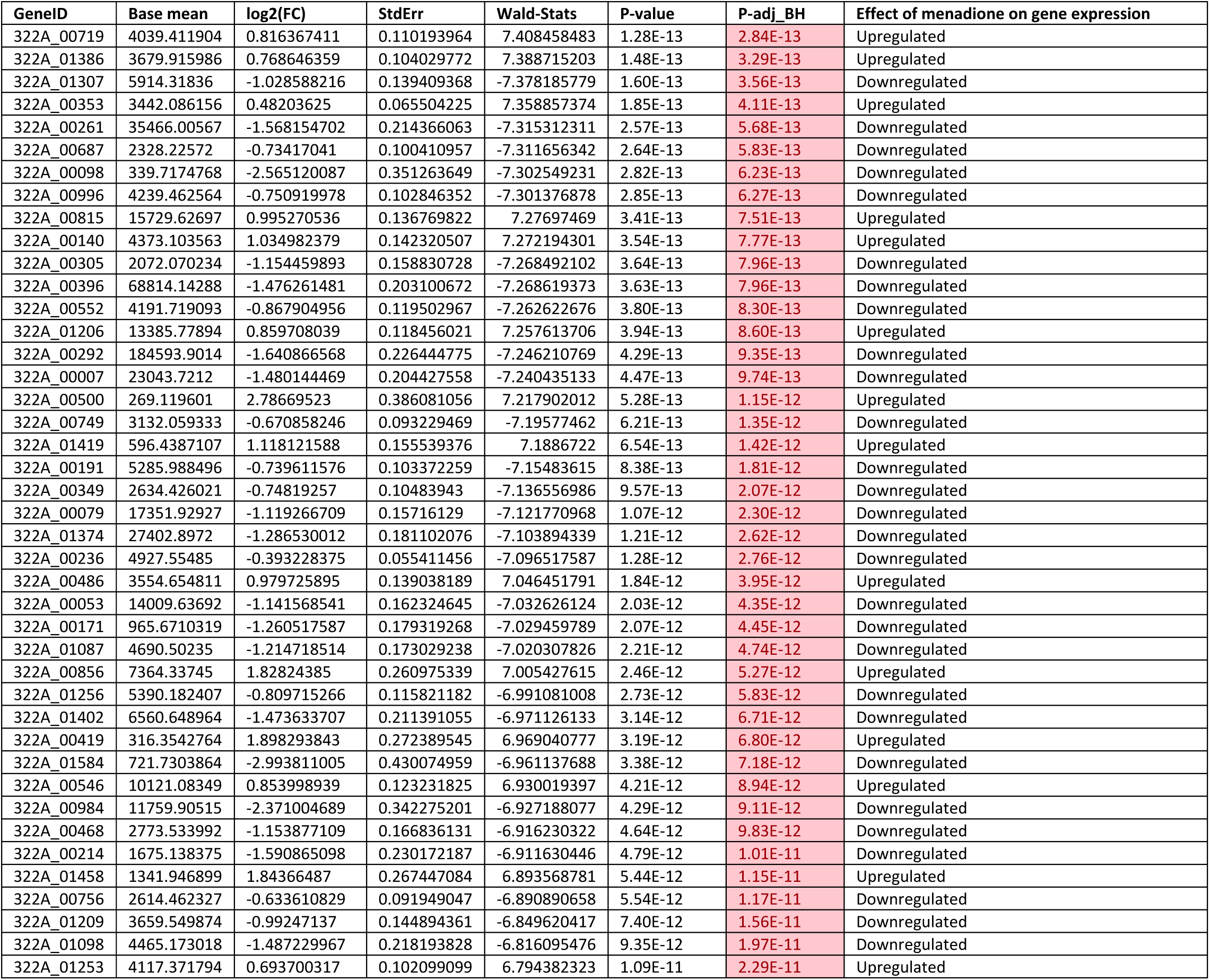
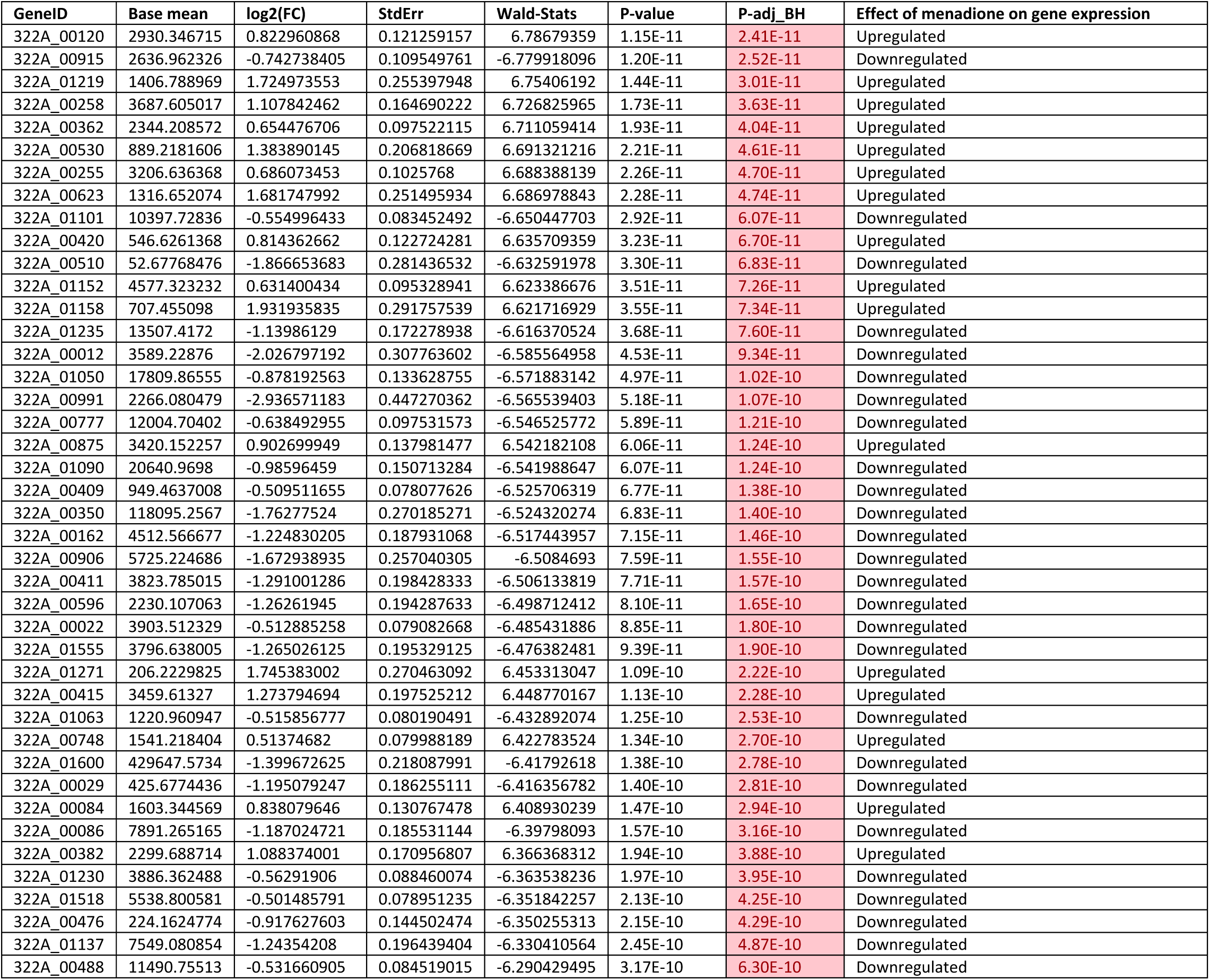
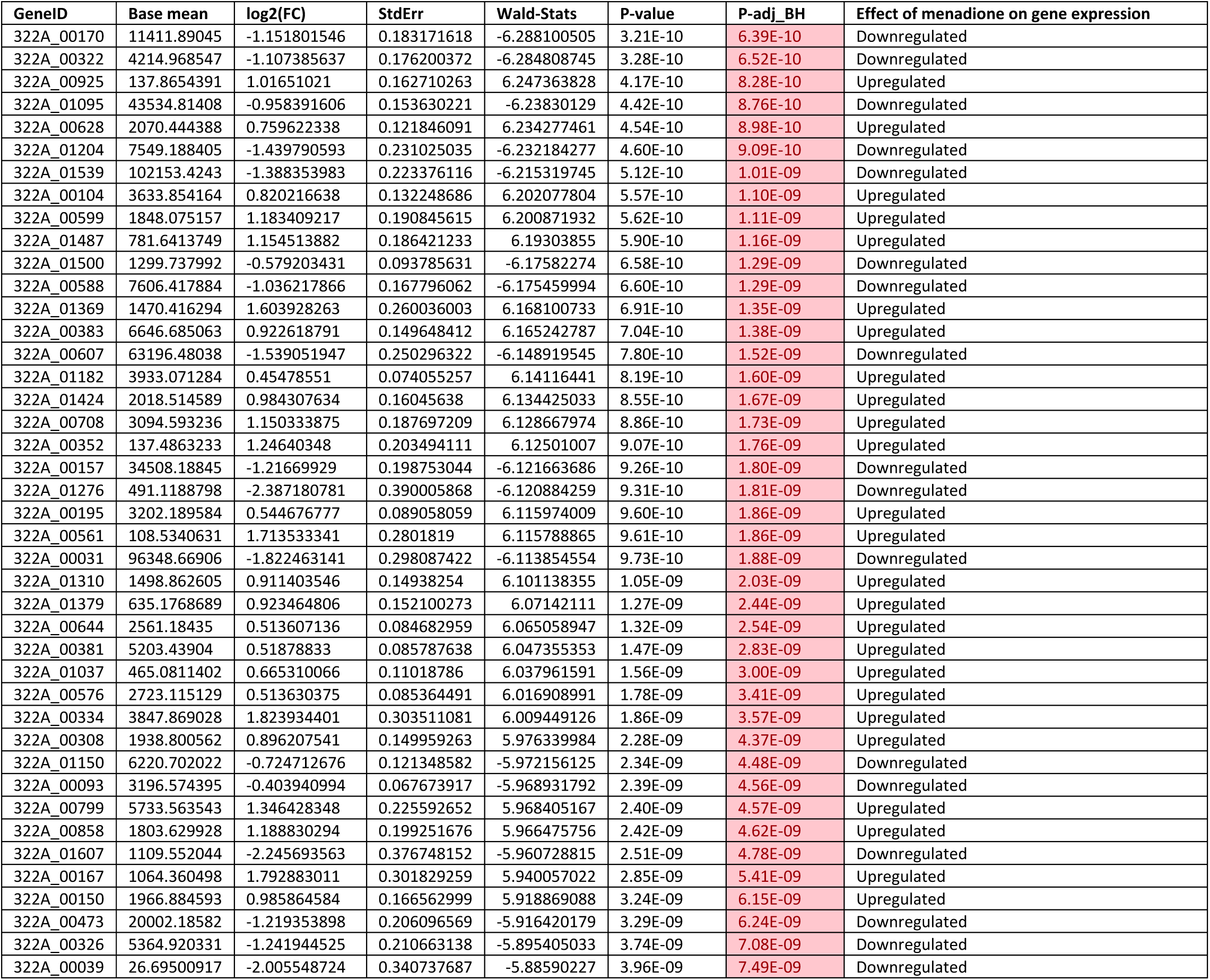
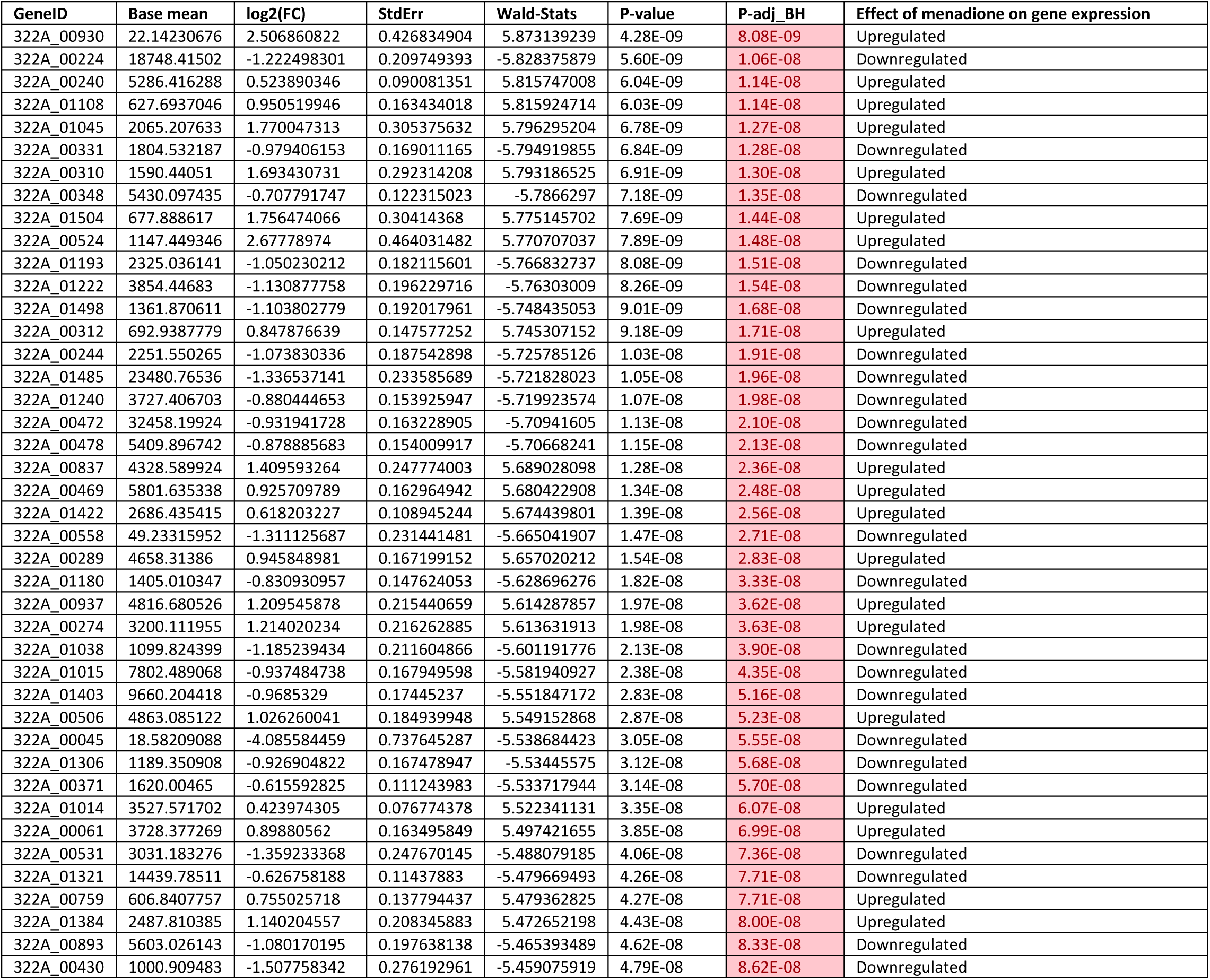
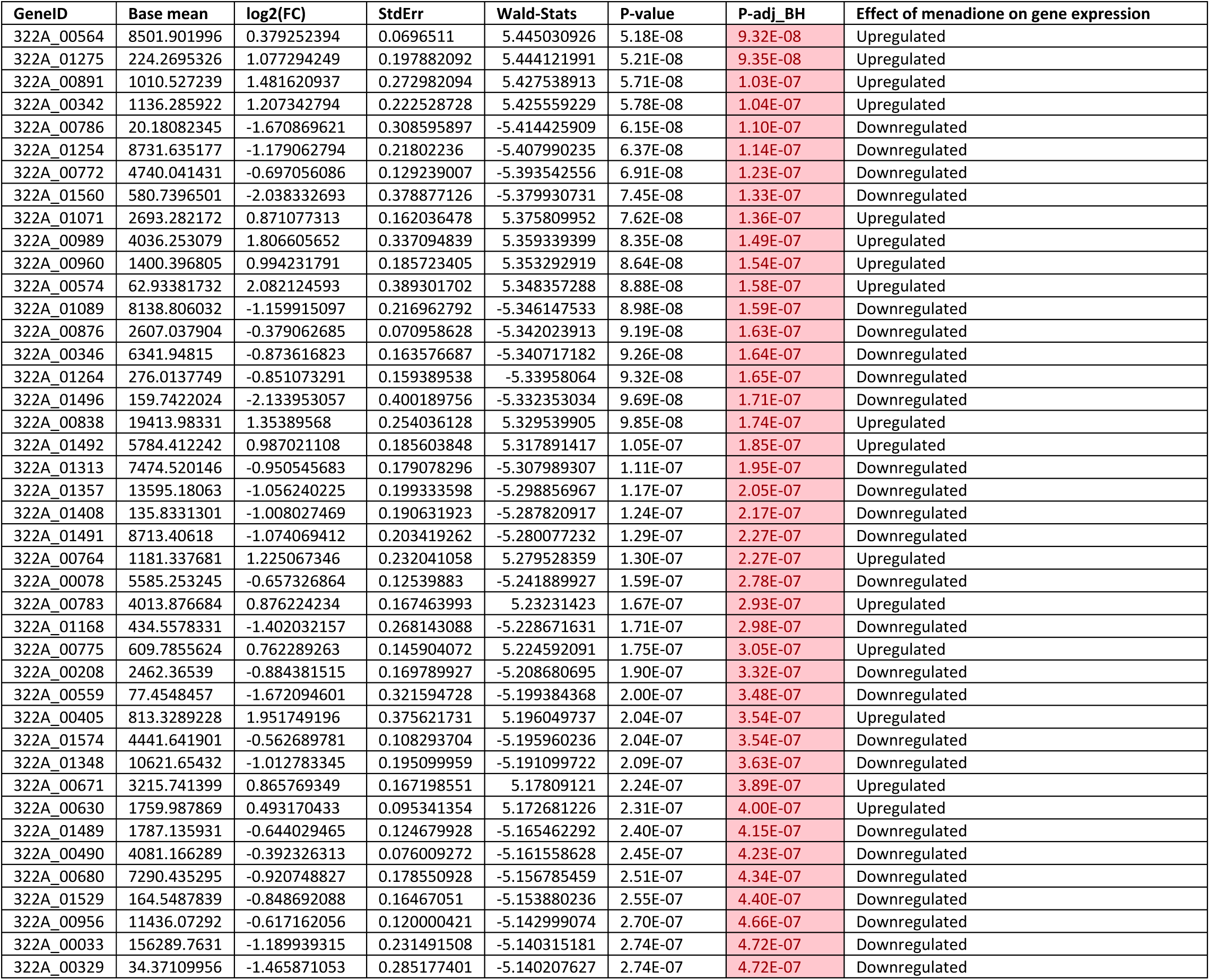
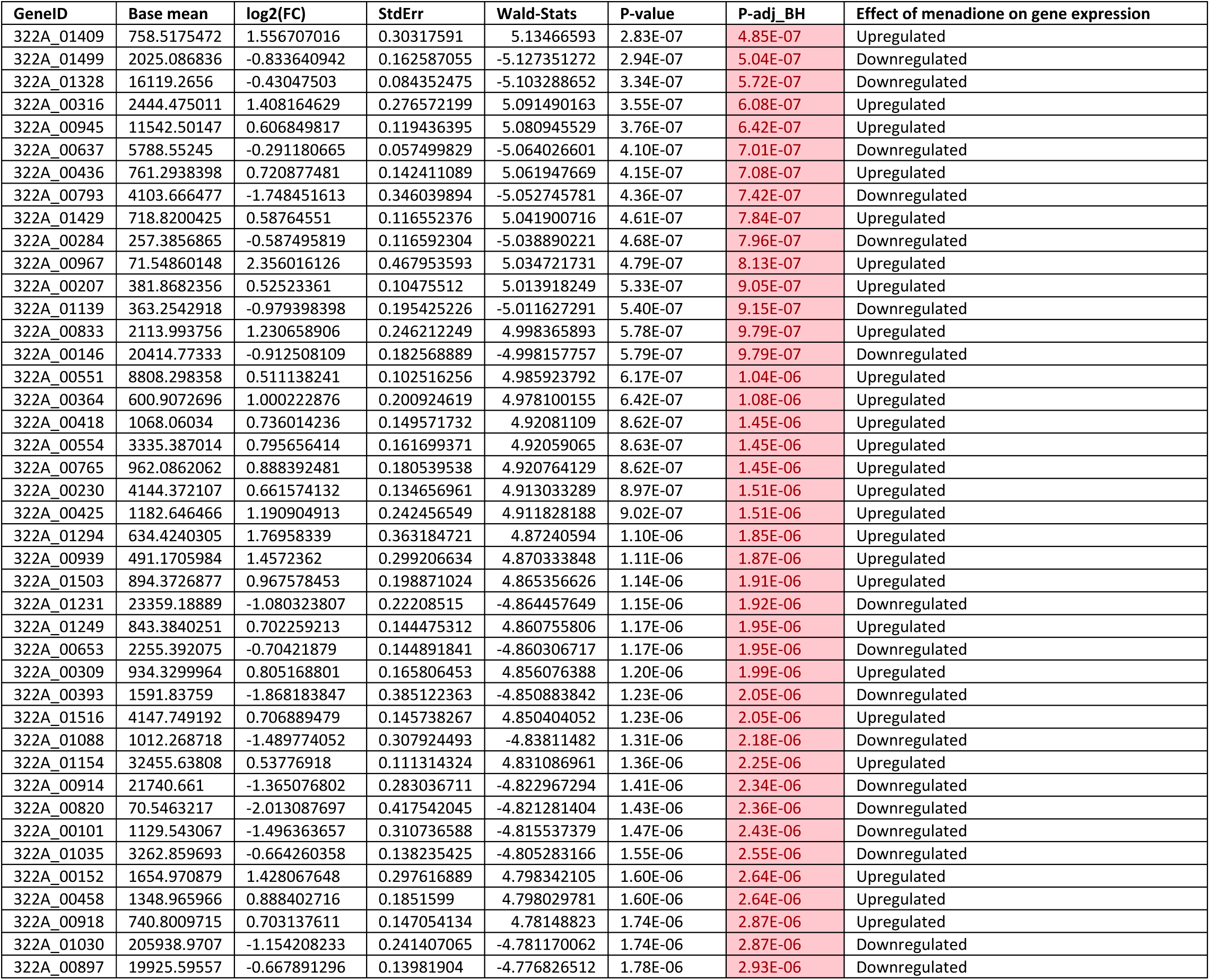
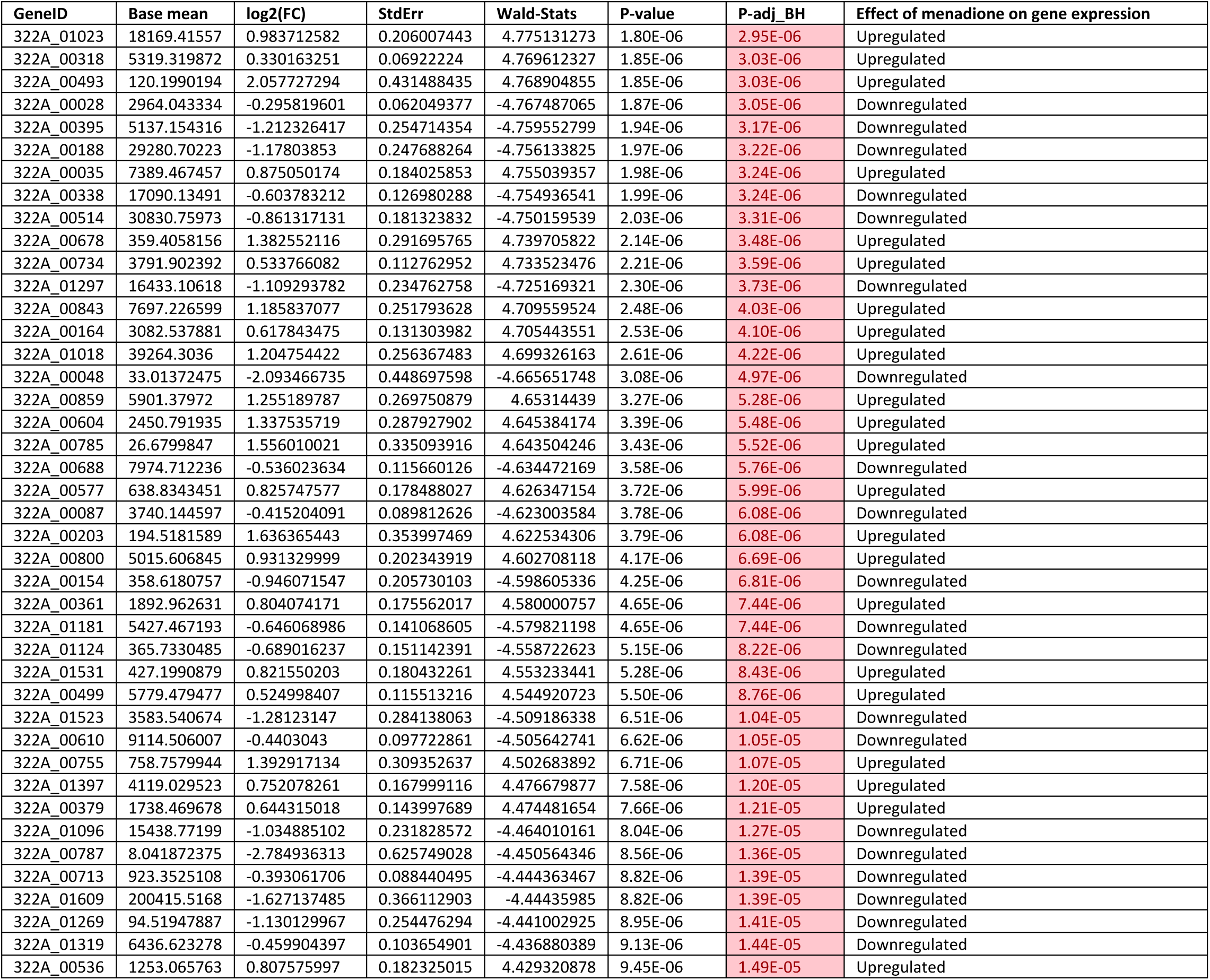
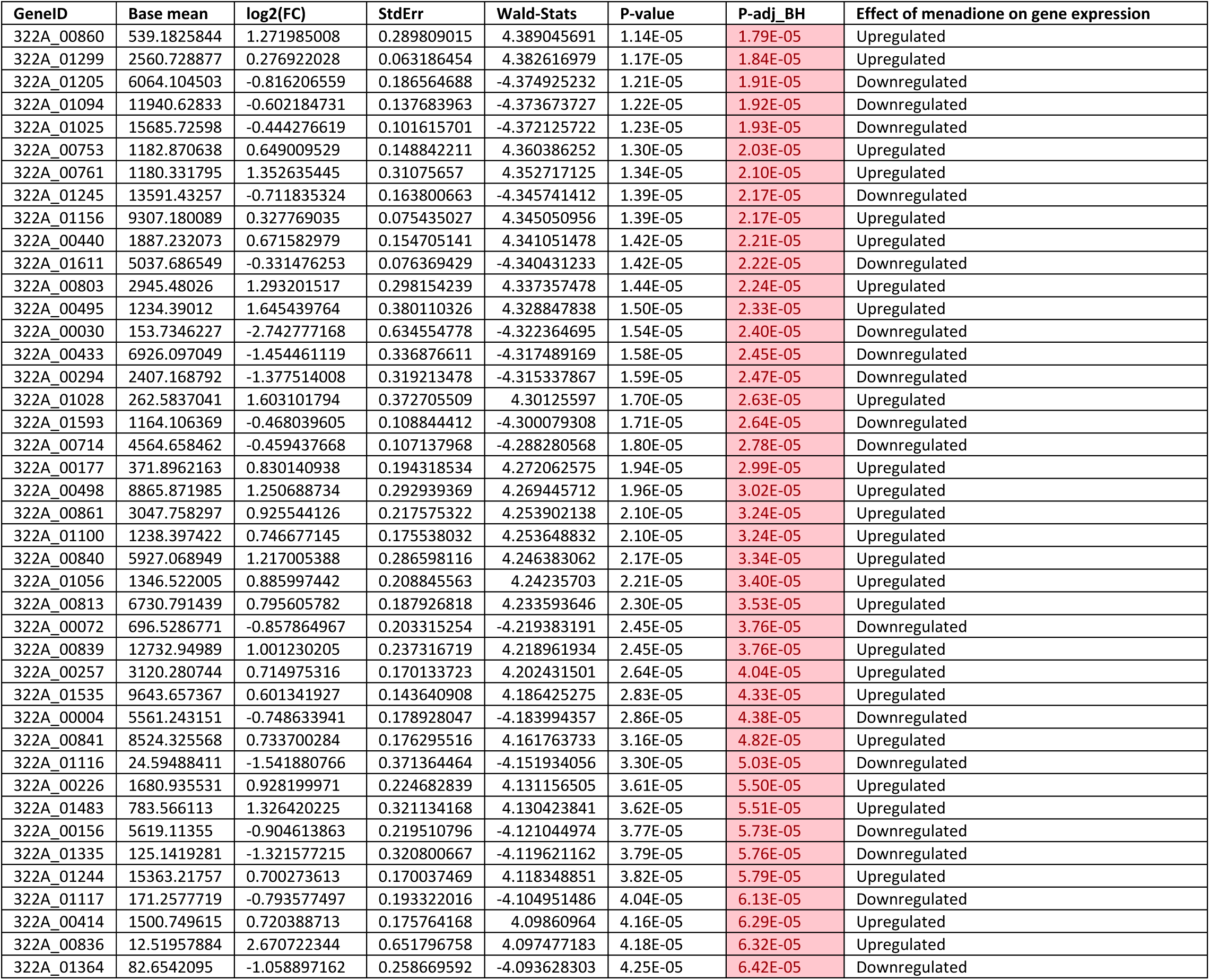
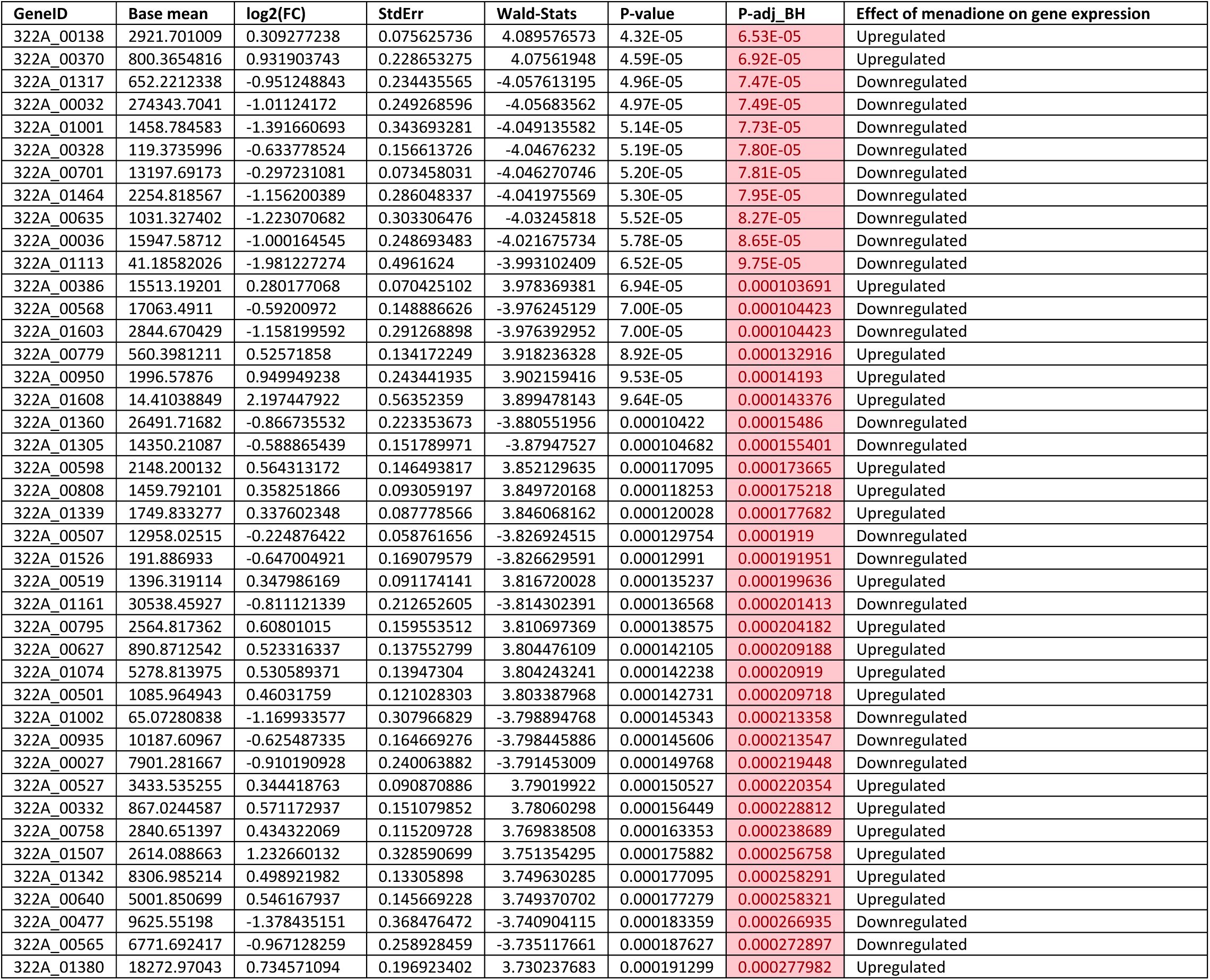
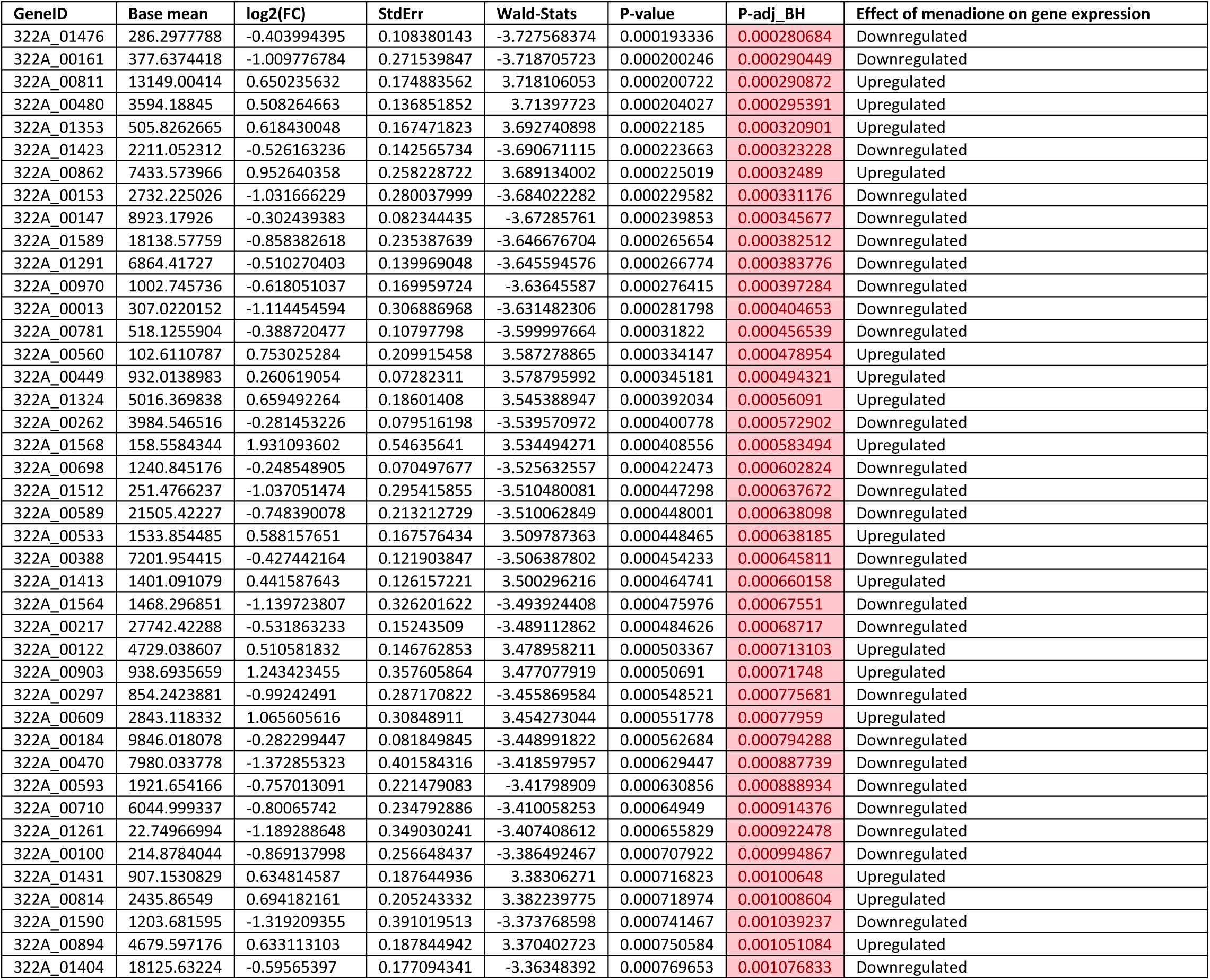
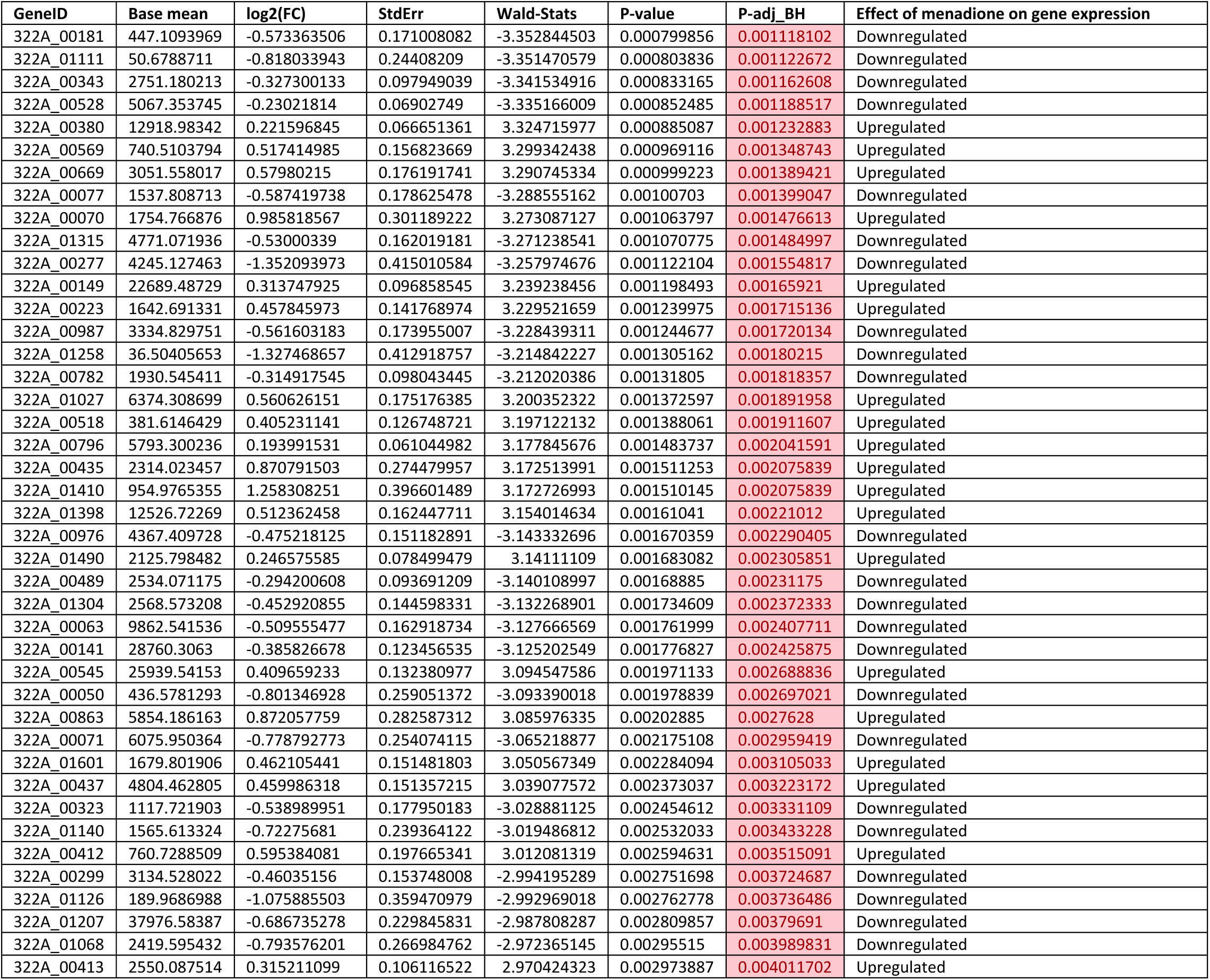
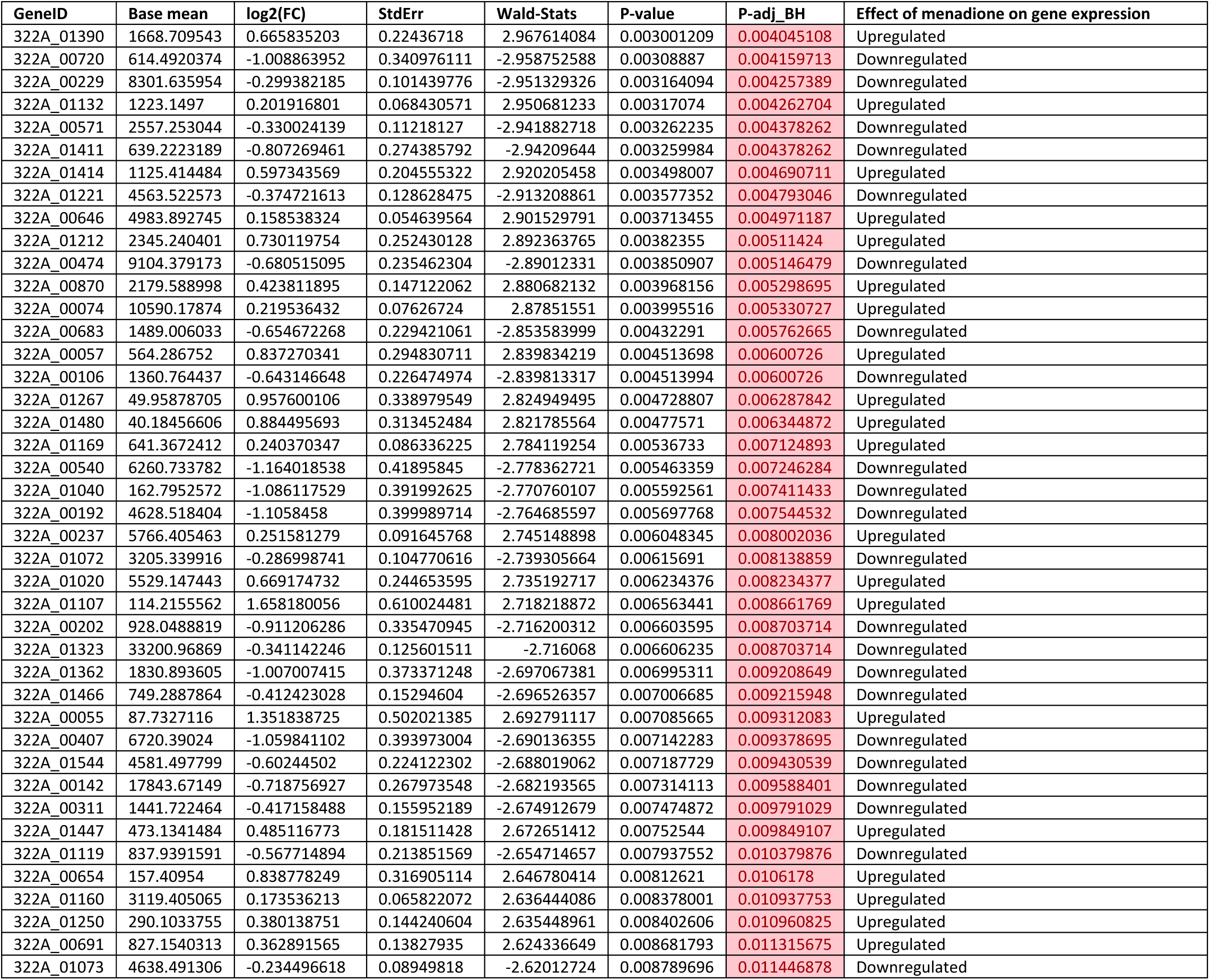
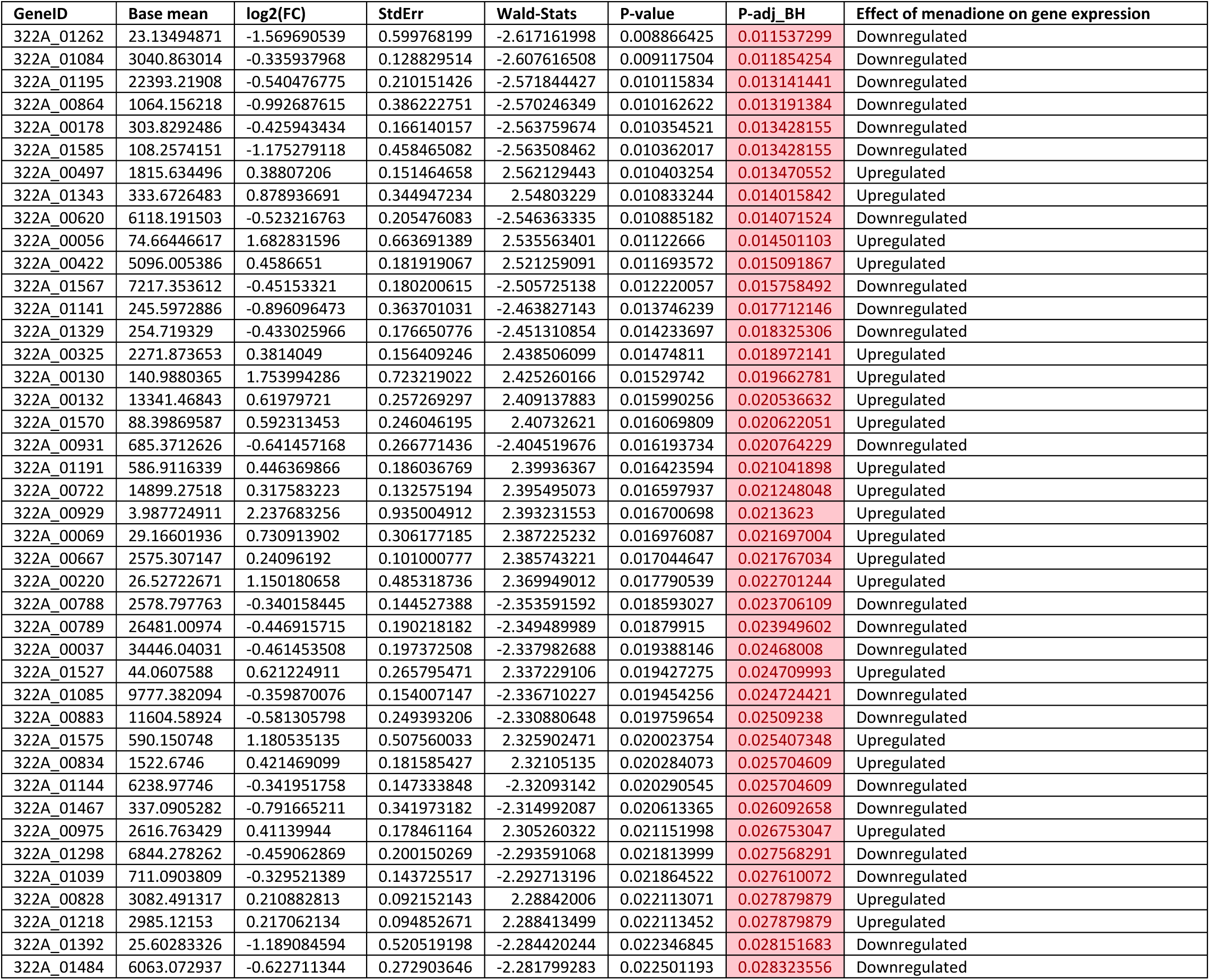
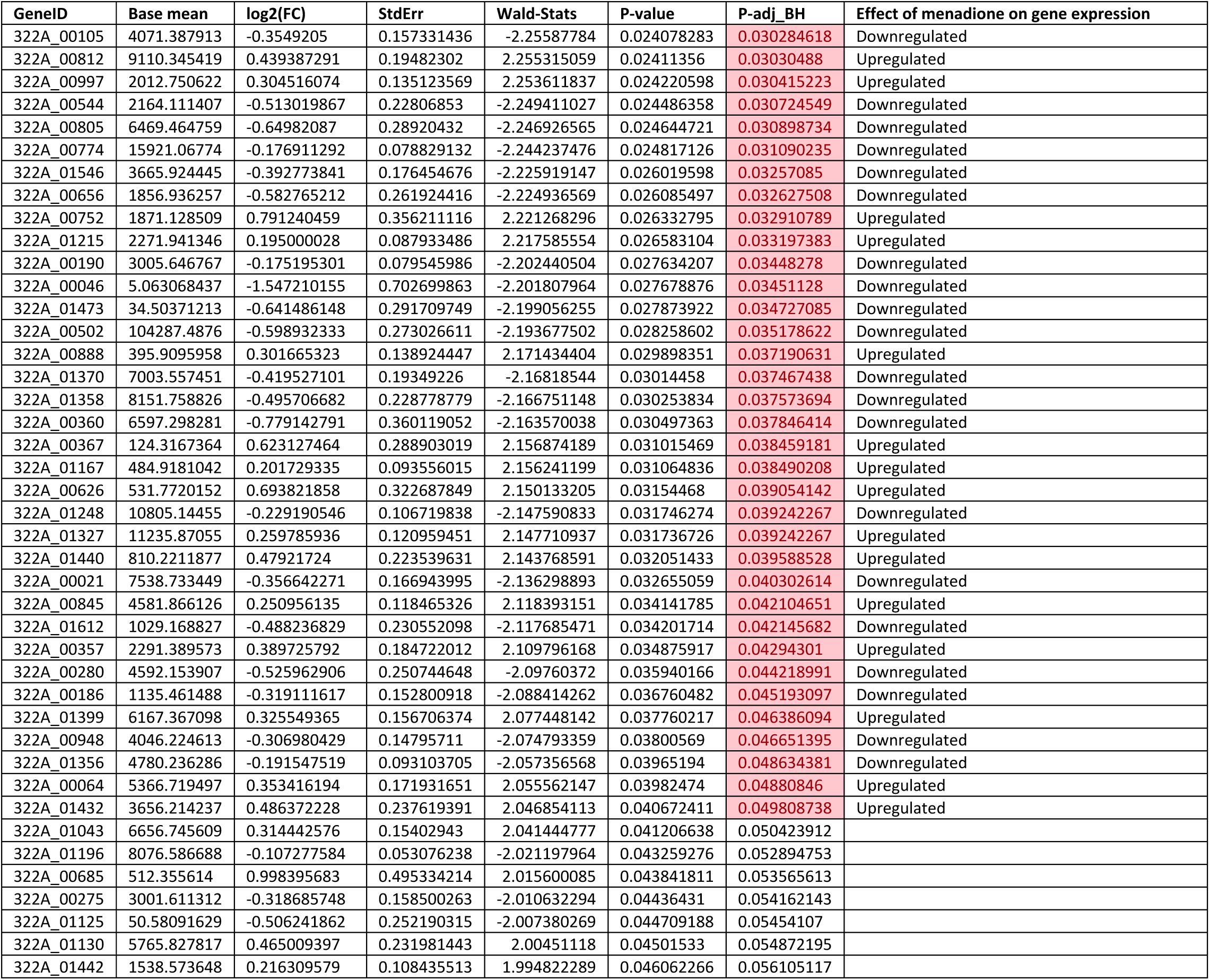
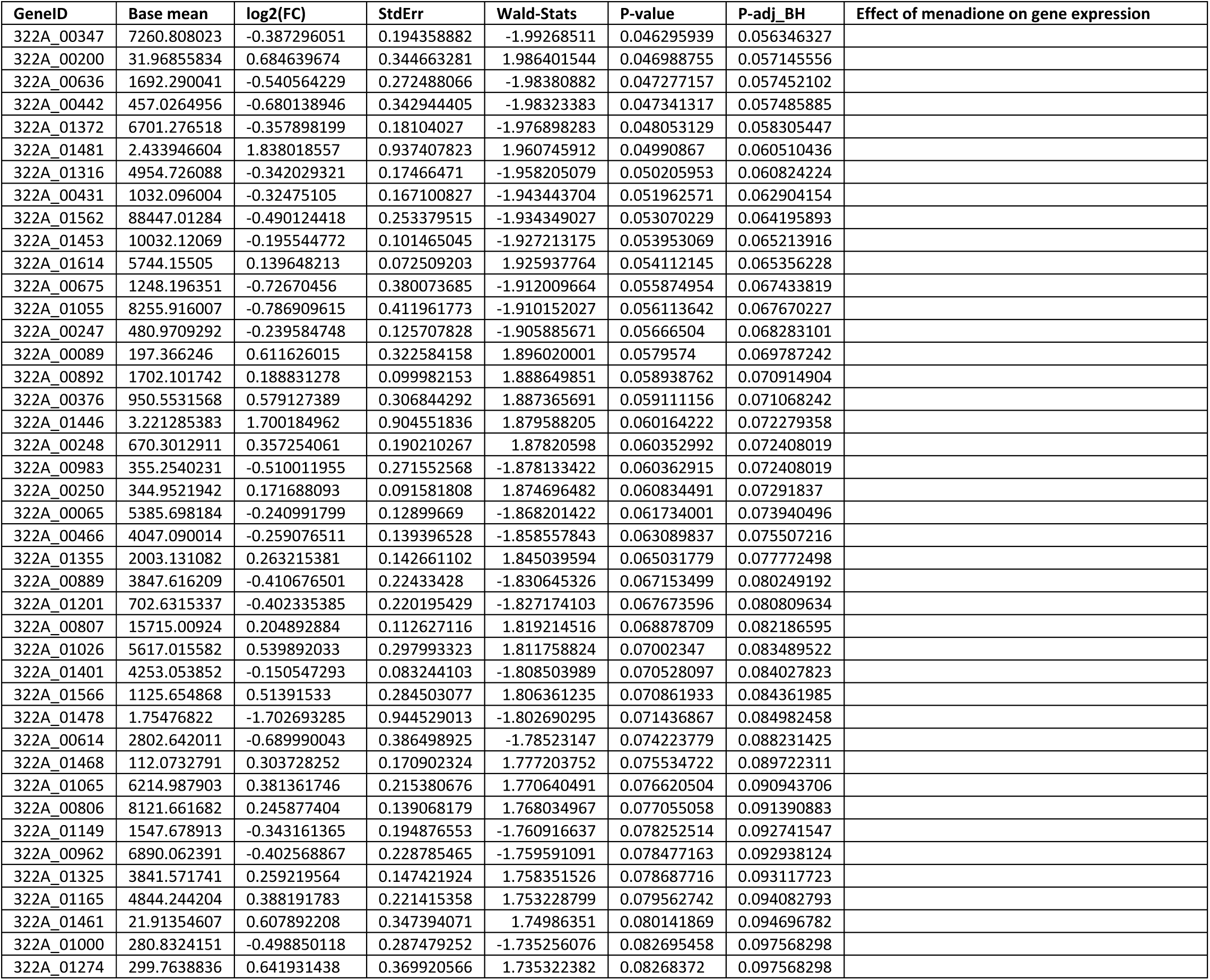
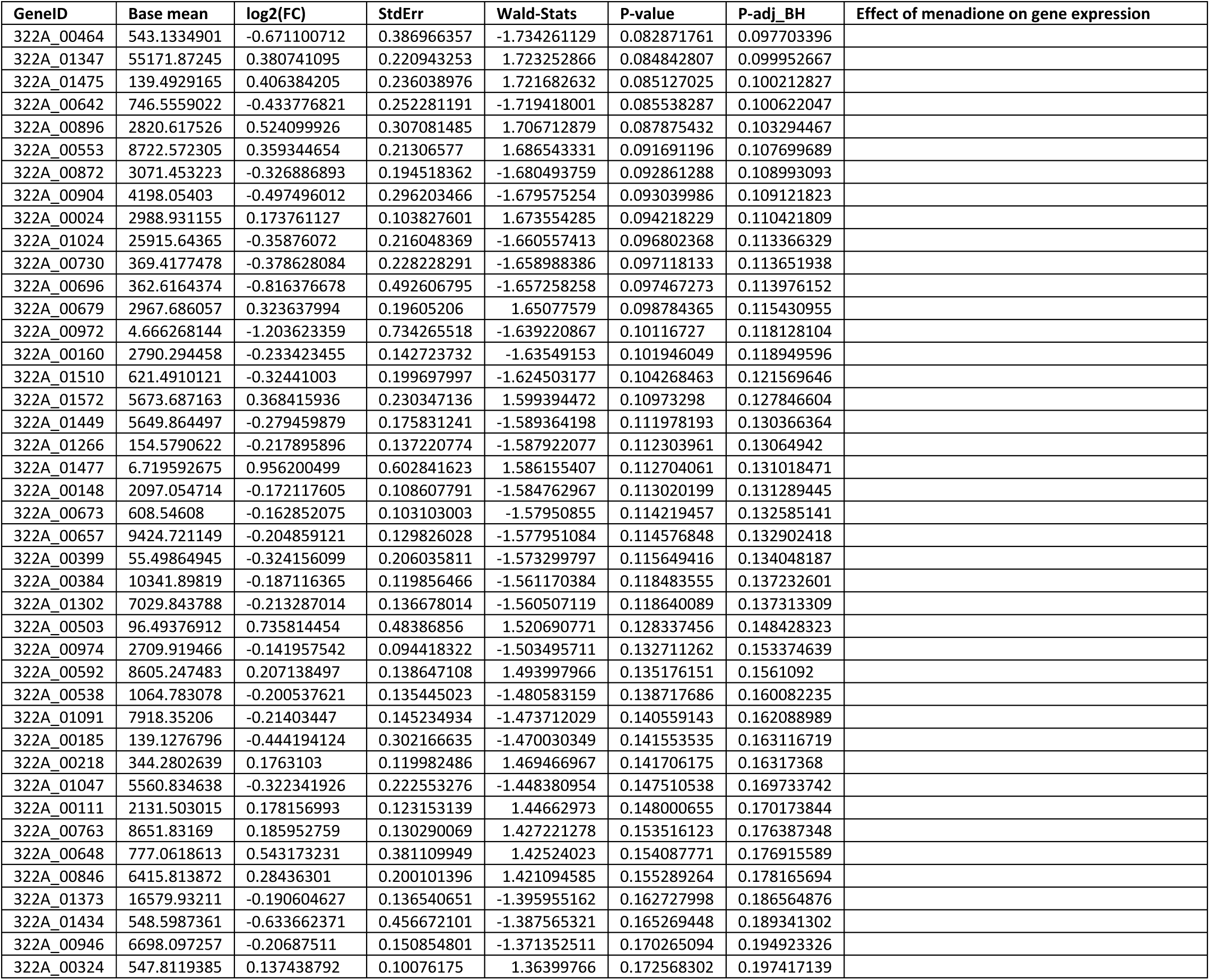
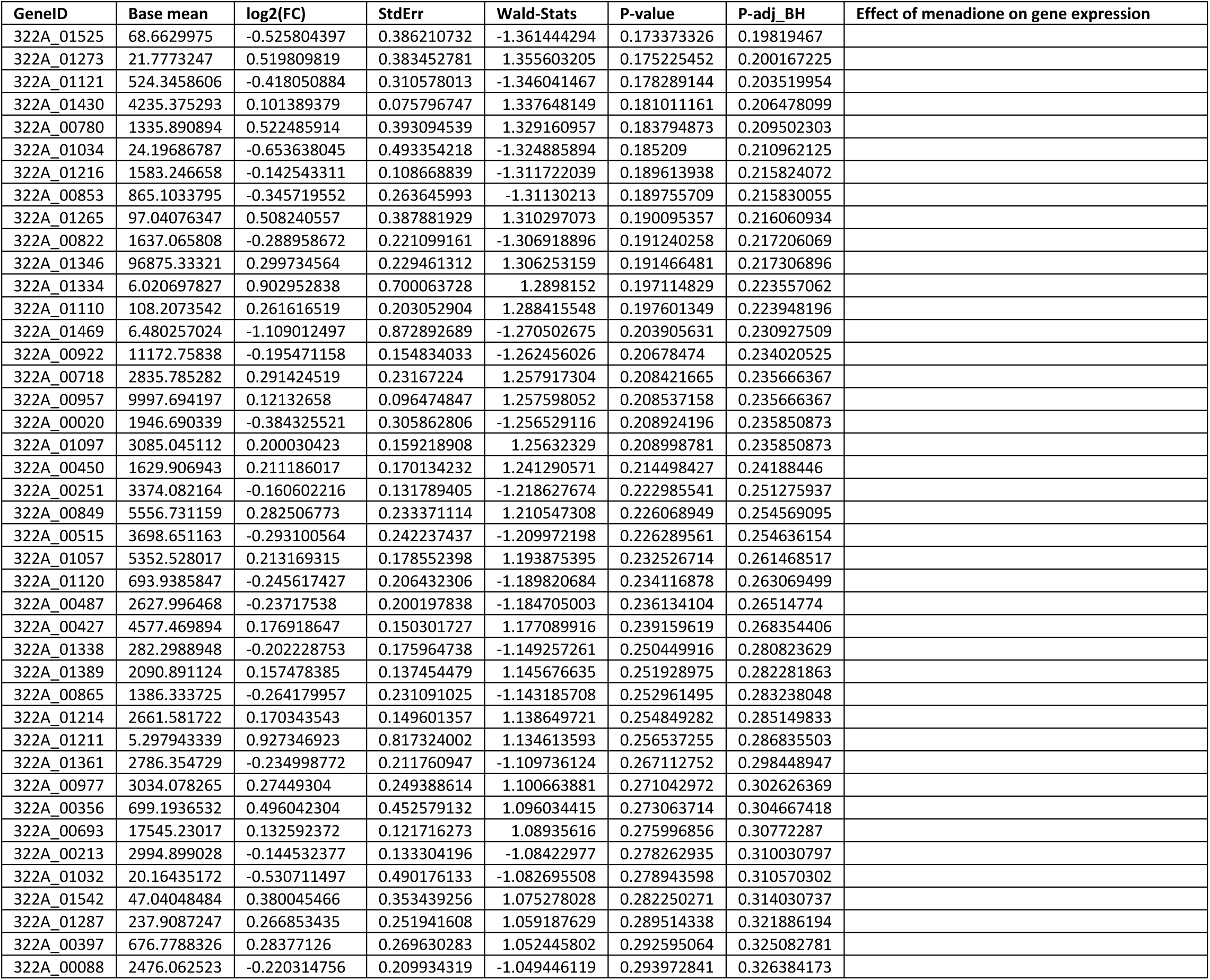
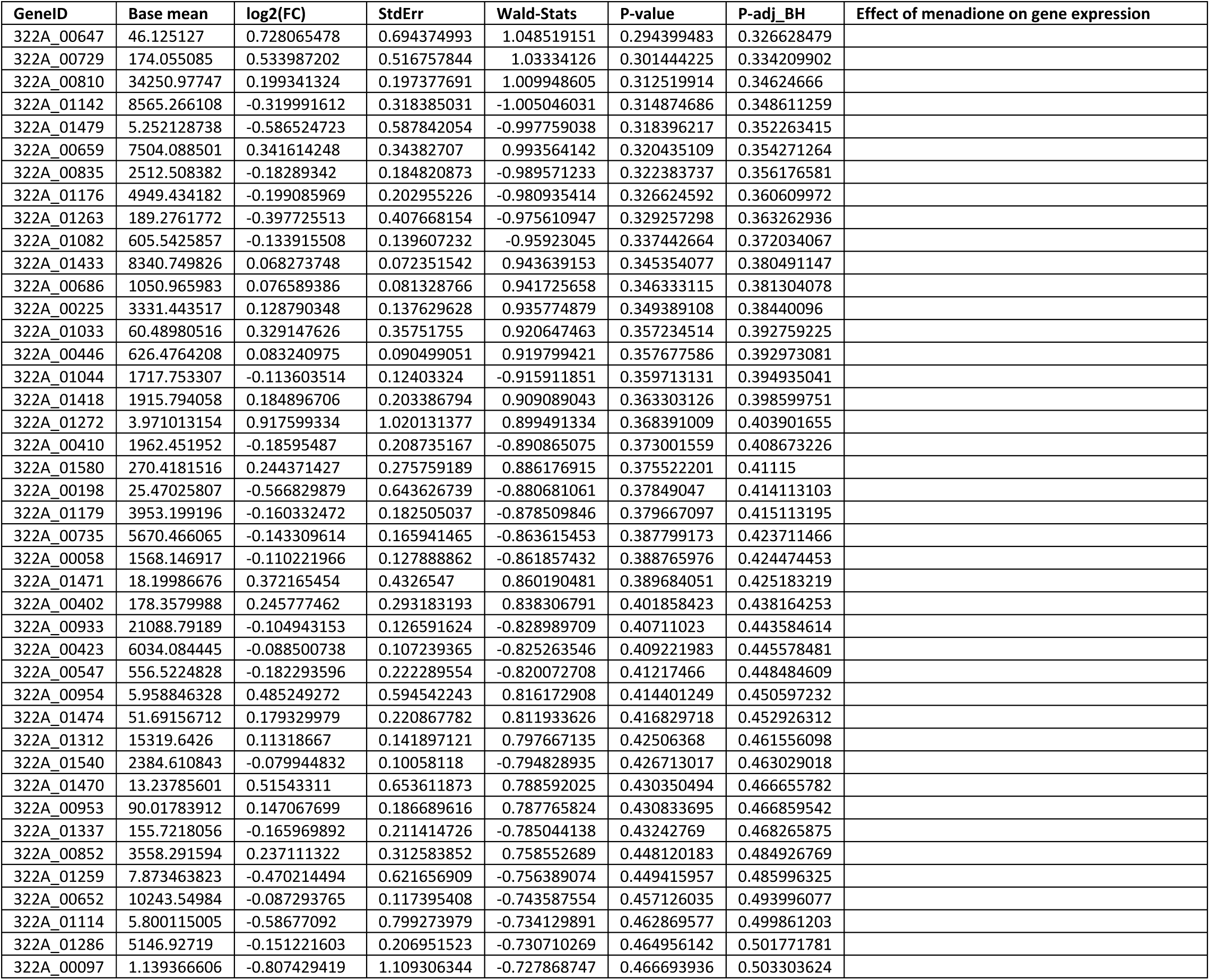
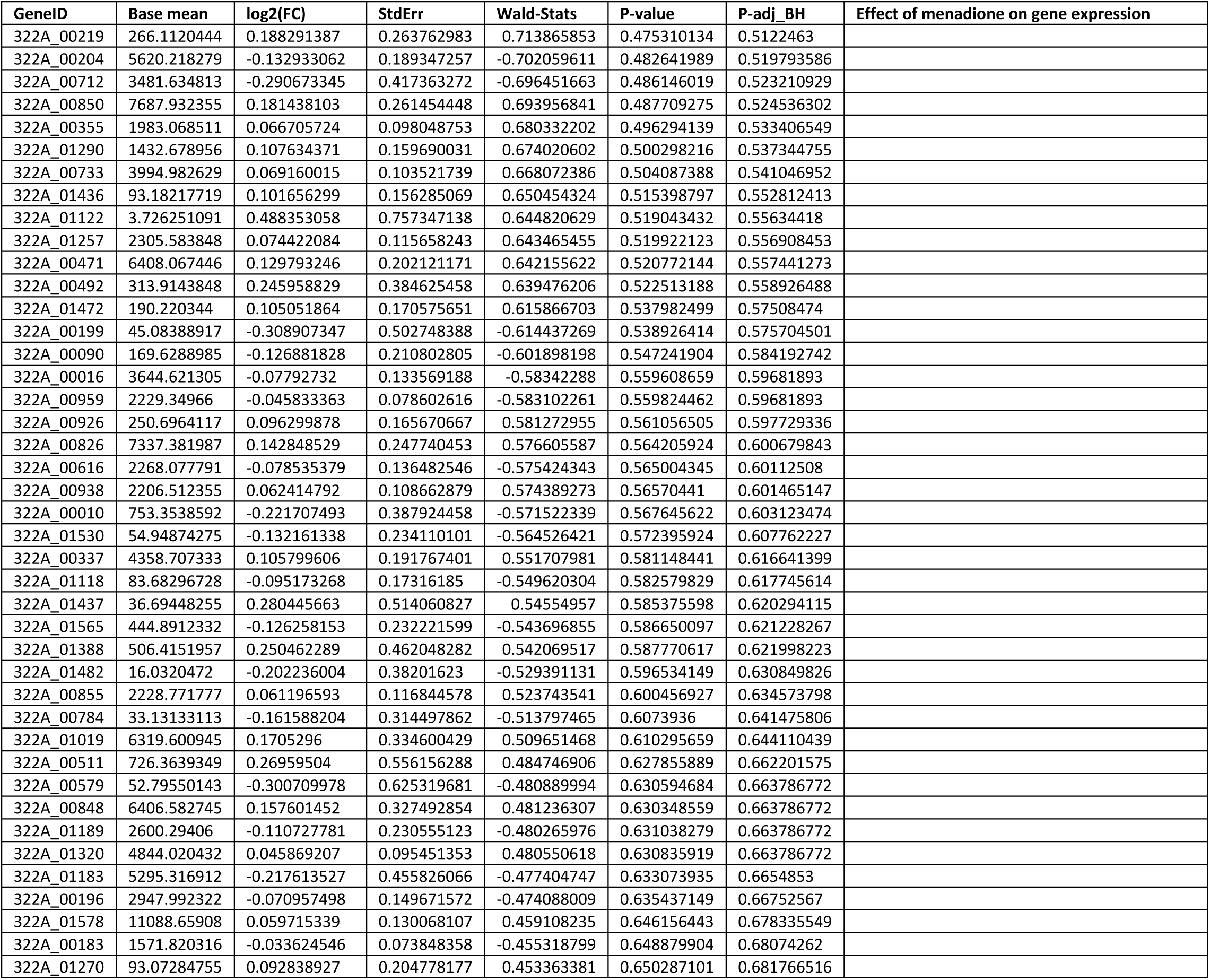
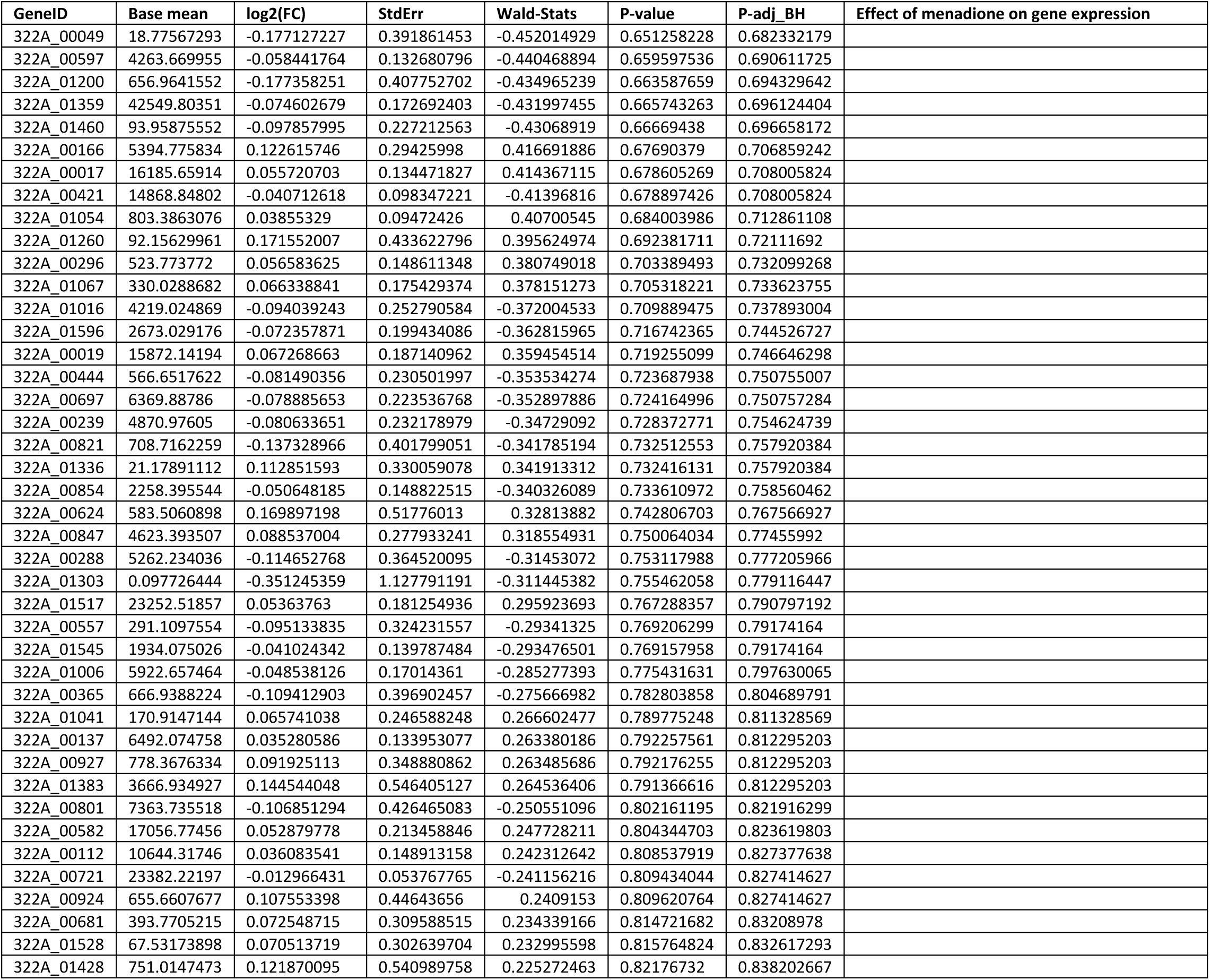
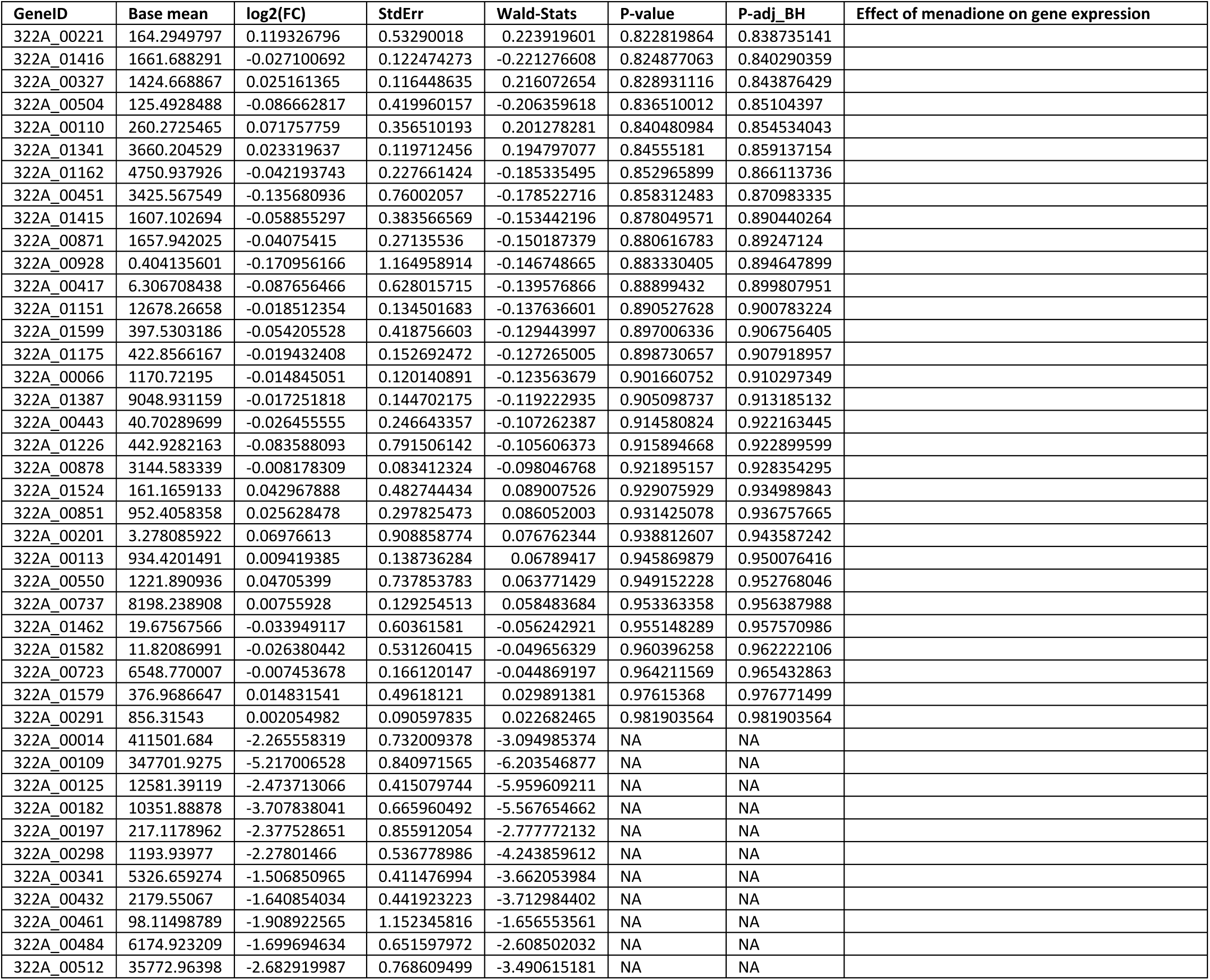
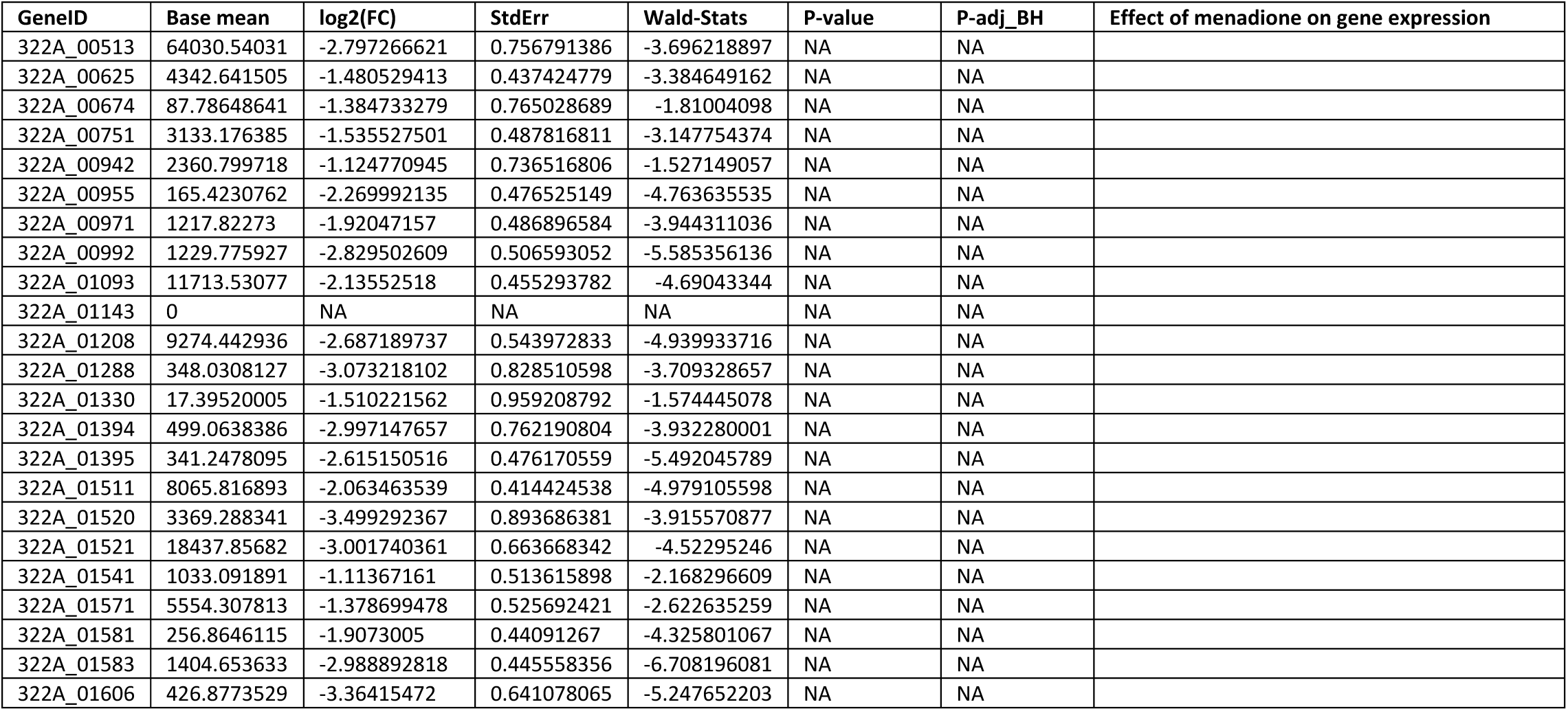
Outputs of DESeq2 analysis, used to identify significantly differentially expressed genes.

**Supplementary Table 3.** Results from Signalling Pathway Impact Analysis of *H. pylori* 322A’s significantly differentially expressed genes mapped to the genome of *H. pylori* 26695.

| Name | ID | pSize | NDE | pNDE | tA | pPERT | pG | pGFdr | pGFWER | Status | KEGGLINK |
| --- | --- | --- | --- | --- | --- | --- | --- | --- | --- | --- | --- |
| Bacterial chemotaxis | 2030 | 16 | 15 | 0.143055 | -3.91292 | 0.485 | 0.254501 | 0.391228 | 0.763503 | Inhibited | http://www.genome.jp/dbget-bin/show_pathway?hpy02030+HP_0816+HP_0815+HP_0584+HP_1030+HP_1031+HP_0352+HP_1067+HP_0392+HP_0298+HP_0019+HP_0393+HP_0616+HP_0082+HP_0099+HP_0103 |
| Two-component system | 2020 | 28 | 24 | 0.320956 | -3.33419 | 0.367 | 0.369727 | 0.391228 | 1 | Inhibited | http://www.genome.jp/dbget-bin/show_pathway?hpy02020+HP_0082+HP_0099+HP_0103+HP_0714+HP_0815+HP_1032+HP_0115+HP_0601+HP_1442+HP_0690+HP_0693+HP_0512+HP_1529+HP_1067+HP_0019+HP_0393+HP_0616+HP_0392+HP_0144+HP_0145+HP_0146+HP_0147+HP_1540+HP_1539 |
| Sulfur relay system | 4122 | 8 | 7 | 0.508121 | 1.844259 | 0.252 | 0.391228 | 0.391228 | 1 | Activated | http://www.genome.jp/dbget-bin/show_pathway?hpy04122+HP_0220+HP_0013+HP_1335+HP_0800+HP_0801+HP_0798+HP_0768 |

**Supplementary Table 4.**
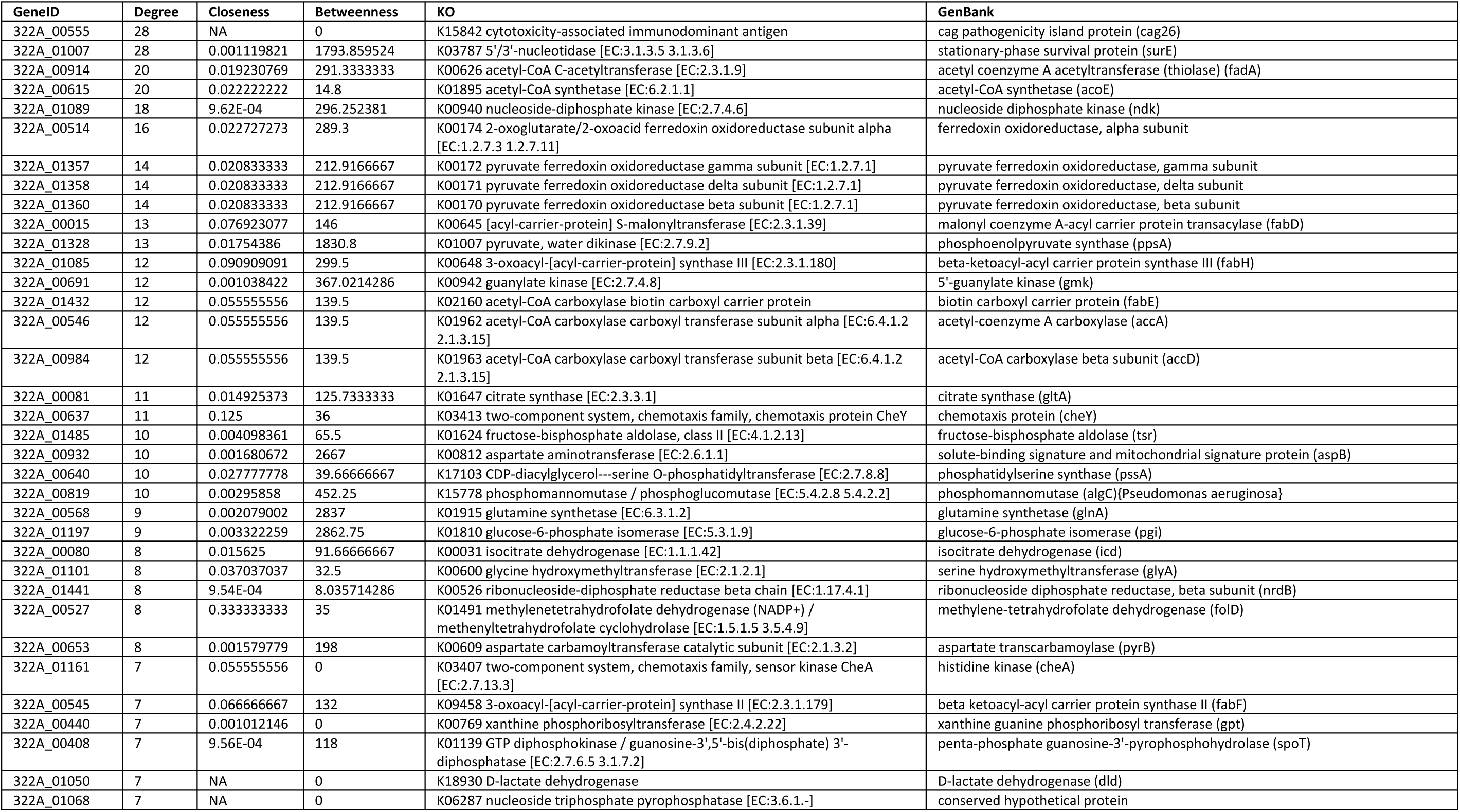
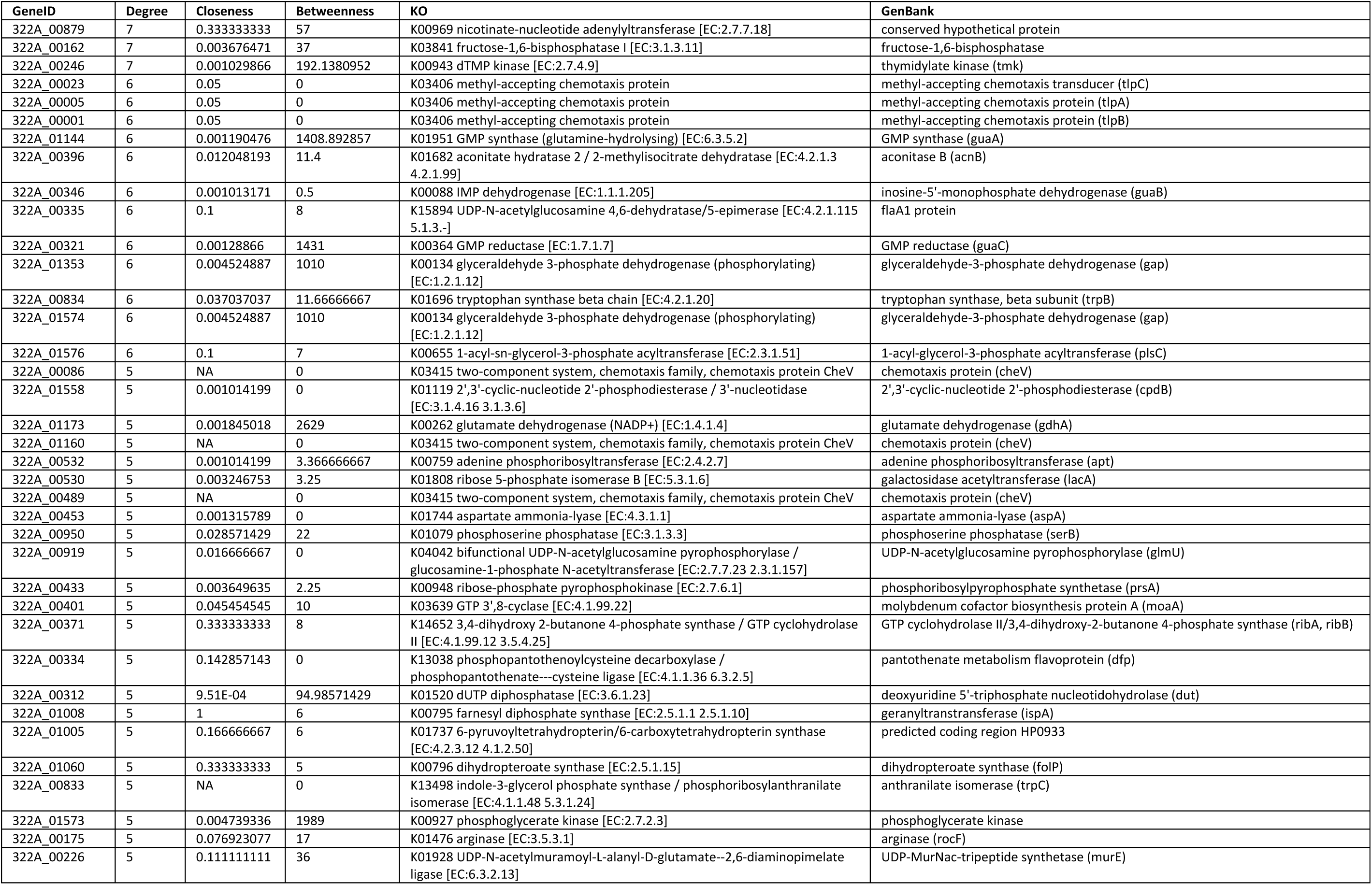
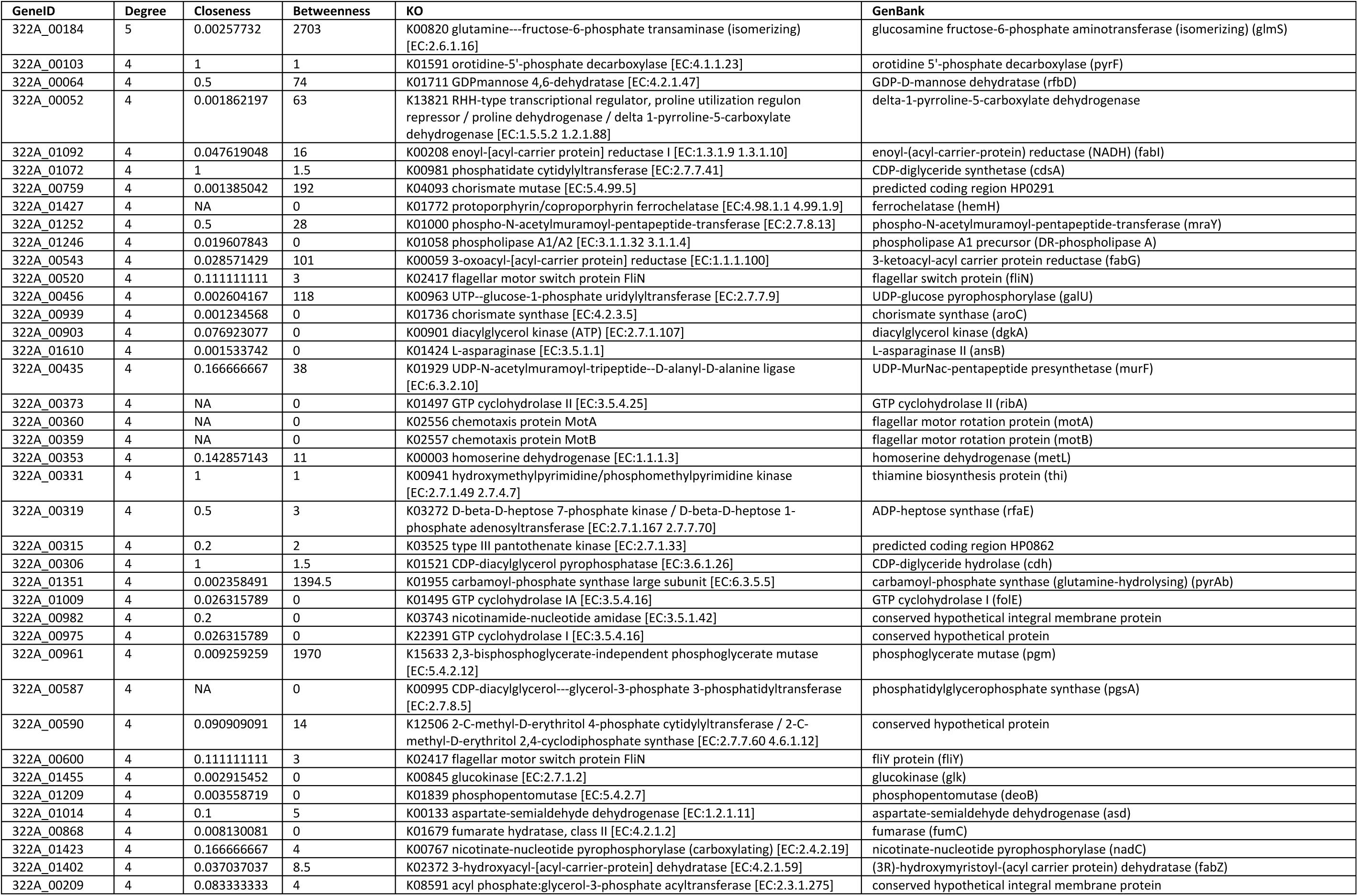
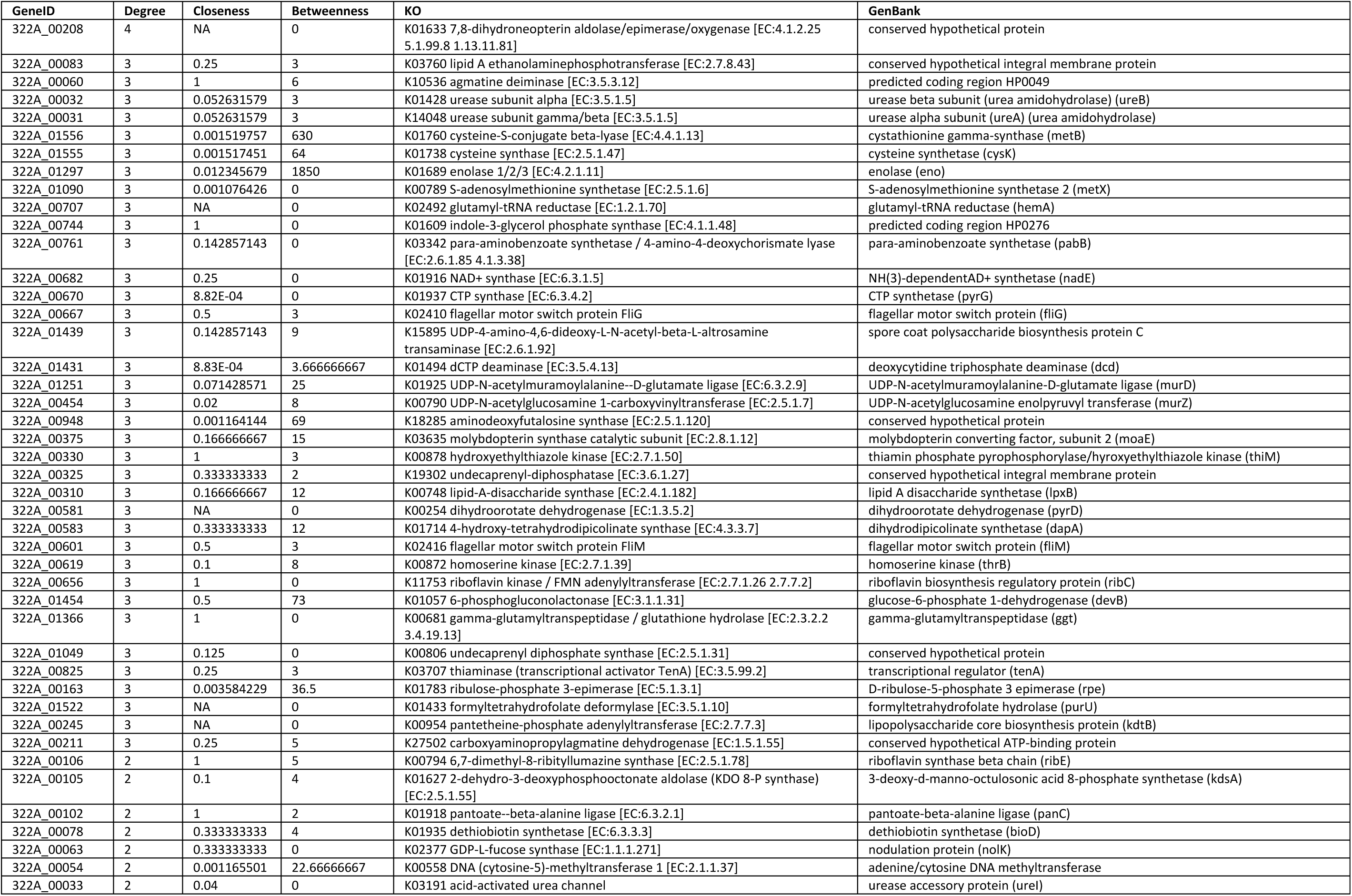
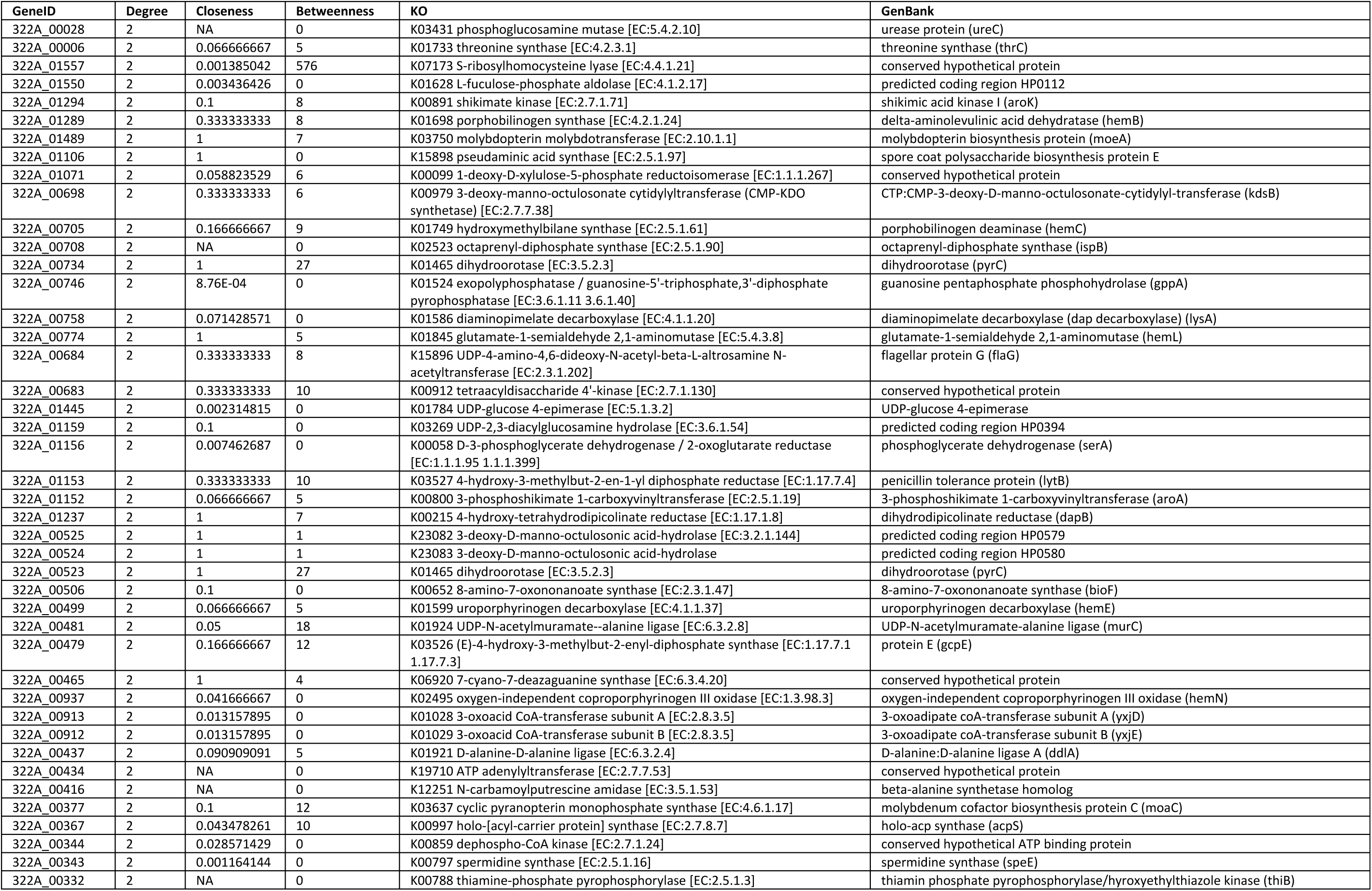
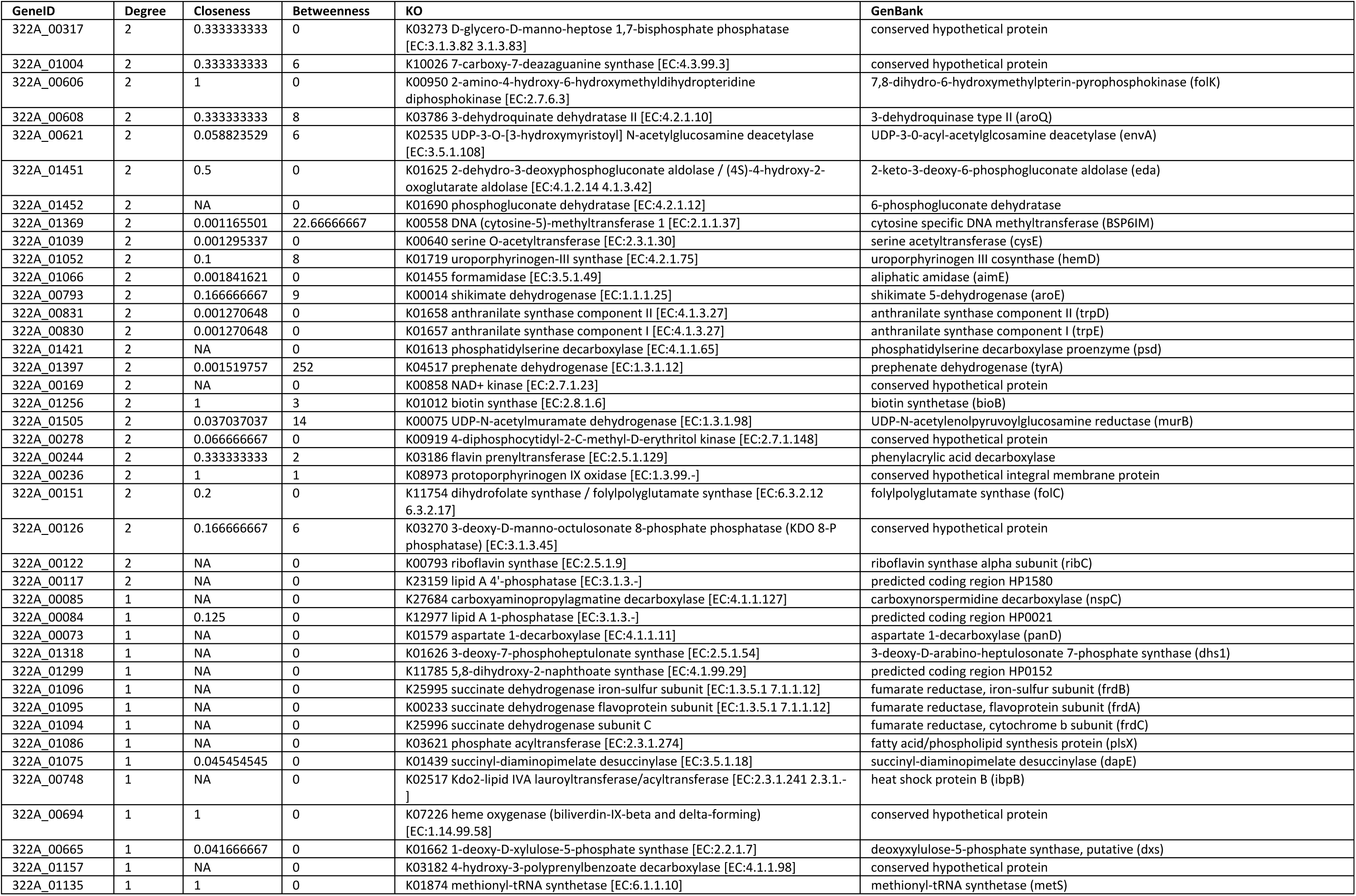
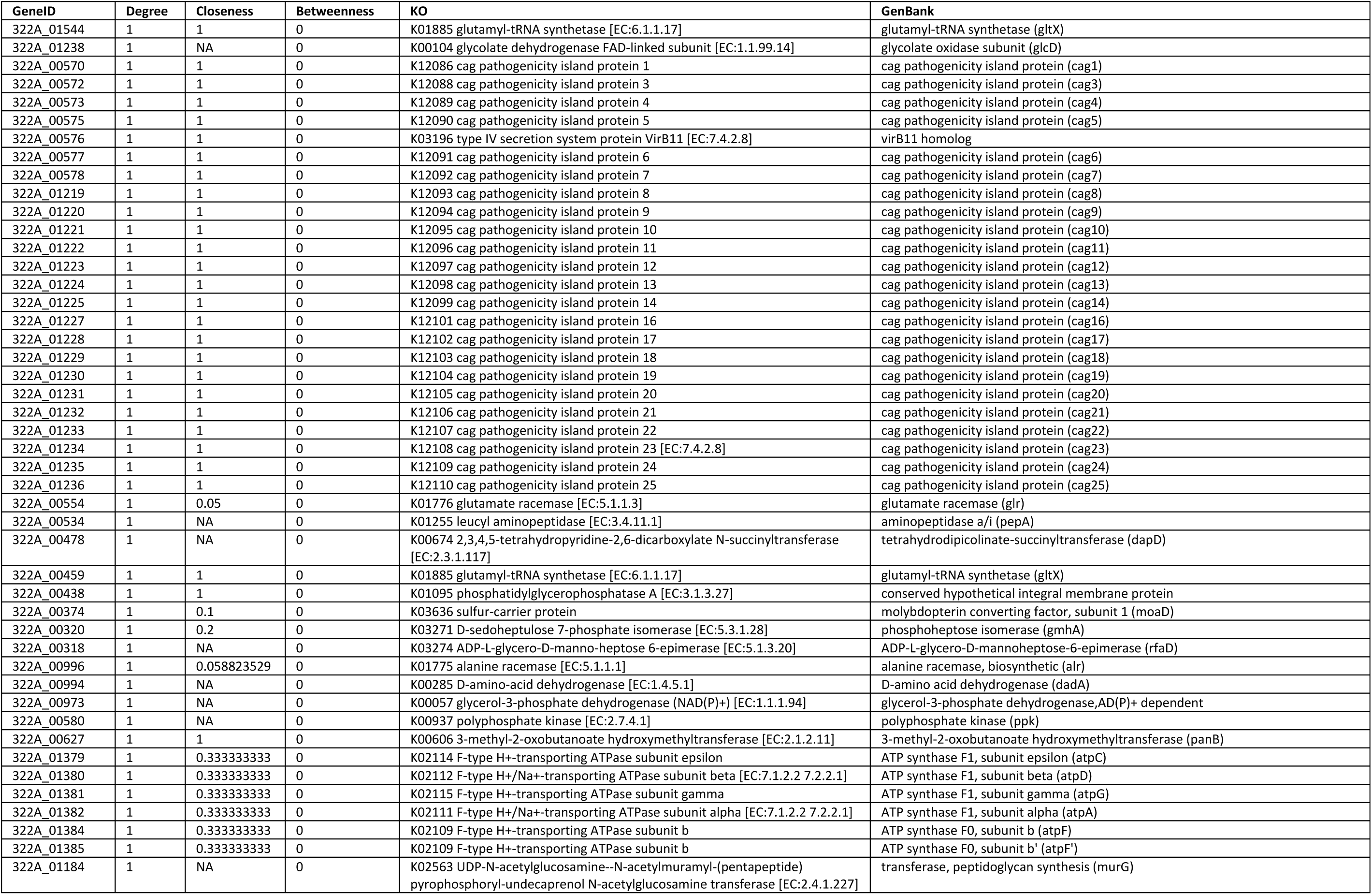
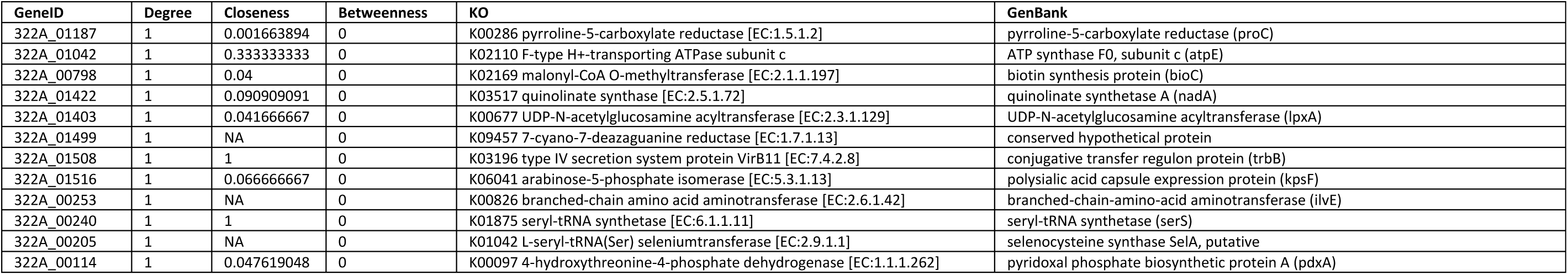
Network statistics calculated for the KEGG-based analysis of the significantly differentially expressed genes Only genes with a degree of 1 or more are shown. Genes with degree 0 are not connected to any other genes in the network.

**Supplementary Table 5.**
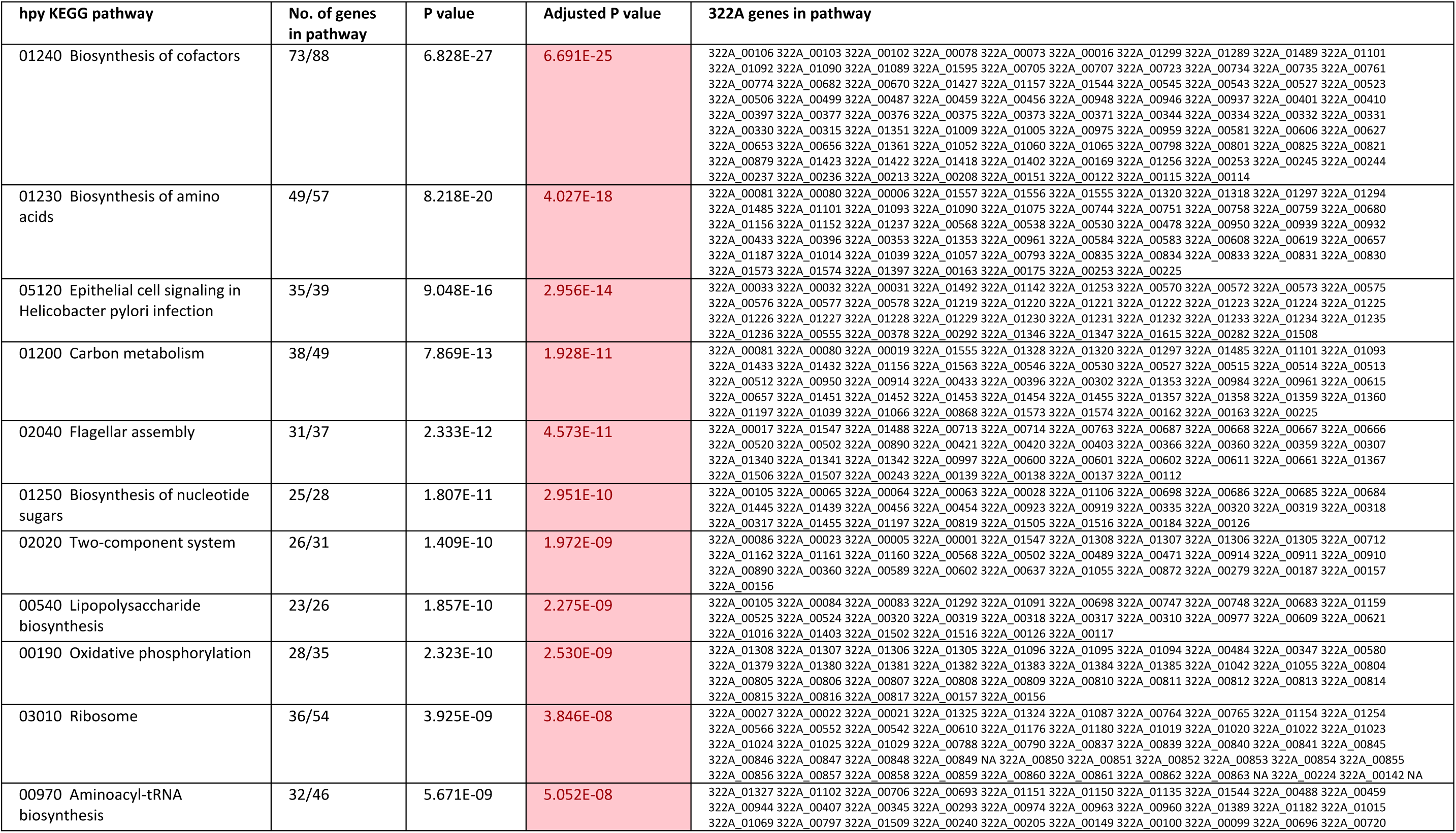
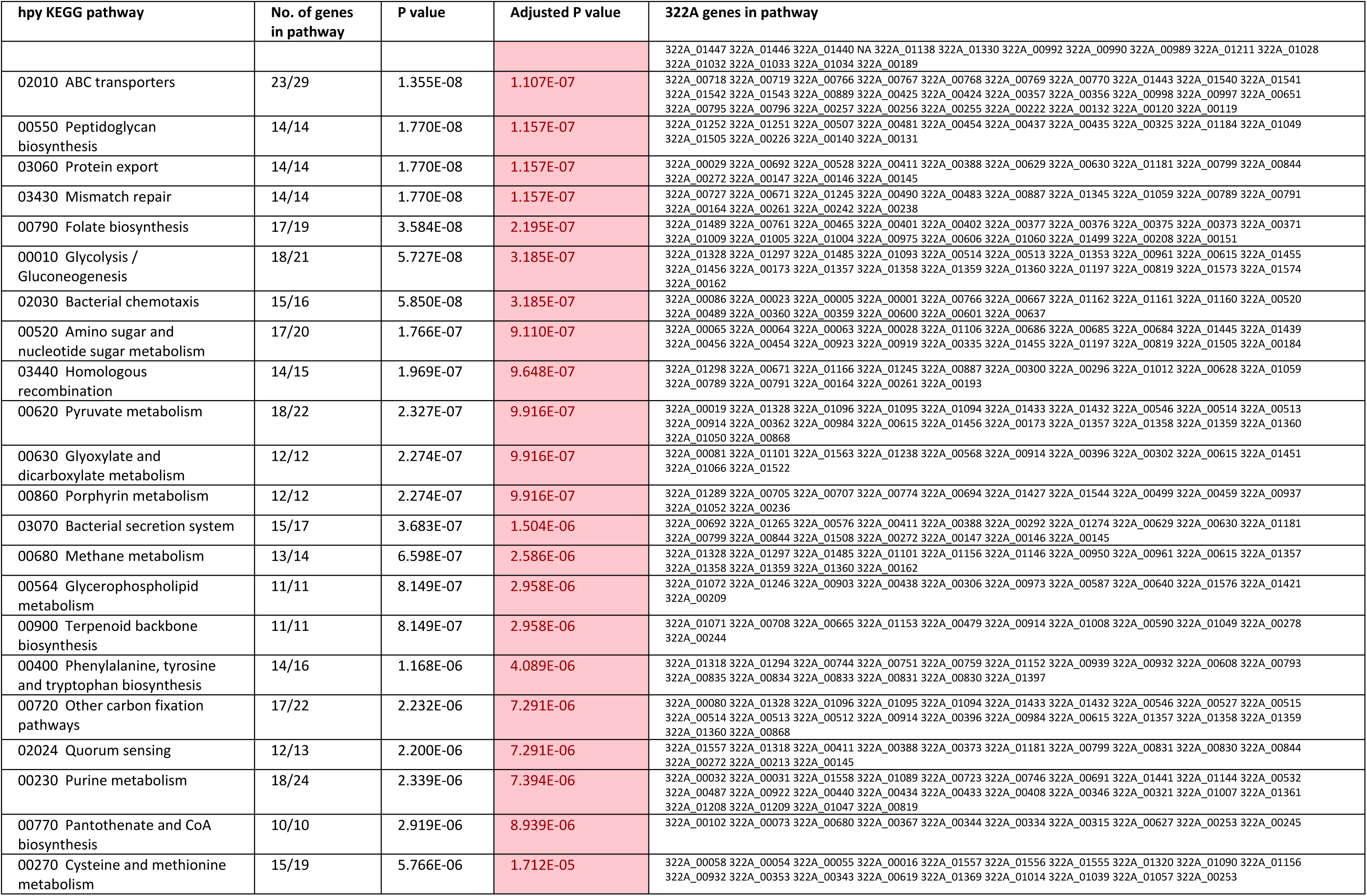
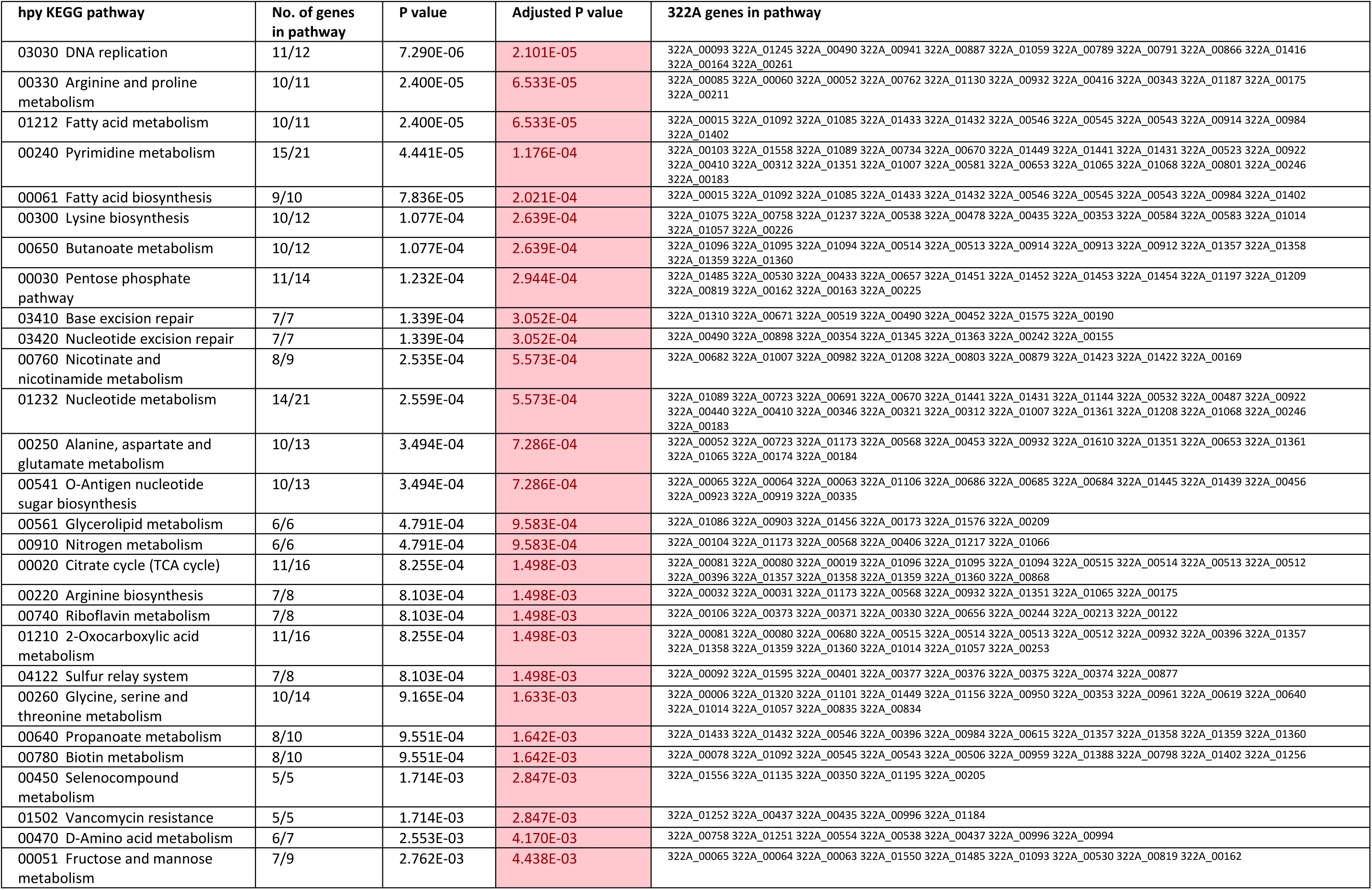
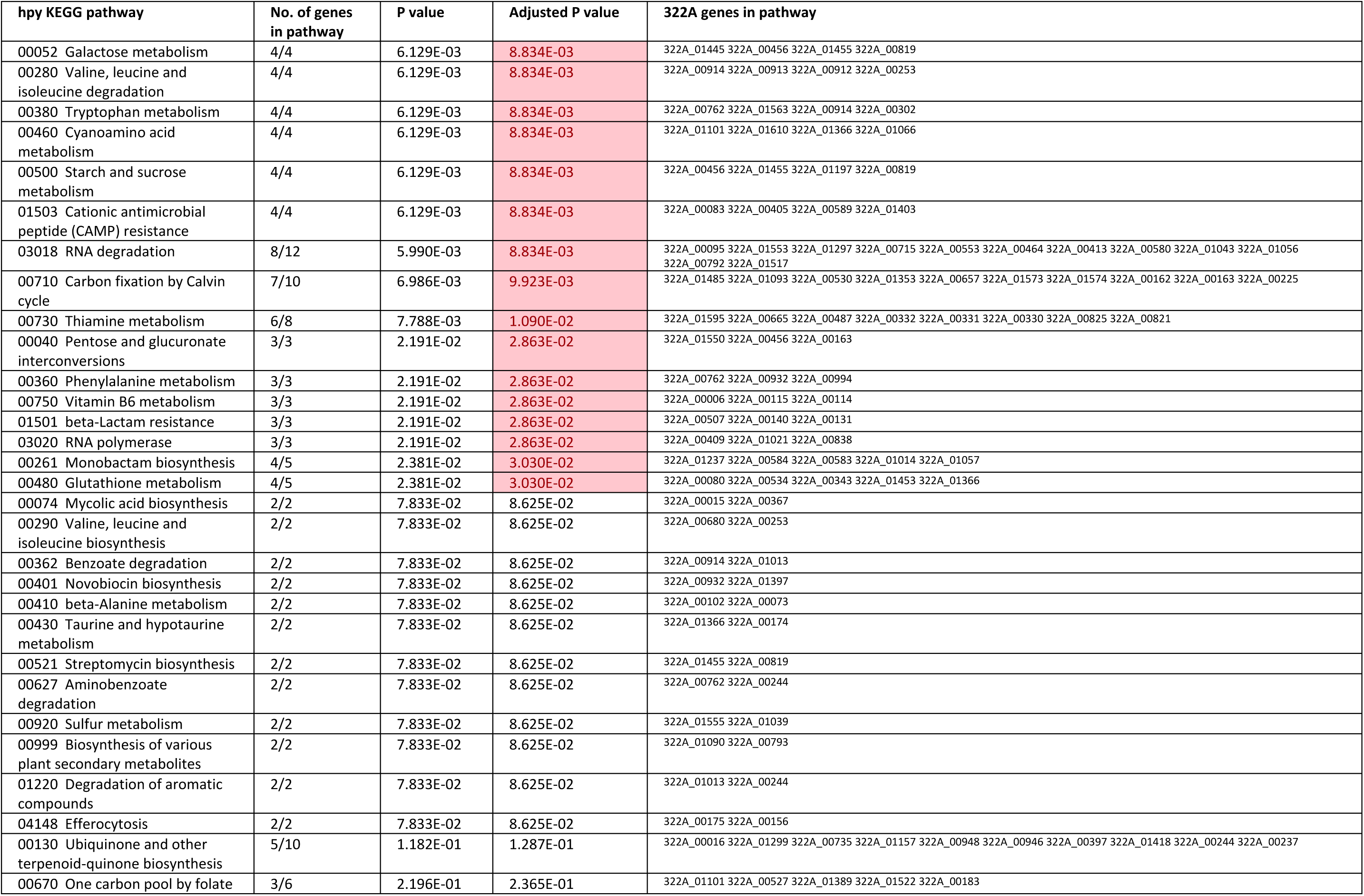
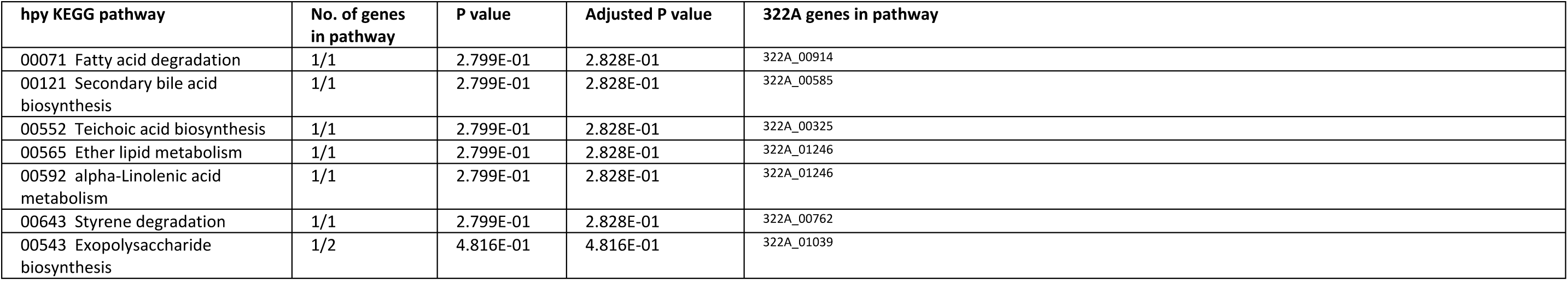
Over-representation analysis of significantly differentially expressed genes associated with KEGG metabolic pathways.

## References

1. Huang JQ, Zheng GF, Sumanac K, Irvine EJ, Hunt RH. Meta-analysis of the relationship between *cagA* seropositivity and gastric cancer. Gastroenterology. 2003 Dec;125(6):1636–44.

2. Rhead JL, Letley DP, Mohammadi M, Hussein N, Mohagheghi MA, Eshagh Hosseini M, et al. A new *Helicobacter pylori* vacuolating cytotoxin determinant, the intermediate region, is associated with gastric cancer. Gastroenterology. 2007 Sep;133(3):926–36.

3. Matos JI, De Sousa HAC, Marcos-Pinto R, Dinis-Ribeiro M. *Helicobacter pylori* CagA and VacA genotypes and gastric phenotype: a meta-analysis. Eur J Gastroenterol Hepatol. 2013 Dec;25(12):1431–41.

4. Garvey E, Rhead J, Suffian S, Whiley D, Mahmood F, Bakshi N, et al. High incidence of antibiotic resistance amongst isolates of *Helicobacter pylori* collected in Nottingham, UK, between 2001 and 2018. J Med Microbiol [Internet]. 2023 Nov 14 [cited 2024 Aug 22];72(11). Available from: https://www.microbiologyresearch.org/content/journal/jmm/10.1099/jmm.0.001776

5. EFSA Panel on Additives and Products or Substances used in Animal Feed (FEEDAP). Scientific Opinion on the safety and efficacy of vitamin K3 (menadione sodium bisulphite and menadione nicotinamide bisulphite) as a feed additive for all animal species. EFSA J [Internet]. 2014 Jan [cited 2024 Sep 3];12(1). Available from: https://data.europa.eu/doi/10.2903/j.efsa.2014.3532

6. Lamson DW, Plaza SM. The anticancer effects of vitamin K. Altern Med Rev J Clin Ther. 2003 Aug;8(3):303–18.

7. Sun JS, Tsuang YH, Huang WC, Chen LT, Hang YS, Lu FJ. Menadione-induced cytotoxicity to rat osteoblasts. Cell Mol Life Sci. 1997;53(12):967.

8. Suresh S, Raghu D, Karunagaran D. Menadione (Vitamin K3) Induces apoptosis of human oral cancer cells and reduces their metastatic potential by modulating the expression of epithelial to mesenchymal transition markers and inhibiting migration. Asian Pac J Cancer Prev. 2013 Sep 30;14(9):5461–5.

9. Marchionatti AM, Picotto G, Narvaez CJ, Welsh J, Tolosa De Talamoni NG. Antiproliferative action of menadione and 1,25(OH)2D3 on breast cancer cells. J Steroid Biochem Mol Biol. 2009 Feb;113(3–5):227–32.

10. Oztopcu-Vatan P, Sayitoglu M, Gunindi M, Inan E. Cytotoxic and apoptotic effects of menadione on rat hepatocellular carcinoma cells. Cytotechnology. 2015 Dec;67(6):1003–9.

11. Schlievert PM, Merriman JA, Salgado-Pabón W, Mueller EA, Spaulding AR, Vu BG, et al. Menaquinone analogs inhibit growth of bacterial pathogens. Antimicrob Agents Chemother. 2013 Nov;57(11):5432–7.

12. Andrade JC, Morais Braga MFB, Guedes GMM, Tintino SR, Freitas MA, Quintans LJ, et al. Menadione (vitamin K) enhances the antibiotic activity of drugs by cell membrane permeabilization mechanism. Saudi J Biol Sci. 2017 Jan;24(1):59–64.

13. Park BS, Lee HK, Lee SE, Piao XL, Takeoka GR, Wong RY, et al. Antibacterial activity of *Tabebuia impetiginosa* Martius ex DC (Taheebo) against *Helicobacter pylori*. J Ethnopharmacol. 2006 Apr;105(1–2):255–62.

14. Lee MH, Yang JY, Cho Y, Woo HJ, Kwon HJ, Kim DH, et al. Inhibitory effects of menadione on *Helicobacter pylori* growth and *Helicobacter pylori*-induced inflammation via NF-κB inhibition. Int J Mol Sci. 2019 Mar 7;20(5):1169.

15. Marcus EA, Sachs G, Scott DR. Acid-regulated gene expression of *Helicobacter pylori* : Insight into acid protection and gastric colonization. Helicobacter. 2018 Jun;23(3):e12490.

16. Loh JT, Beckett AC, Scholz MB, Cover TL. High-salt conditions alter transcription of *Helicobacter pylori* genes encoding outer membrane proteins. Young VB, editor. Infect Immun. 2018 Mar;86(3):e00626–17.

17. Miles AA, Misra SS, Irwin JO. The estimation of the bactericidal power of the blood. Epidemiol Infect. 1938 Nov;38(6):732–49.

18. Afgan E, Baker D, Batut B, van den Beek M, Bouvier D, Čech M, et al. The Galaxy platform for accessible, reproducible and collaborative biomedical analyses: 2018 update. Nucleic Acids Res. 2018 Jul 2;46(W1):W537–44.

19. Wilkinson DJ, Dickins B, Robinson K, Winter JA. Genomic diversity of *Helicobacter pylori* populations from different regions of the human stomach. Gut Microbes. 2022 Dec 31;14(1):2152306.

20. Wingett S, Andrews S. FastQ Screen: A tool for multi-genome mapping and quality control. F1000Research. 2018;24(7):1338.

21. Kim D, Paggi JM, Park C, Bennett C, Salzberg SL. Graph-based genome alignment and genotyping with HISAT2 and HISAT-genotype. Nat Biotechnol. 2019 Aug;37(8):907–15.

22. Liao Y, Smyth GK, Shi W. featureCounts: an efficient general purpose program for assigning sequence reads to genomic features. Bioinformatics. 2014 Apr 1;30(7):923–30.

23. Love MI, Huber W, Anders S. Moderated estimation of fold change and dispersion for RNA-seq data with DESeq2. Genome Biol. 2014 Dec;15(12):550.

24. Tarca AL, Draghici S, Khatri P, Hassan SS, Mittal P, Kim JS, et al. A novel signaling pathway impact analysis. Bioinforma Oxf Engl. 2009 Jan 1;25(1):75–82.

25. Zhang JD, Wiemann S. KEGGgraph: a graph approach to KEGG PATHWAY in R and bioconductor. Bioinforma Oxf Engl. 2009 Jun 1;25(11):1470–1.

26. Csardi G, Nepusz T. The igraph software package for complex network research. InterJournal. 2006;Complex Systems:1695.

27. Gustavsen JA, Pai S, Isserlin R, Demchak B, Pico AR. RCy3: Network biology using Cytoscape from within R. F1000Research. 2019;8:1774.

28. Shannon P, Markiel A, Ozier O, Baliga NS, Wang JT, Ramage D, et al. Cytoscape: a software environment for integrated models of biomolecular interaction networks. Genome Res. 2003 Nov;13(11):2498–504.

29. Hoyles L, Pontifex MG, Rodriguez-Ramiro I, Anis-Alavi MA, Jelane KS, Snelling T, et al. Regulation of blood-brain barrier integrity by microbiome-associated methylamines and cognition by trimethylamine *N*-oxide. Microbiome. 2021 Nov 27;9(1):235.

30. Francke C, Groot Kormelink T, Hagemeijer Y, Overmars L, Sluijter V, Moezelaar R, et al. Comparative analyses imply that the enigmatic sigma factor 54 is a central controller of the bacterial exterior. BMC Genomics. 2011 Dec;12(1):385.

31. Inatsu S, Ohsaki A, Nagata K. Idebenone acts against growth of *Helicobacter pylori* by inhibiting its respiration. Antimicrob Agents Chemother. 2006 Jun;50(6):2237–9.

32. Gupta A, Imlay JA. How a natural antibiotic uses oxidative stress to kill oxidant-resistant bacteria. Proc Natl Acad Sci. 2023 Dec 26;120(52):e2312110120.

33. Jahn LJ, Munck C, Ellabaan MMH, Sommer MOA. Adaptive Laboratory evolution of antibiotic resistance using different selection regimes lead to similar phenotypes and genotypes. Front Microbiol. 2017 May 11;8:816.

34. Maeda T, Iwasawa J, Kotani H, Sakata N, Kawada M, Horinouchi T, et al. High-throughput laboratory evolution reveals evolutionary constraints in *Escherichia coli*. Nat Commun. 2020 Nov 24;11(1):5970.

35. Comtois SL. Role of the thioredoxin system and the thiol-peroxidases Tpx and Bcp in mediating resistance to oxidative and nitrosative stress in *Helicobacter pylori*. Microbiology. 2003 Jan 1;149(1):121–9.

36. Li C, Wu PY, Hsieh M. Growth-phase-dependent transcriptional regulation of the pcm and surE genes required for stationary-phase survival of *Escherichia coli*. Microbiol Read Engl. 1997 Nov;143 (Pt 11):3513–20.

37. Sun Y, Liu S, Li W, Shan Y, Li X, Lu X, et al. Proteomic analysis of the function of sigma factor σ54 in *Helicobacter pylori* survival with nutrition deficiency stress in vitro. PloS One. 2013;8(8):e72920.

38. Kazmierczak MJ, Wiedmann M, Boor KJ. Alternative sigma factors and their roles in bacterial virulence. Microbiol Mol Biol Rev MMBR. 2005 Dec;69(4):527–43.

39. Tarique KF, Abdul Rehman SA, Devi S, Tomar P, Gourinath S. Structural and functional insights into the stationary-phase survival protein SurE, an important virulence factor of *Brucella abortus*. Acta Crystallogr Sect F Struct Biol Commun. 2016 May;72(Pt 5):386–96.

40. Natrajan G, Noirot-Gros MF, Zawilak-Pawlik A, Kapp U, Terradot L. The structure of a DnaA/HobA complex from *Helicobacter pylori* provides insight into regulation of DNA replication in bacteria. Proc Natl Acad Sci. 2009 Dec 15;106(50):21115–20.

41. Slager J, Kjos M, Attaiech L, Veening JW. Antibiotic-induced replication stress triggers bacterial competence by increasing gene dosage near the origin. Cell. 2014 Apr;157(2):395–406.

42. Karnholz A, Hoefler C, Odenbreit S, Fischer W, Hofreuter D, Haas R. Functional and topological characterization of novel components of the *comB* DNA transformation competence system in *Helicobacter pylori*. J Bacteriol. 2006 Feb;188(3):882–93.

43. Hofreuter D, Odenbreit S, Haas R. Natural transformation competence in *Helicobacter pylori* is mediated by the basic components of a type IV secretion system. Mol Microbiol. 2001 Jul;41(2):379–91.

44. Engelmoer DJP, Rozen DE. Competence increases survival during stress in *Streptococcus pneumoniae*. Evolution. 2011;65(12):3475–85.

45. Kaźmierczak-Barańska J, Karwowski BT. Vitamin K contribution to DNA damage—advantage or disadvantage? A human health response. Nutrients. 2022 Oct 11;14(20):4219.

46. Seitz P, Blokesch M. Cues and regulatory pathways involved in natural competence and transformation in pathogenic and environmental Gram-negative bacteria. FEMS Microbiol Rev. 2013 May 1;37(3):336–63.

47. Dorer MS, Fero J, Salama NR. DNA damage triggers genetic exchange in *Helicobacter pylori*. PLoS Pathog. 2010 Jul 29;6(7):e1001026.

48. Pereira L, Hoover TR. Stable accumulation of sigma54 in *Helicobacter pylori* requires the novel protein HP0958. J Bacteriol. 2005 Jul;187(13):4463–9.

49. Doherty NC, Shen F, Halliday NM, Barrett DA, Hardie KR, Winzer K, et al. In *Helicobacter pylori*, LuxS is a key enzyme in cysteine provision through a reverse transsulfuration pathway. J Bacteriol. 2010 Mar;192(5):1184–92.

50. Lee WK, Ogura K, Loh JT, Cover TL, Berg DE. Quantitative effect of *luxS* gene inactivation on the fitness of *Helicobacter pylori*. Appl Environ Microbiol. 2006 Oct;72(10):6615–22.

51. Osaki T, Hanawa T, Manzoku T, Fukuda M, Kawakami H, Suzuki H, et al. Mutation of *luxS* affects motility and infectivity of *Helicobacter pylori* in gastric mucosa of a Mongolian gerbil model. J Med Microbiol. 2006 Nov 1;55(11):1477–85.

52. Loughlin MF, Barnard FM, Jenkins D, Sharples GJ, Jenks PJ. *Helicobacter pylori* mutants defective in RuvC Holliday junction resolvase display reduced macrophage survival and spontaneous clearance from the murine gastric mucosa. Infect Immun. 2003 Apr;71(4):2022–31.

53. Ha NC, Oh ST, Sung JY, Cha KA, Lee MH, Oh BH. Supramolecular assembly and acid resistance of *Helicobacter pylori* urease. Nat Struct Biol. 2001 Jun 1;8(6):505–9.

54. Hu LT, Foxall PA, Russell R, Mobley HL. Purification of recombinant *Helicobacter pylori* urease apoenzyme encoded by *ureA* and *ureB*. Infect Immun. 1992 Jul;60(7):2657–66.

55. Labigne A, Cussac V, Courcoux P. Shuttle cloning and nucleotide sequences of *Helicobacter pylori* genes responsible for urease activity. J Bacteriol. 1991 Mar;173(6):1920–31.

56. Weeks DL, Eskandari S, Scott DR, Sachs G. A H^+^-gated urea channel: the link between *Helicobacter pylori* urease and gastric colonization. Science. 2000 Jan 21;287(5452):482–5.

57. Skouloubris S, Thiberge JM, Labigne A, De Reuse H. The *Helicobacter pylori* UreI protein is not involved in urease activity but is essential for bacterial survival in vivo. Orndorff PE, editor. Infect Immun. 1998 Sep;66(9):4517–21.

58. Debowski AW, Walton SM, Chua EG, Tay ACY, Liao T, Lamichhane B, et al. *Helicobacter pylori* gene silencing in vivo demonstrates urease is essential for chronic infection. Dove SL, editor. PLOS Pathog. 2017 Jun 23;13(6):e1006464.

59. Kavermann H, Burns BP, Angermüller K, Odenbreit S, Fischer W, Melchers K, et al. Identification and characterization of *Helicobacter pylori* genes essential for gastric colonization. J Exp Med. 2003 Apr 7;197(7):813–22.

60. Lytton SD, Fischer W, Nagel W, Haas R, Beck FX. Production of ammonium by *Helicobacter pylori* mediates occludin processing and disruption of tight junctions in Caco-2 cells. Microbiology. 2005 Oct 1;151(10):3267–76.

61. Wroblewski LE, Shen L, Ogden S, Romero–Gallo J, Lapierre LA, Israel DA, et al. *Helicobacter pylori* dysregulation of gastric epithelial tight junctions by urease-mediated myosin ii activation. Gastroenterology. 2009 Jan;136(1):236–46.

62. Beswick EJ, Pinchuk IV, Minch K, Suarez G, Sierra JC, Yamaoka Y, et al. The *Helicobacter pylori* urease B subunit binds to CD74 on gastric epithelial cells and induces NF-κB activation and interleukin-8 production. Infect Immun. 2006 Feb;74(2):1148–55.

63. Harris P, Mobley H, Perez-Perez G, Blaser M, Smith P. *Helicobacter pylori* urease is a potent stimulus of mononuclear phagocyte activation and inflammatory cytokine production. Gastroenterology. 1996 Aug;111(2):419–25.

64. Censini S, Lange C, Xiang Z, Crabtree JE, Ghiara P, Borodovsky M, et al. *cag*, a pathogenicity island of *Helicobacter pylori*, encodes type I-specific and disease-associated virulence factors. Proc Natl Acad Sci. 1996 Dec 10;93(25):14648–53.

65. Ohnishi N, Yuasa H, Tanaka S, Sawa H, Miura M, Matsui A, et al. Transgenic expression of *Helicobacter pylori* CagA induces gastrointestinal and hematopoietic neoplasms in mouse. Proc Natl Acad Sci. 2008 Jan 22;105(3):1003–8.

66. Neal JT, Peterson TS, Kent ML, Guillemin K. *H. pylori* virulence factor CagA increases intestinal cell proliferation by Wnt pathway activation in a transgenic zebrafish model. Dis Model Mech. 2013 Jan 1;dmm.011163.

67. Robinson K, White J, Winter J. Differential inflammatory response to *Helicobacter pylori*infection: etiology and clinical outcomes. J Inflamm Res. 2015 Aug;137.

68. Foegeding N, Caston R, McClain M, Ohi M, Cover T. An overview of *Helicobacter pylori* VacA toxin biology. Toxins. 2016 Jun 3;8(6):173.

69. Osada S, Tomita H, Tanaka Y, Tokuyama Y, Tanaka H, Sakashita F, et al. The utility of vitamin K3 (menadione) against pancreatic cancer. Anticancer Res. 2008;28(1A):45–50.

70. Lee MH, Cho Y, Kim DH, Woo HJ, Yang JY, Kwon HJ, et al. Menadione induces G2/M arrest in gastric cancer cells by down-regulation of CDC25C and proteasome mediated degradation of CDK1 and cyclin B1. Am J Transl Res. 2016;8(12):5246–55.

71. Lee MH, Yang JY, Cho Y, Park M, Woo HJ, Kim HW, et al. Menadione induces apoptosis in a gastric cancer cell line mediated by down-regulation of X-linked inhibitor of apoptosis. Int J Clin Exp Med. 2016;9(2):2437–43.

72. Shahzad S, Ashraf MA, Sajid M, Shahzad A, Rafique A, Mahmood MS. Evaluation of synergistic antimicrobial effect of vitamins (A, B1, B2, B6, B12, C, D, E and K) with antibiotics against resistant bacterial strains. J Glob Antimicrob Resist. 2018 Jun;13:231–6.

73. Li LH, Lu HF, Liu YF, Lin YT, Yang TC. *fadACB* and *smeU1VWU2X* contribute to oxidative stress-mediated fluoroquinolone resistance in *Stenotrophomonas maltophilia*. Antimicrob Agents Chemother. 2022 Apr 19;66(4):e02043–21.

74. Blanco P, Corona F, Sánchez MB, Martínez JL. Vitamin K _3_ Induces the Expression of the *Stenotrophomonas maltophilia* SmeVWX multidrug efflux pump. Antimicrob Agents Chemother. 2017 May;61(5):e02453–16.

